# A *Legionella* toxin mimics tRNA and glycosylates the translation machinery to trigger a ribotoxic stress response

**DOI:** 10.1101/2022.06.10.495705

**Authors:** Advait Subramanian, Lan Wang, Tom Moss, Mark Voorhies, Smriti Sangwan, Erica Stevenson, Ernst H. Pulido, Samentha Kwok, Nevan J. Krogan, Danielle L. Swaney, Stephen N. Floor, Anita Sil, Peter Walter, Shaeri Mukherjee

**Author notes:** Authors contributed equally (Advait Subramanian, Lan Wang). Bay Area Institute of Science, Altos Labs, Redwood City, USA.

## Abstract

Pathogens often secrete proteins or nucleic acids that mimic the structure and/or function of molecules expressed in their hosts. Molecular mimicry empowers pathogens to subvert critical host processes and establish infection. We report that the intracellular bacterium *Legionella pneumophila* secretes the toxin SidI (substrate of icm/dot transporter I), which possesses a transfer RNA (tRNA)-like shape and functions as a mannosyl transferase. The 3.1 Å cryo-EM structure of SidI reveals an N-terminal domain that exhibits a characteristic ‘inverted L-shape’ and charge distribution that is present in two other known protein mimics of tRNAs, the bacterial elongation factor EF-G and the mammalian release factor eRF1. In addition, we show that SidI’s C-terminal domain adopts a glycosyl transferase B fold similar to a mannosyl transferase. This molecular coupling of the protein’s fold and enzymatic function allows SidI to bind and glycosylate components of the host translation apparatus, including the ribosome, resulting in a robust block of protein synthesis that is comparable in potency to ricin, one of the most powerful toxins known. Additionally, we find that translational pausing activated by SidI elicits a stress response signature reminiscent of the ribotoxic stress response that is activated by elongation inhibitors that induce ribosome collisions. SidI-mediated effects on the ribosome activate the stress kinases ZAKα and p38, which in turn drive an accumulation of the protein activating transcription factor 3 (ATF3). Intriguingly, ATF3 escapes the translation block imposed by SidI, translocates to the nucleus, and orchestrates the transcription of stress-inducible genes that promote cell death. Thus, using *Legionella* and its effectors as tools, we have unravelled the role of a ribosome-to-nuclear signalling pathway that regulates cell fate.

## Main

*Legionella pneumophila* (*L.p.*) is an intracellular bacterial pathogen that is the causative organism of an atypical form of pneumonia known as Legionnaires’ disease^1^. In an infected cell, *L.p.* secretes ∼300 bacterial protein effectors or toxins via its type IV secretion system (T4SS) into the host cell’s cytosol that manipulate several host cellular pathways in order to establish an endoplasmic reticulum (ER)-like replicative niche^2^. A vital process targeted by a subset of *L.p.* derived toxins is protein synthesis^3–11^, resulting in a disruption of the quality and quantity of proteins produced in an infected cell. Seven *L.p.* toxins have been proposed to have redundant functions in inhibiting the elongation step of protein synthesis^5,7,8, 10–12^. The secretion of these elongation inhibitor toxins (EITs) has been linked to the manipulation of diverse cellular outcomes that include ribosome stalling^5^, immune gene expression^4,5,8,13,14^, stress signaling^10,11,15–17^, cell cycle arrest^18^, and cell death^10,19^. Amongst these toxins, however, only the *Legionella* glucosyl transferase family of proteins (Lgt 1-3) and the *L.p.* kinase LegK4 have defined mechanistic targets^7,11,12^, while the roles of SidI, SidL and RavX remain unknown. Moreover, the relative contributions of each of these toxins towards protein synthesis inhibition and the resulting consequences of such perturbations to host cell physiology are poorly understood.

### SidI potently inhibits protein synthesis and binds to ribosomes

To address these gaps in our knowledge, we first undertook a reductionist approach by purifying N-terminally GST-tagged versions of the *L.p.* EITs, namely Lgt2, SidI, SidL, LegK4 Δ1-58 (labelled as LegK4) and RavX after recombinantly expressing them in *Escherichia coli* (*E.coli*) (Figure S1a). We then titrated these toxins into cell free translation extracts isolated from rabbit reticulocytes, covering a concentration range of 1 picomolar to 10 micromolar, and tested their relative effectiveness to inhibit the translation of a reporter luciferase mRNA into functional luciferase. The incubation of GST-SidI with the translation extract elicited strong inhibition of protein synthesis (Figure 1a; red line) with a measured half-maximal inhibitory concentration (IC_50_) of ∼900 picomolar, which is ∼100-fold to ∼1000-fold more potent than the other tested *L.p.* toxins (Figure 1b), and comparable to the potency of the toxin ricin^20^, one of the most powerful toxins known. In line with these results, the transient expression of SidI fused to a N-terminal FLAG-tag (FLAG-SidI) in HEK293T cells also induced a strong inhibition (∼90%) of protein synthesis rates as measured by puromycin pulse/chase assays (Figure S1b). The results suggest that SidI targets important host factor(s) to inhibit translation and argue against a mechanism shared between SidI and the other tested EITs, as has been proposed previously^3,10,21^.

**Figure 1.**
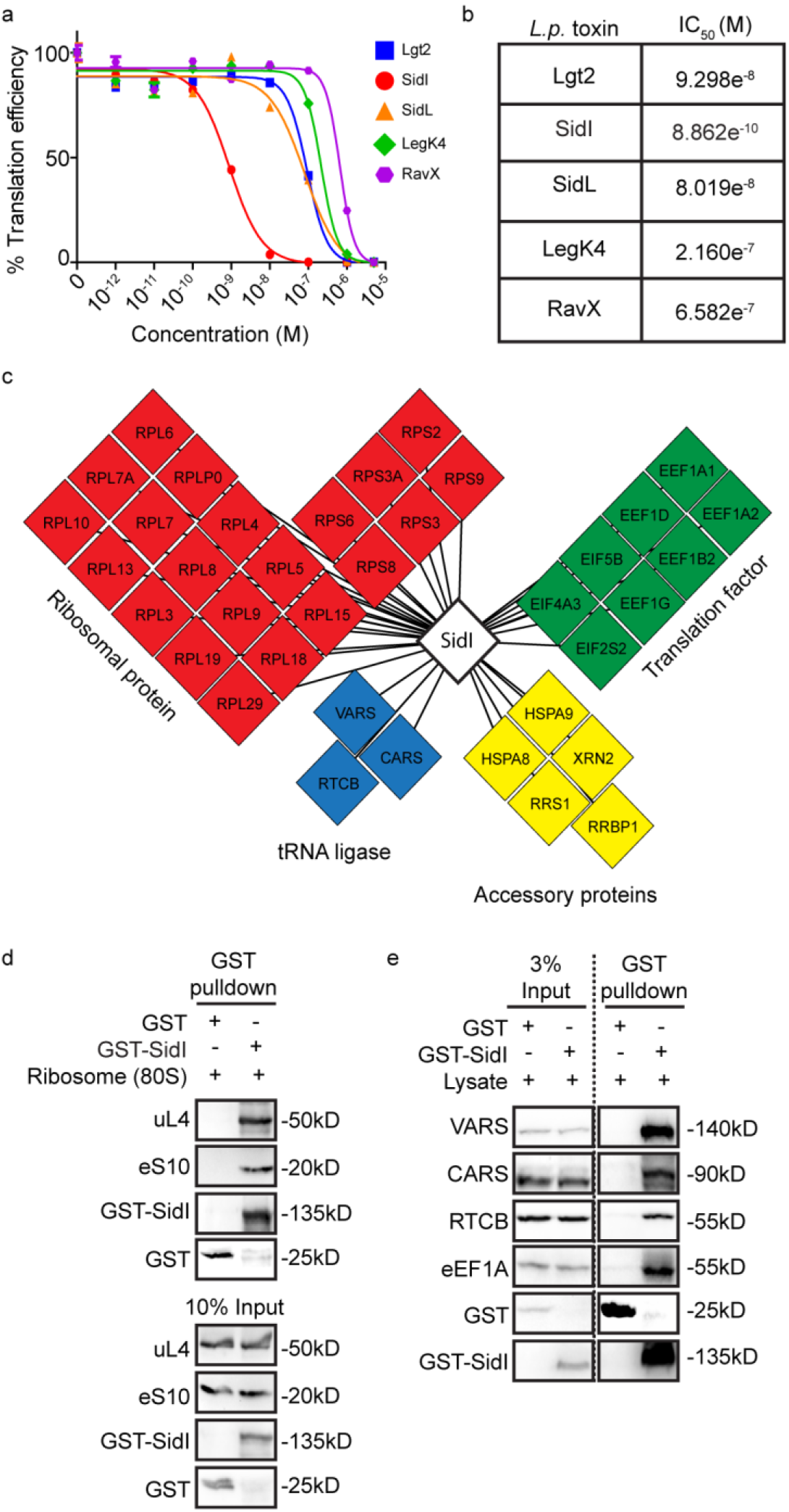
Sid potently inhibits protein synthesis and binds to the cellular translation machinery. (a) Cell free translation of luciferase mRNA in rabbit reticulocyte lysates (RRLs) incubated with purified GST-tagged *L.p.* toxins. Graph depicts the %translation efficiency of luciferase upon incubation with log-fold concentrations of *L.p.* toxins (Data represent mean ± standard error of the mean; n = 3 replicates per treatment condition). (b) IC_50_ values of *L.p.* toxins calculated by non-linear regression analysis. (c) Mass spectrometry analysis of SidI interacting partners from HEK293T and rabbit reticulocyte cell lysates. Cellular translation machinery components enriched by GST-SidI pulldown in atleast 2 out of 3 experiments are depicted. (d) GST-SidI puldown of purified monosomes. Immunoblotting was performed using antibodies against 60S (uL4/RPL4) and 40S (eS10/RPS10) ribosomal proteins and GST. Data are representative of 3 independent experiments. (e) Immunoblot of tRNA interacting proteins VARS, CARS, RTCB and eEF1A precipiated by GST-SidI from HEK293T lysates. Data are representative of 3 independent experiments.

To determine the putative target(s) of SidI in mammalian cells, we undertook an unbiased proteomics approach in which we incubated either GST or GST-SidI separately with rabbit reticulocyte lysates (RRLs) or HEK293T lysates and identified the interacting partners in each of these cell extracts by affinity purification with glutathione-coupled Sepharose beads followed by mass spectrometry (AP-MS). After peptide intensities in the eluted pulldown fractions were calculated, we first curated the datasets by removing any peptides from proteins that are common contaminants in AP-MS procedures and peptides that were enriched in the GST-pulldown eluates (GST-SidI/GST intensity ratio <1). Further analyses on the remaining peptides revealed a conserved set of 81 interacting partners that were either uniquely detected or enriched after GST-SidI pulldowns in at least two out of the three AP-MS experiments (Figure S1c and Table S1). Forty-five percent of these interactors (37/81) were *bona fide* components of the cellular protein synthesis apparatus and included multiple ribosomal proteins, elongation factors and tRNA ligases (Figure 1c and Figure S1d).

Because of the enrichment of both large and small ribosomal subunit proteins in our SidI pulldown eluates (Figure S1d), we wondered if SidI might bind to ribosomes. Indeed, affinity tag pulldown and immunoblotting experiments determined that SidI directly interacted with individual ribosomes (80S ribosome fraction) purified from cell extracts (Figure 1d). SidI also precipitated elongation factor eEF1A and a repertoire of tRNA ligases (valine tRNA ligase VARS, cysteine tRNA ligase CARS and the tRNA splicing ligase RTCB) from HEK293T cell extracts (Figure 1e). The results validate the quality of our AP-MS-determined interactome, as well as previously known interaction partners of SidI reported in other studies^10,22^. These initial findings focused our efforts to understand the mechanism of action of this *L.p.* EIT.

### Structural characterization of SidI by cryo-electron microscopy reveals molecular mimicry

To gain further mechanistic insights into the function of SidI, we next determined the structure of SidI using single particle cryo-electron microscopy (cryo-EM). To this end we subjected purified full-length SidI (110kD) with an N-terminal GST tag to cryo-EM imaging. Multiples rounds of two- and three-dimensional (3D) classifications of 749,000 particles analyzed resulted in a consensus structure, that upon refinement, yielded a map with an average resolution of 3.1 Å, approaching 2.5 Å in the core, with most side chain density well-resolved (Figure 2a, Figure S2, Table S2). A structure prediction analysis^23^ using the primary sequence of SidI showed that the C-terminal part of SidI (amino acids 355-865) shared significant homology to a few glycosyl transferase (GT) enzymes, as has been suggested previously^22^. These enzymes utilize nucleotide-linked monosaccharides as precursors to catalyze the transfer of monosaccharide moieties to acceptor substrates^24^. We further divided the C-terminal domain into two subdomains (domains B and C) based on their positions relative to the predicted nucleotide sugar binding pocket (Figures 2a, b, m; blue and yellow colored; see below). Structure prediction analyses did not find any proteins in the protein database that shared significant homology with the N-terminal part of SidI (amino acids 136-343). We therefore built an atomic model into the EM map of SidI starting from the C-terminal domain using the crystal structure of the *B. subtilis* GT BshA as a template (PDB: 5D00)^25^ and built the N-terminal domain *de novo*. We did not observe any additional density that was not explained by the primary sequence of SidI. We also observed no density representing the GST-tag, as it was likely averaged out during reconstruction due to a flexible linkage between GST and SidI.

**Figure 2.**
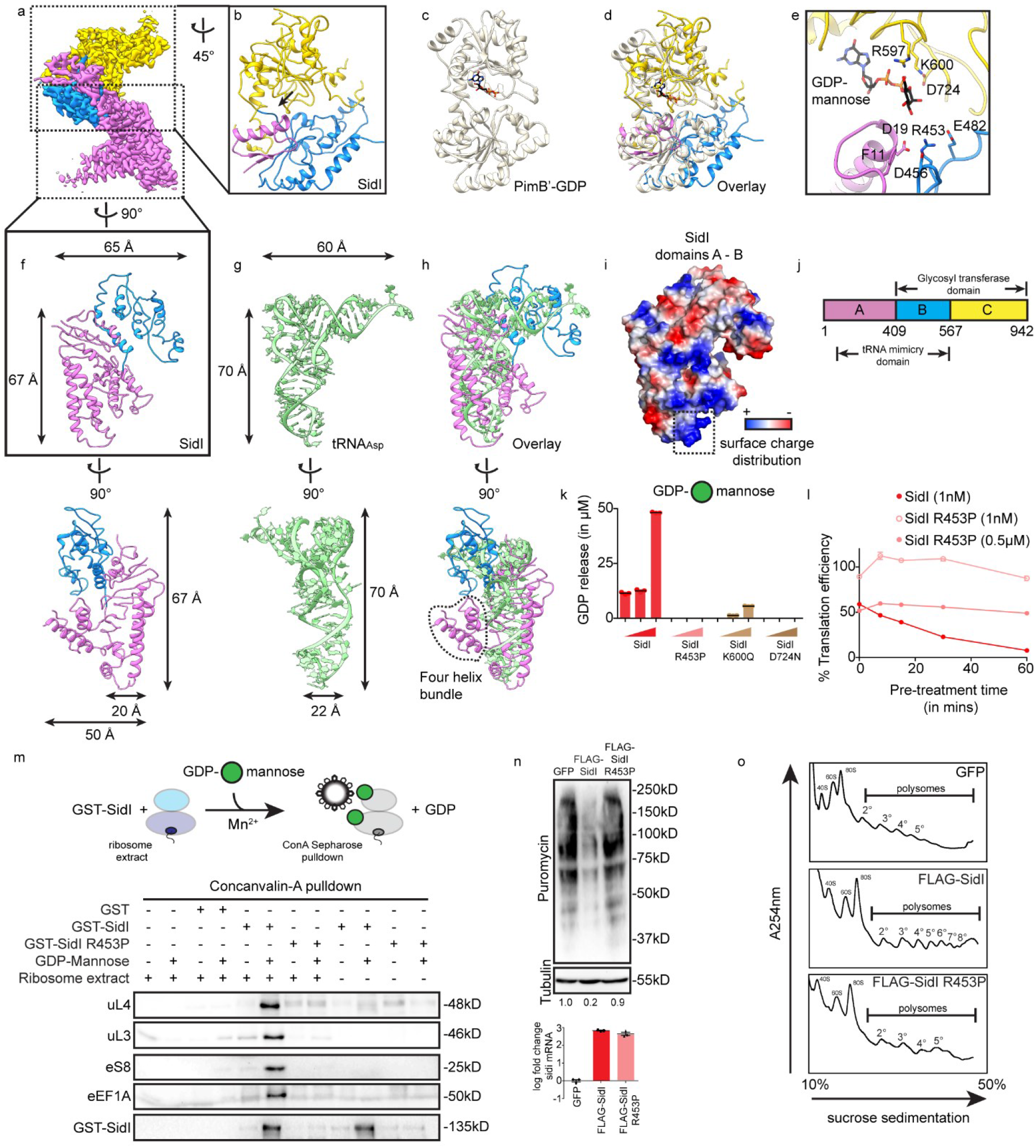
SidI couples a tRNA-like shape and glycosyl transferase function to induce ribosome stalling. (a) Cryo-EM map of SidI with individual domains coloured separately. (b) Rotate view of panel (a) showing the glycosyl transferase domain. Arrow depicts the putative nucleotide-sugar binding cleft. (c) Structure of the mannosyl-transferase PimB’ from *Corynebacterium glutamicum.* bound to GDP in the enzymatic pocket (PDB:3OKA). (d) Overlay of (b) and (c) showing the structural similarity between SidI and GDP-PimB’. (e) Enzymatic pocket of SidI with GDP-mannose docked. Amino acids potentially forming direct contact with GDP-mannose or stabilizing the fold of the enzymatic pocket are shown in sticks. (f – h) Structures of the tRNA mimicry domain of SidI in comparison with tRNA_Asp_ with length and width calculated. (i) Surface charge distribution of the tRNA mimicry domain of SidI (red represents negative charges and blue positive charged). Box highlights a positively charged loop conserved in other tRNA mimicry proteins. (j) Schematic representation of the domain architecture of SidI. Domain A of SidI is coloured magenta, domain B in blue and domain C in gold. PimB’ is coloured in light grey. tRNA_Asp_ is coloured in green. In all stick representations, atoms are coloured using the CPK scheme. (k) GDP hydrolysis assay with GST-SidI or enzymatic pocket mutants (R453P; K600Q, D724N) at concentrations of 0.5µM, 1µM and 2µM. 50µM of GDP-mannose was povided as a precursor. Histograms depict amount of GDP release in µM (Data represent mean ± standard error of the mean; n = 3 replicates per treatment condition). (l) Cell free luciferase translation in RRLs pretreated with GST-SidI or GST-SidI R453P at indicated concentrations. Graph depicts the %translation efficiency of luciferase with different pre-treatment times (Data represent mean ± standard error of the mean; n = 3 replicates per time point). (m) Concanavalin-A mediated enrichment of proteins in 30nM ribosome extracts after *in vitro* mannoylation reactions with 1.4µM of GST, GST-SidI or GST-SidI R453P. Immunoblotting was performed using antibodies against uL4 (RPL4), uL3 (RPL3), eS8 (RPS8), eEF1A and GST. Data are representative of 2 independent experiments. (n) Cells expressing GFP, FLAG-SidI or FLAG-SidI R453P subjected to puromycin pulse-chase analyses. Immunoblotting of lysates was performed using antibodies against puromycin and Tubulin. Data are representative of 2 independent experiments. Values below the immunoblots in (n) depict densitometric quantification of puromycylated peptides normalized to the amount of Tubulin in cells. Expression levels of SidI and SidI R453P were determined by qRT-PCR (Data represent mean ± standard error of the mean; n = 3). Data were normalized to the average Ct values of the reference gene ribosomal protein S29 (*RPS29*). (o) Post nuclear supernatants of cells expressing GFP, FLAG-SidI or FLAG-SidI R453P subjected to ultracentrifugation in 10% - 50% sucrose gradients followed by polysome profiling. The 40S, 60S and 80S peaks are marked. Progressive polysome peaks mark number of ribosomes per mRNA (2°-disome; 3°-trisome etc.). Plots are representative of 3 independent experiments.

### SidI adopts a glycosyl transferase B fold

Structurally, GT enzymes possess one of three folds in their catalytic domains^26^. The GT-A fold contains only one Rossman-like domain; the GT-B fold contains two individual Rossman-like domains; and the GT-C fold presents a complex architecture containing multiple transmembrane α-helices^26^. The 3D model of SidI shows that the C-terminal domain possesses two subdomains (B and C) with Rossman-like folds that are connected by a flexible hinge region (Figure 2b), with a putative nucleotide-sugar binding pocket located within a cleft between the two subdomains (Figure 2b, arrow). This domain architecture is highly conserved in a group of GTs that adopt the GT-B fold^26,27^.

To understand if the GT-B fold adopted by SidI has functional significance, we tested the ability of GST-SidI to catalyze the hydrolysis of nucleotides from UDP-glucose, GDP-mannose or GDP-fucose, precursors commonly found in cells, by measuring the release of the nucleotide, a by-product of the glycosyl transferase reaction^24^. As a control we also incubated these precursors with GST-Lgt2, an *L.p.* EIT that adopts the GT-A fold and utilizes UDP-glucose as a precursor^7,22,28,29^. Upon measuring nucleotide release, we determined that SidI hydrolyzed GDP from GDP-mannose most efficiently, while the hydrolysis of GDP from GDP-fucose was less efficient (Figure S3a). The hydrolysis of UDP from UDP-glucose was undetectable under our assay conditions, in accordance with previous studies^22^. By contrast, Lgt2 only catalyzed the release of UDP from UDP-glucose (Figure S3a).

We, hence, compared the structure of domains B and C to that of a known mannosyl transferase, the *C. glutamicum* enzyme PimB’ (PDB ID: 3OKA)^30^. Indeed, an overlay of PimB’ (Figure 2c) to SidI showed a close matching between their secondary structures (Figure 2d), with a conservation of amino acids that form the catalytic pocket in both proteins (Figure 2e and S3b). Notably, the first fifty amino acids of domain A folds on top of domains B and C and together they constitute the GT domain of SidI (Figures 2b and 2j).

### SidI possesses a tRNA-mimicry domain

Visualization and analyses of the 3D map suggested that an important feature of SidI’s domain architecture may lie in three characteristic distinct kinks in its overall structure (Figure S3c). One kink is near 90 degrees and is located in the GT domain (Figure S3c; panel 1). Another kink of approximately 60 degrees lies in domain A that connects this domain to a four-helix bundle (Figure S3c; panel 3). A third kink revealed another striking 90-degree angle that connects domains A and B. Together the domain architecture confers a characteristic ‘inverted L-shape’ (Figure 2f and Figure S3c, panel 2), resembling the shape of a tRNA. Similar shapes are found in a unique class of proteins and nucleic acids that function as molecular mimics of tRNAs^31^. An overlay of the structure of SidI’s domains A and B (excluding the 50 amino acids that form the enzymatic cleft; see Figure 2b) to the structure of a tRNA showed a tight fit into the space occupied by domains A and B (Figure 2h), with both showing a near 90-degree angle between the two arms (Figure 2f – g). This region of SidI is around 67 Å long and 65 Å wide, similar in both size and width of the tRNA which is 70 Å long and 60 Å wide (Figure 2f – h), although SidI is ∼30 Å thicker than the tRNA due to a four-helix bundle extending out from domain A (Figure 2h). Similarly, we compared the structure of SidI to those of proteins that are known to function as tRNA mimics in cells, such as the bacterial elongation factor EF-G (eEF2 in mammals)^32,33^ and the mammalian release factor eRF1^34^. In addition to the resemblance in sizes and shapes (Figure S3d – g), we found a stretch of positively charged amino acids in EF-G and eRF1 that form contacts with the ribosome^34–36^ (Figure S3h – i; boxes). This feature is also shared by SidI (Figure 2i; box), which might enable it to interact with the negatively charged backbone phosphates of ribosomal RNA.

These results together suggest that the SidI is a multidomain protein with its C-terminal part adopting a GT-B fold which gives it the ability to bind and hydrolyze GDP-mannose, and the N-terminal part adopting a shape closely mimicking a tRNA. Accordingly, we henceforth refer to this domain as SidI’s ‘tRNA-mimicry domain’ (Figure 2j). Notably, the tRNA mimicry domain provides a plausible explanation of the results from our AP-MS experiments (Figure 1c – e; Figure S1c – d), which showed SidI’s ability to bind to molecules that also transiently bind to tRNAs in cells (ribosomes, elongation factors and tRNA ligases).

### Molecular coupling of tRNA-like shape and glycosyl transferase activity may explain SidI potency

To address if and how the above structural features of SidI impact its function, we attempted to decouple the tRNA-like shape adopted by SidI from its enzymatic activity by introducing point mutations in its enzymatic pocket. We presumed that such mutations in SidI would only affect its enzymatic function without perturbing its overall fold. To this end, we examined the amino acids constituting the putative GDP-mannose binding pocket of SidI and focused our attention on the amino acids K600, D724 and R453 (Figure 2e). K600 and D724 are conserved in the catalytic pocket of PimB’ (Figure S3b) and single point mutations in the analogous residues K211Q and E290N on PimB’ result in significant defects in GDP-mannose binding efficiency and enzymatic function, suggesting that these residues help in accommodating GDP-mannose within the enzymatic pocket^30^. R453 does not appear to contact GDP-mannose directly. However, it is involved in a network of interactions across domains A and B, primarily ionic interactions with D19, D456 and E482, as well as a cation-π interaction with F11 (Figure 2e). These interactions could allow it to stabilize the local conformation, thus enabling the enzymatic pocket to be correctly positioned for catalysis. Indeed, previous studies have shown that a R453P mutation significantly attenuates SidI function^10,22^.

Guided by these considerations, we expressed and purified GST-SidI bearing R453P, K600Q and D724N mutations and determined that these mutant proteins were defective at hydrolyzing GDP-mannose (Figure 2k). Furthermore, all three enzymatically defective mutants were ∼1000-fold less potent (IC_50_ R453P = ∼0.85 µM; IC_50_ K600Q = 0.22 µM; IC_50_ D724N = 0.25 µM) at inhibiting the translation of luciferase mRNA in cell free RRL extracts in comparison to SidI (Figure S4a). However, these mutant proteins still reduced translation efficiency to near zero values at the highest concentration tested, which prompted us to explore whether additional features could contribute to SidI’s function. Indeed, pulldown of both GST-SidI and the enzymatic-deficient GST-SidI R453P robustly enriched purified 80S ribosomes (Figure S4b), as well as a repertoire of tRNA ligases and eEF1A from HEK293T lysates (Figure S4c), supporting the notion that SidI’s interaction with its targets is dependent on its tRNA-like shape, and not on its enzymatic activity. These results, together, suggest that while the tRNA-mimicry domain of SidI might confer the specificity to target components of the translation machinery, the glycosyl transferase activity of SidI might provide its potency to inhibit protein synthesis.

To test this notion, we first pre-treated RRL extracts with SidI or SidI R453P at 1nM (IC_50_ of SidI) for varying durations. In addition, we also pre-treated RRL extracts with SidI R453P at 0.5µM (IC_50_ of SidI R453P). We then utilized these pre-treated extracts to translate luciferase mRNA *in vitro* for another 60 minutes. Preincubation of RRL extracts with SidI resulted in a progressive reduction of translation efficiency relative to the increase of SidI pre-treatment time (Figure 2l), consistent with the model that the enzymatic action of SidI is required for its potency. By contrast, preincubation with SidI R453P at the same concentration or at an ∼1000-fold higher concentration did not result in a changed translation efficiency at all times tested (Figure 2l), supporting the notion that SidI’s enzymatic function and not just its tRNA-like shape is important to potently inhibit protein synthesis.

Guided by our AP-MS and pulldown experiments (Figure 1c – e), we reasoned that ribosomes and associated translation factors might serve as substrates for SidI mediated glycosylation. It is understood that the addition of acceptor substrates to a glycosyl transferase reaction accelerates the hydrolysis of the nucleotide-sugar precursor by the enzyme^37^. We hence prepared crude ribosome-enriched cytosolic extracts from HEK293T cells by sucrose gradient centrifugation to enrich for ribosomes and factors co-sedimenting with these ribosome populations. We then tested for GDP-mannose hydrolysis by co-incubating SidI and ribosomes. Indeed, incubation of increasing concentrations of crude ribosome extracts with SidI but not SidI R453P resulted in a dose-dependent increase in the rate of GDP hydrolysis from GDP-mannose (Figure S5a), suggesting that one or more factor(s) present in these extracts is a substrate of SidI.

We next determined if some of the ribosomal proteins and translation factors that bound to SidI exhibited affinity to the mannose-binding lectin concanavalin-A (Con-A). To this end, we treated the ribosome extracts with either GST, GST-SidI or GST-SidI R453P both in the absence or presence of GDP-mannose, pulled-down mannosylated molecules with Con-A coupled to Sepharose beads and subjected the eluted fractions to SDS-PAGE and immunoblotting with antibodies against GST, specific ribosomal proteins, and eEF1A (Figure 2m, schematic). Con-A only bound to the ribosomal proteins uL4, uL3 and eS8, and also to eEF1A strongly when the ribosome extracts were incubated with GST-SidI and GDP-mannose (Figure 2m, lane 6). Incubation of ribosome extracts containing GDP-mannose with either GST or GST-SidI R453P resulted in little to no enrichment of the proteins by Con-A (Figure 2m, lanes 4 and 8). Interestingly, Con-A also only bound to GST-SidI strongly under conditions where the toxin was incubated with GDP-mannose (Figure 2m, lanes 6 and 10), suggesting that SidI itself becomes auto-mannosylated. Indeed, several *L.p.* effectors with enzymatic domains show evidence of autocatalysis^38–40^. In sum, our results suggest that SidI enzymatically modifies core components of the translation machinery, including the ribosome and eEF1A, to inhibit protein synthesis.

We reaffirmed these findings by experiments performed in commercially available cell-free translation extracts (S30) isolated from *E.coli*. Incubating SidI or SidI R453P with S30 extracts resulted in a modest inhibition of luciferase mRNA translation at micromolar concentrations (Figure S5b; no GDP-mannose). Supplementing these extracts with GDP-mannose vastly accelerated translation inhibition under reaction conditions where SidI but not SidI R453P was present (Figure S5b; + GDP-mannose), indicative of a conserved mechanism by which SidI inhibits protein synthesis in both bacteria and eukaryotes. These findings also give credence to the studies that describe the role of an antitoxin MesI encoded by *L.p.* that limits SidI-induced toxicity^22,41,42^ (see Discussion). We note that supplementing eukaryotic RRL extracts with exogenously added GDP-mannose did not influence the rates of translation inhibition induced by SidI (Figure S5c), likely because GDP-mannose is not limiting in these preparations.

### Elongation defects caused by SidI result in ribosome stalling

In line with the above *in vitro* data, in HEK293T cells expressing equal amounts of FLAG-SidI and FLAG-SidI R453P, only FLAG-SidI expression resulted in a substantial reduction of nascent protein synthesis rates (Figure 2n). We tested whether this reduction was indeed caused by a block in translation elongation by subjecting lysates collected from these cells to polysome profiling. Polysome sedimentation in sucrose gradients provides a measure of the number of ribosomes per mRNA. Perturbations that result in elongation defects result in the association of multiple ribosomes per mRNA, thus causing an increase in the fraction of sedimented polysomes. Consistent with the notion that SidI acts at the elongation rather than initiation steps of protein synthesis, FLAG-SidI expression in cells caused a significant increase in the number of sedimented polysomes while FLAG-SidI R453P expression did not (Figure 2o), indicating an accumulation of stalled ribosomes on mRNAs in these cells.

### SidI activates a ribosome-associated stress response in cells

Perturbations to the protein synthesis machinery that result in ribosome stalling often activate signaling and transcriptional quality control programs that help cells adapt and respond to the perceived insult^43–46^. We reasoned that *L.p.*, by way of secreting SidI, might activate some of these programs in infected cells (Figure 3a). To that end, we first employed next-generation RNA sequencing (RNA-seq) to reveal the changes in mRNA abundance in Fc gamma receptor (FcγR) expressing HEK293 (HEK293 FcγR) cells infected either with wild type (WT) *L.p.* or an isogenic T4SS defective mutant of *L.p.* (*ΔdotA L.p.*) (Figure 3b) (Table S3). FcγR is stably expressed in these cells to allow for the efficient uptake of *L.p.* (see Methods). We used *ΔdotA L.p.* as a negative control for effector-mediated processes because it fails to secrete any effectors into host cells and eventually ends up getting degraded in lysosomes^47,48^. Additionally, in independent experiments, we induced an accumulation of stalled ribosomes in HEK293T cells by treating them with intermediate doses (0.1µM) of the elongation inhibitors anisomycin (ANS)^44,49^ and didemnin-B (DD-B)^49,50^ and analysed their respective transcriptomes by RNA-seq as well (Figure 3b). We chose these drugs because the act by entirely different mechanisms. ANS binds directly to the 60S subunit of ribosome and inhibits the peptidyl transferase reaction^51^, while DD-B binds to eEF1A and locks it in a post-hydrolysis GDP-bound state, preventing amino-acyl tRNA release and eEF1A dissociation from the ribosome^50^. We then compared and contrasted the transcriptional profiles of cells exposed to these different treatments.

**Figure 3.**
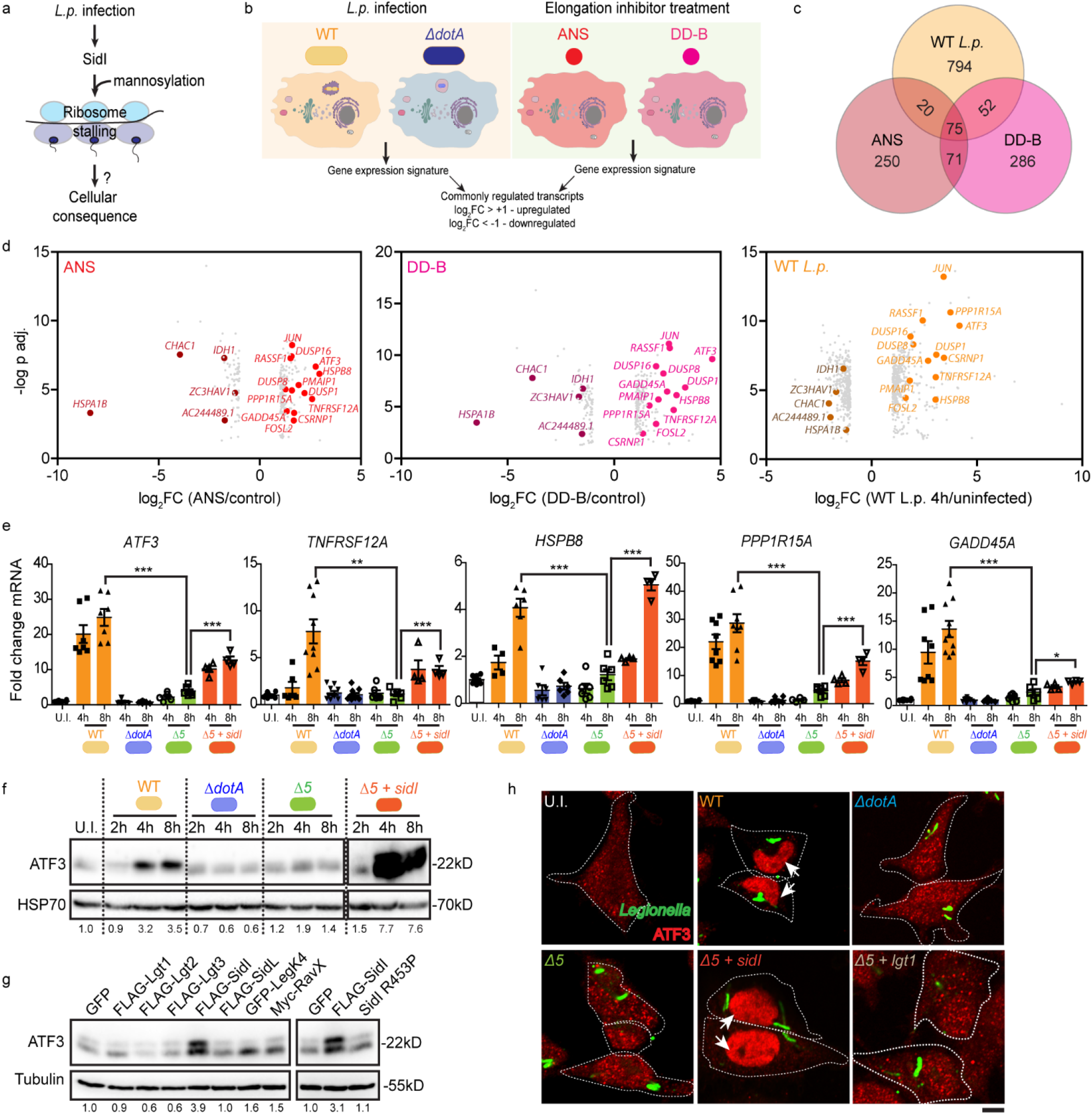
Transcriptional profiling reveals a conserved stress signature activated by SidI-mediated ribosome stress. (a – b) Schematic model of the RNA sequencing experimental pipeline to interrogate the cellular response to ribosome stalling events. Expression profiles of HEK293 FcγR cells infected with WT or *ΔdotA L.p.* (MOI = 50) and HEK293T cells treated with anisomycin (ANS, 0.1µM) or didemnin-B (DD-B, 0.1µM) were compared. (c) Venn diagram and (d) log_2_fold change induction of differentially expressed co-regulated transcripts between ANS and DD-B treatments for 4 hours in HEK293T cells and WT *L.p.* infection after 4 hours in HEK293 FcγR cells. Values were normalized to respective controls (DMSO for drug treatments and uninfected cells for *L.p.* infections) (n = 2 independent replicates per condition). Upregulated transcripts that map to stress signaling and cell death pathways are highlighted. p-values were calculated using the LIMMA Bioconductor package (5% false discovery rate with an effect size of at least 2x in comparison to controls). (e) Fold change induction of stress inducible transcripts measured by qRT-PCR in HEK293 FcγR cells infected with WT, *ΔdotA*, *Δ5 or Δ5* + *sidI L.p* strains (MOI = 50; Data represent mean ± standard error of the mean; n > 3 replicates per condition). Transcript expression values were normalized internally to the reference gene *RPS29* and expressed as fold change over the levels in uninfected controls. p-values were calculated by students t-test. **p<0.01; ***p<0.001. (f) Immunoblotting of ATF3 in lysates from HEK293 FcγR cells infected with WT, *ΔdotA*, *Δ5 or Δ5* + *sidI L.p.* strains (MOI = 50). Data are representative of 3 independent experiments. (g) Immunoblotting of ATF3 in lysates from cells expressing GFP, tagged-*L.p.* toxins or FLAG-SidI R453P. Data are representative of 2 independent experiments. (h) Indirect immunofluorescence of ATF3 and *L.p.* in HEK293 FcγR cells infected with WT, *ΔdotA*, *Δ5, Δ5* + *sidI* or *Δ5 + lgt1 L.p.* strains (MOI = 5) for 6 hours. Scale bar = 10µM. Data are representative of 2 independent experiments. Nuclear region is marked by arrows.

Principle component analysis (PCA) on the experimental replicates showed that the transcriptomes of cells treated with ANS- (red circles) or DD-B- (pink circles) displayed a high degree of similarity and clustered together, indicating that the cellular response to ribosome stalling elicited a conserved signature (Figure S6a). The transcriptional signatures of *L.p.* infected cells, although indicative of a general host response to bacterial infection by gene set enrichment analyses^52^ (Table S4), differed in a manner dependent on the duration of infection (Figure S6a, labelled orange circles) and varied considerably from the transcriptomes of ANS and DD-B treated cells (Figure S6a). Despite the variance in the global transcriptional profiles, a conserved set of 75 genes was significantly co-regulated in cells infected for 4 hours with *L.p.* and in cells treated for 4 hours with either ANS or DD-B (Figure 3c) (Table S3). Approximately 20% of the upregulated transcripts (13/61) mapped to pathways involved in stress signaling and cell death (Figure 3d, filled red, pink, and orange circles) while no specific commonalities could be attributed to the downregulated genes. Profiling the expression changes of these transcripts for up to 8 hours of *L.p.* infection revealed a dynamic regulation of their abundance (Figure S6b and S7a; WT) that was dependent on the secretion of effectors via a functional T4SS (Figure S6b and S7a; *ΔdotA*). We confirmed these findings with a targeted validation of five highly upregulated transcripts (*ATF3*, *TNFRSF12A*, *HSPB8*, *PPP1R15A* and *GADD45A*) by quantitative real time polymerase chain reaction (qRT-PCR) assays. WT *L.p.* but not *ΔdotA L.p.* infected HEK293 FcγR cells activated a robust increase in the mRNA abundance of all five transcripts that peaked at 8 hours post infection in comparison to uninfected cells (Figure 3e, compare white, orange and blue bars). Significantly, these transcripts were also abundantly induced in HEK293T cells treated with mild doses of ANS and DD-B (Figure S8a, red and pink bars), suggesting that their increase is dependent on an accumulation of stalled ribosomes.

As further controls, we expanded the repertoire of translation inhibitors to include cycloheximide (CHX; ribosome E-site blocker)^51^, deoxynivalenol (DON; ribosome A-site blocker)^51^ and harringtonine (HTN; initiation and elongation inhibitor)^51,53,54^, and tested their respective abilities to activate stress inducible gene expression in cells utilizing a dosage regimen of 0.01µM to 10µM. While DON treatment strongly activated transcript induction at specific doses (Figure S8a, purple bars), CHX and HTN treatments resulted in poor induction of transcripts at all tested concentrations (Figure S8a light and dark brown bars). In light of these results, we note that ANS, DD-B and DON specifically interfere with an early step of the translation elongation cycle^50,51^, suggesting that an inhibition of this particular step renders cells hyper-sensitive to stalled ribosome stress. Moreover, a suppressed response to CHX or HTN excludes a causal role for general protein synthesis inhibition to activate an upregulation of these specific transcripts. Concordant with our data, the heterologous expression in HEK293T cells of the toxin FLAG-SidI, that induces ribosome stalling (Figure 2o), but not FLAG-SidI R453P, also resulted in a significant increase of all five stress inducible mRNAs (Figure S9a – e), supporting the notion that SidI is necessary and sufficient to induce these stress transcripts.

To confirm if these observations parallel live *L.p.* infections, we utilized an isogenic mutant strain of *L.p.* with genomic deletions of five EITs, including *sidI* (*Δlgt1*, *Δlgt2*, *Δlgt3*, *ΔsidI* and *ΔsidL*; hereafter *Δ5 L.p.*)^8^ and a *Δ5* strain complemented with a plasmid encoding only *sidI* (*Δ5*+*sidI L.p.*). Cells infected with *Δ5 L.p.* exhibited minimal changes in stress transcript abundance, with mRNA levels similar to those observed during infections with *ΔdotA L.p.* (Figures 3e; green bars). Importantly, cells infected with the complemented *Δ5*+*sidI L.p.* strain activated all five stress inducible transcripts to levels significantly higher than those observed during *Δ5 L.p.* infections (Figure 3e; orange bars) and to levels similar to those observed during expression of FLAG-SidI in cells (Figure S9a – e), albeit not to the quantities observed during infections with WT *L.p.* for all transcripts except *HSPB8* (Figure 3e; orange and dark red bars). These results indicate that, while SidI secretion is sufficient for the stress transcript induction during *L.p.* infection, other possible T4SS effectors also regulate their abundance.

Interestingly, both our RNA-seq datasets and qRT-PCR analyses pointed to *ATF3* mRNA as the most differentially expressed transcript co-upregulated under all treatment conditions (Figure 3d – e). *ATF3* mRNA levels also significantly rose as early as 2 hours post-infection with *L.p.* and remained at sustained high levels for up to 8 hours (see Figure S6b). Its protein product, the activating transcription factor ATF3, is a member of the ATF/cyclic AMP response element binding (CREB) family of transcription factors that is induced in response to a variety of stress stimuli^55^. While *ATF3* mRNA is robustly induced by *L.p.* infections, heterologous SidI expression and treatments with ANS, DD-B and DON, we wondered if ATF3 protein levels are induced under the same conditions, as these treatments concomitantly reduce nascent protein synthesis rates in cells (see Figure 2n and Figure S10a – c). To address this paradox, we determined that *ATF3* mRNA escapes the imposed translation block and ATF3 accumulates in the nucleus of cells, concordant with its role as a transcription factor. We observed a striking accumulation of ATF3 by immunoblotting of lysates collected after infections with the WT and the *Δ5*+*sidI L.p.* strains, but not the *ΔdotA* and *Δ5* strains (Figure 3f). ATF3 accumulation was also observed with the expression of FLAG-SidI but not the other *L.p.* EITs (Figure 3g, lanes 1 – 8), nor the enzymatic deficient mutant FLAG-SidI R453P (Figure 3g, lanes 9 – 11); and with the treatment of cells with mild doses of ANS, DD-B and DON, but not CHX and HTN (Figure S8b). Furthermore, by indirect immunofluorescence and confocal microscopy analyses, we visualized the translocation of ATF3 from the cytosol in control cells to the nucleus in cells that were infected with the WT and the *Δ5*+*sidI L.p.* strains, as well as in cells treated with a mild dose of ANS, DD-B or DON (Figure S8c; white arrows). ATF3, however, remained diffuse in cells infected with the *ΔdotA, Δ5* and *Δ5+lgt1* (*Δ5* with a plasmid encoding *lgt1*) strains (Figure 3h; arrows). Taken together, our results suggest that cells, in response to a build-up of stalled ribosomes, activate a conserved stress signature that results in the accumulation of ATF3 in the nucleus.

### ATF3 accumulation is dependent on stress activated kinase signaling from stalled ribosomes

Since ribosome stalling has been linked to the activation of multiple ribosome-associated stress response pathways^44–46^, we asked which of these pathways regulate the increase in ATF3 (Figure 4a; schematic). First, we tested the role of the integrated stress response (ISR), a signaling pathway that is turned on by the phosphorylation of serine 51 (S51) on the alpha subunit of the translation initiation factor eIF2^56^. Phosphorylated eIF2 acts as an inhibitor of eIF2’s guanine nucleotide exchange factor eIF2B, thereby reducing general protein synthesis while simultaneously activating the translation of specific mRNAs, including those encoding the transcription factors ATF4 and ATF3^56^. We challenged mouse embryonic fibroblasts (MEFs) that carried either the wild type eIF2α S51 alleles (ISR active) or mutant eIF2α S51A alleles (S51 mutated to alanine; ISR null)^57^ with mild doses of ANS for 4 hours and assayed for ATF3, ATF4 and eIF2α phosphorylation by immunoblotting. As controls, we treated cells with a known ISR activator thapsigargin (Tg) for 4 hours^58,59^. In accordance with published studies^59^, Tg activated the accumulation of both ATF3 and ATF4 in ISR active but not ISR-null MEFs (Figure 4b).

**Figure 4.**
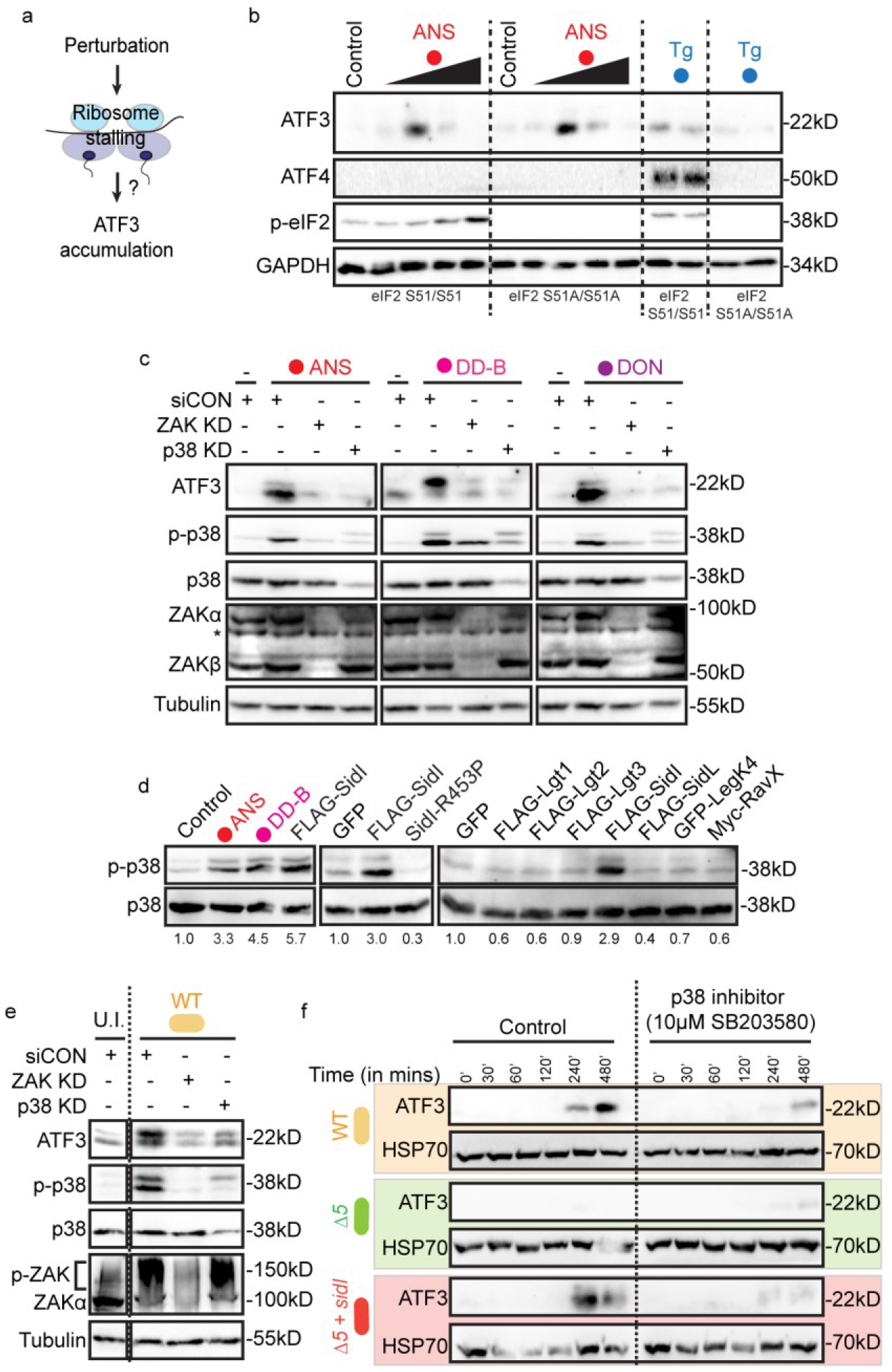
ATF3 accumulation is dependent on stress-activated kinase signaling from collided ribosomes. (a) Schematic model of the putative stress signaling pathway that results in ATF3 accumulation downstream of ribosome stalling. (b) Immunoblotting of ATF3, ATF4, phospho-eIF2 and GAPDH in lysates from WT or eIF2 *S51A^+/+^* mouse embryonic fibroblast cells treated with ANS (0.01µM, 0.1µM, 1µM and 10µM) or thapsigargin (Tg, 1µM) for 4 hours. (c) Immunoblotting of ATF3, phospho- and total p38, ZAK and Tubulin in HeLa cells depleted for ZAK or p38 by siRNA for 72 hours and treated with ANS (0.1µM), DD-B (0.1µM) or deoxynivalenol (DON, 1µM) for 4 hours. Data are representative of 3 independent experiments. *-unspecific band. (d) Immunoblotting of phospho- and total p38 in lysates from HEK293T cells treated with ANS and DD-B for 4 hours or cells expressing GFP, FLAG-SidI, FLAG-SidI R453P or epitope tagged *L.p.* toxins. Values below immunoblots depict densitometric quantification of phospho-p38 band intensities relative to total p38 and normalized to controls. Data are representative of 2 independent experiments. (e) Immunoblotting of ATF3, phospho-p38, total p38, ZAKα and Tubulin in lysates from HEK293 FcγR cells treated with control siRNA or siRNAs against ZAK and p38 for 72 hours followed by infection with WT *L.p.* (MOI = 50) for 4 hours. Data are representative of 2 independent experiments. Lanes were cropped from the same gel. (f) Immunoblotting of ATF3 and HSP70 in lysates from HEK293 FcγR cells infected with WT, *Δ5* or *Δ5+sidI L.p.* strains and treated with p38 inhibitor (SB203580, 10µM). Data are representative of 2 independent experiments.

Surprisingly, ANS induced a marked accumulation of ATF3 in both ISR active and ISR-null MEFs, while ATF4 accumulation was entirely absent (Figure 4b). ATF3 accumulation also occurred when MEFs were challenged with mild doses of DD-B, with a more pronounced increase of ATF3 in the ISR null MEF cells (Figure S11a). Furthermore, treatment of HEK293T cells with the ISR inhibitor ISRIB^60^ also failed to suppress the increases of ATF3 when these cells were treated with ANS or transfected with FLAG-SidI (Figure S11b). In sum, these results argue against a role for the ISR pathway to enable ATF3 accumulation in response to ribosome stalls.

Together with the ISR, a distinct stress-signaling pathway is also activated when ribosomes undergo collisions. Collided ribosomes recruit and activate the leucine zipper and sterile α motif stress kinase ZAKα (MAP3K20), which in turn phosphorylates the stress kinase p38 via pathway termed as the ribotoxic stress response (RSR)^44,45,61^. Indeed, our treatment regimen of ANS, DD-B and DON in HEK293T cells strongly activated p38 phosphorylation (p-p38) as assessed by immunoblotting, while corresponding treatments with CHX and HTN did not (Figure S11c). We also detected similar increases of p-p38 and ATF3 in HeLa cells treated with an intermediate dose of ANS, DD-B and DON (Figure 4c, compare lanes 1 and 2; 5 and 6; 9 and 10). Depleting these cells of ZAK (ZAK knockdown; ZAK KD) with small interfering RNA (siRNA) suppressed the drug induced increases of both p-p38 and ATF3 (Figure 4c, lanes 3,7,11). Concordantly, p38 depletion with siRNA also significantly inhibited the accumulation of ATF3 (Figure 4c, lanes 4,8,12). These results suggested that ZAKα is the upstream kinase that phosphorylates p38 to regulate ATF3 induction.

In line with these findings, FLAG-SidI expression in HEK293T cells also activated a marked increase in p-p38 levels, similar to the intensity elicited by ANS and DD-B treatments (Figure 4d, first four lanes) and dependent on SidI’s glycosyl tranferase activity (Figure 4d, middle 3 lanes). Furthermore, the expression of only SidI but not the other *L.p.* EITs strongly activated p38 phosphorylation in HEK293T cells (Figure 4d, last 8 lanes).

In an *L.p.* infection context, global phosphoproteomics analyses in HEK293 FcγR cells from our lab^62-preprint^ indicated that as early as one hour post infection, *L.p.* induced a significant activation of the stress activated protein kinase pathway in infected cells, as determined by an enrichment of substrates phosphorylated by the stress kinases p38α, p38β and MAPKAP2 (MK2) (Figure S11d, Table S5). This result is supported by previous data suggesting that p38 is activated after WT *L.p.*^63^, but not the *Δ5 L.p.* infection^15^. We confirmed these results by determining that *L.p.* robustly induced the phosphorylation and activation of p38 (p-p38) and ZAKα (high molecular weight species; p-ZAK) in infected cells (Figure 4e; lanes 1 and 2). p38 activation in infected cells was largely dependent on the secretion of SidI by *L.p.* (Figure S11e, compare WT, *Δ5* and *Δ5*+*sidI* panels). Furthermore, cells depleted of ZAKα and p38 by siRNAs or treated with a small molecule p38 inhibitor failed to accumulate ATF3 in response to a *L.p.* challenge (Figures 4e-f). We surmise from these data that p38 signaling in *L.p.* infected cells primarily initiates from stalled ribosomes and that the RSR regulates ATF3 accumulation.

### ATF3 orchestrates a transcriptional program that regulates cell fate

We next focused on understanding the significance of the stress-induced ATF3 accumulation in cells. As alluded to earlier, ATF3 belongs to an ‘expanded’ family of basic leucine zipper (bZIP) domain containing AP-1 transcription factors^55,64,65^ and binds to a diverse array of cyclic AMP response elements (CRE) on the promoters of genes under both basal and stress-induced conditions to regulate their expression^65^. Paradoxically, ATF3 has been shown to function as both a transcriptional activator and a repressor in different cellular contexts, dependent on its protein isoform, its ability to homodimerize (repressor), and/or its ability to form heterodimers with other AP-1 family members, including JUN and FOS^55,65^. We noted that the mRNA levels of both *JUN* and *FOSL2* were also significantly upregulated in response to ribosome stalling events (Figures 3d and S6b), suggesting that ATF3, in this context, might function as a transcriptional activator as previously reported^55,66^. Upon mining genome-wide chromatin immunoprecipitation and next generation sequencing (ChIP-seq) datasets collected from four diverse cell types, we found that ATF3 bound to euchromatin rich genomic regions in one or more cell type (marked by acetylated histone H3K27Ac), at or near the exons of genes coding for its own mRNA (*ATF3*), as well as the mRNAs of *TNFRSF12A*, *HSPB8*, *PPP1R15A* and *GADD45A* (Figure S12).

Prompted by this observation, we next tested whether ATF3 regulated the expression of the latter four upregulated mRNAs in response to ribosome stalling events. Towards this end, we generated an ATF3 knockout (KO) HEK293T cell line using clustered regularly interspaced short palindromic repeat (CRISPR)/Cas9 mediated mutagenesis by targeting genomic regions in exon 2 of the *ATF3* gene using a pool of three small guide RNAs (Figure 5a). We validated a complete loss of ATF3 protein levels in these cells under both basal and ANS treatment conditions by immunoblotting (Figure 5b). We then challenged these cells and the isogenic HEK293T parental cells with FLAG-SidI, FLAG-SidI R453P or a mild ANS dose and assayed for the mRNA levels of the four stress-inducible transcripts by qRT-PCR. As expected, FLAG-SidI expression and ANS treatment led to a strong induction in the mRNA levels of all four genes in the parental cell line (Figure 5c – f and Figure S13a – d). In ATF3 KO cells, however, stress transcript induction was attenuated (ATF3 KO panels, Figure 5c – f and S13a – d). Importantly, the expression of an enzymatically deficient SidI mutant (FLAG-SidI R453P) in both cell lines failed to activate mRNA induction (Figure 5c – f). These results strongly support for a role of ATF3 as a master transcription factor that regulates gene expression in response to ribosome stalling.

**Figure 5.**
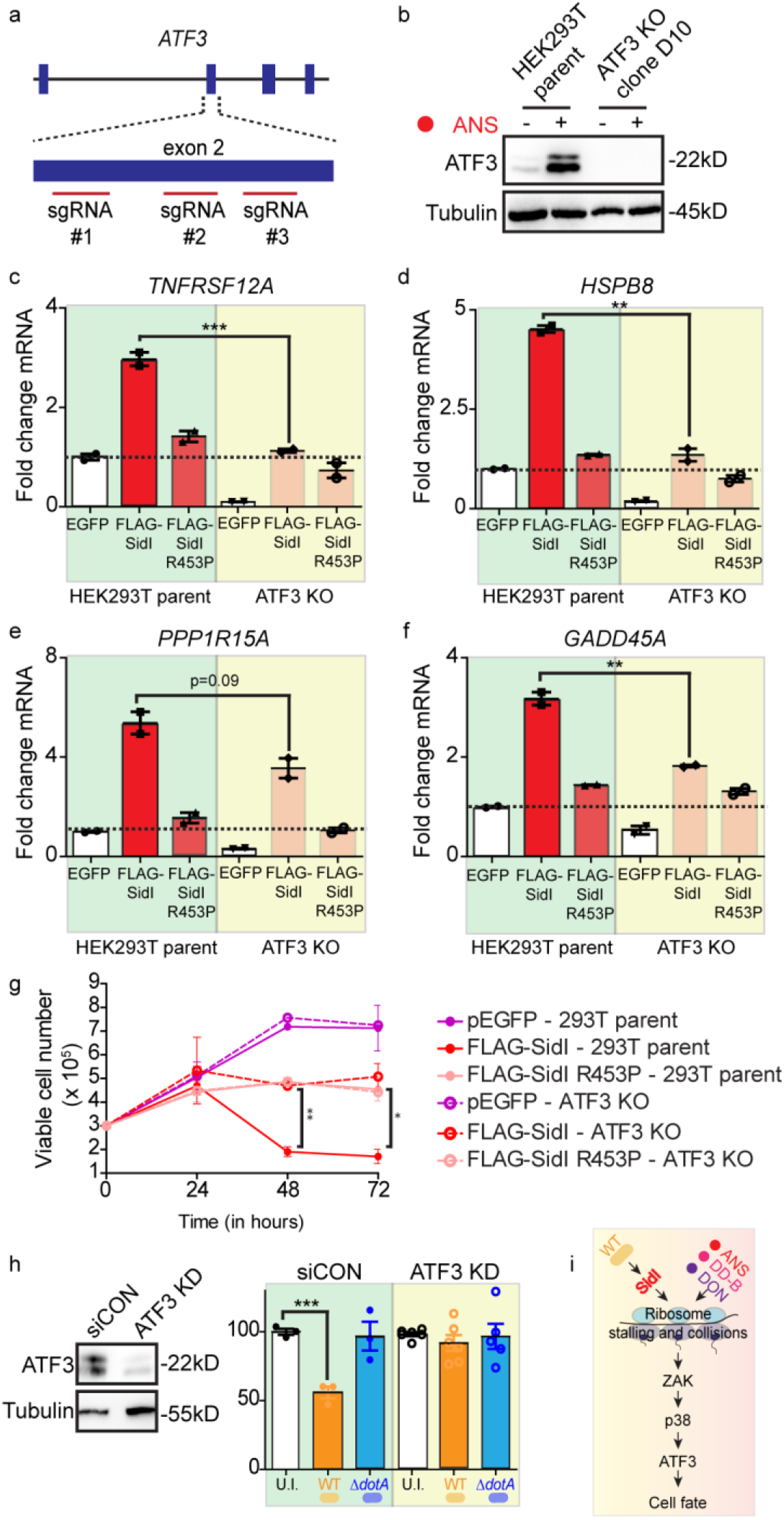
ATF3 orchestrates a transcriptional program that regulates cell fate. (a) Schematic of the strategy for CRISPR-Cas9 mediated depletion of ATF3 in HEK293T cells. (b) Immunoblotting of ATF3 in lysates from HEK293T parent or ATF3 knockout (ATF3 KO) HEK293T cells at steady state and after challenge with ANS (0.1µM) for 4 hours. (c – f) Fold change induction of *TNFRSF12A*, *HSPB8*, *PPP1R15A* and *GADD45A* mRNA in parent and ATF3 KO HEK293T cells expressing GFP, FLAG-SidI or FLAG-SidI R453P measured by qRT-PCR (Data represent mean ± standard error of the mean; n = 2 independent experiments). Transcript expression values were normalized internally to the reference gene *RPS29* and expressed as fold change over the levels in GFP expressing cells. p values calculated by students t-test. **p<0.01; ***p<0.001. (g) Cell viability measurements in parent and ATF3 KO HEK293T cells expressing GFP, FLAG-SidI or FLAG-SidI R453P (Data represent mean ± standard error of the mean; n = 2 independent replicates per condition). p-values were calculated by students t-test. *p<0.05; **p<0.01. (h) Cell viability measurements in HEK293 FcγR cells treated with control and ATF3 siRNAs for 72 hours and infected with WT or *ΔdotA L.p.* for 10 hours (Data represent mean ± standard error of the mean; n ≥ 3 independent replicates per condition). p-values were calculated by students t-test. ***p<0.001. Immunoblot for ATF3 depicts knockdown efficiency. (i) Schematic model of the ribosome-associated stress response pathway activated by colliding ribosome stress.

As noted earlier, a significant fraction of co-upregulated transcripts during *L.p.* infection mapped to cell death pathways (Figure 3c). As SidI expression results in cell death^10^, we tested whether ATF3’s role as a transcriptional regulator influenced this outcome. We hence monitored the viability of parental and ATF3 KO cells transiently expressing GFP, FLAG-SidI or FLAG-SidI R453P for three days. In both cell types, GFP expression did not interfere with cellular growth, with a progressive doubling of viable cells that plateaued at day 3 post-transfection (Figure 5g, purple solid and dashed lines). FLAG-SidI R453P expression resulted in similar rates of cell growth in both parental and ATF3 KO cells, but at slower rates when compared to GFP expressing cells (Figure 5g, pale coloured solid and dashed lines). In contrast, FLAG-SidI expression in parental 293T cells resulted in a severe reduction of viability at days 2 and 3 post transfection (Figure 5g, solid red line). However, a depletion of ATF3 rescued cell viability in cells expressing FLAG-SidI, with these ATF3 KO cells mirroring the growth rates observed with cells expressing the enzymatic deficient mutant R453P (Figure 5g, dashed red line). In sum, these results indicate that (i) SidI has a cytopathic effect on cellular growth via ATF3; (ii) In the absence of ATF3, SidI causes cytostasis; and (iii) enzymatically defective SidI also causes cytostasis independent of ATF3 function, likely due to its inhibitory effect on protein synthesis (see Figure S4a).

Finally, we tested the role of ATF3 during *L.p.* infections in HEK293 FcγR cells. To this end, we first established that challenging these cells for 10 hours with a large pathogenic burden (multiplicity of infection = 50) of WT *L.p* resulted in significant reduction of cell viability as compared to uninfected controls (Figure 5h and S13e; orange bars). This result suggested that such high amounts of *L.p.* replicating for ten hours would recapitulate the final stages of its intracellular replication cycle that ultimately leads to cell lysis^67–69^. In contrast, challenging cells with either the *ΔdotA L.p.* or *Δ5 L.p.* strains resulted in minimal cell death (Figure S13e; blue and green bars). Strikingly, infecting cells with the complemented *Δ5+sidI L.p.* strain caused a similar reduction of cell viability to that observed during infections with WT *L.p.* (Figure S13e, dark red bars), indicating that the secretion of SidI by *L.p.* is sufficient for this phenomenon. We then depleted ATF3 with siRNAs in HEK293 FcγR cells (Figure 5h; blot) and challenged these cells with WT *L.p.* or *ΔdotA L.p.* for ten hours. Indeed, in control siRNA treated cells, WT *L.p.* but not *ΔdotA L.p.* led to an almost fifty percent reduction in the number of viable cells when compared to uninfected controls (Figure 5h, siCON orange bar). In ATF3 depleted cells, however, WT *L.p.* infections failed to reduce cell viability (Figure 5h, ATF3 KD orange bar), underscoring the notion that ATF3 accumulation activated during *L.p.* infections affects a terminal program that regulates cell fate.

## Discussion

Understanding the principles that govern the interaction of infectious microorganisms with their host counterparts requires a mechanistic elucidation of: a) the toxins derived from the pathogen, b) the targets of the toxin in host cells, and c) the cellular consequences evoked by the toxin-target relationship. In our study of the *L.p.* toxin SidI, we present multiple lines of structural, biochemical, and genetic evidence that addresses each of these three considerations (see Figure S14 for a schematic model).

### The toxin

SidI was first discovered in a heterologous expression screen for *L.p.* effectors that cause cytotoxicity in yeast cells^10^. Follow up studies established that SidI is secreted into the cytosol of infected cells in a T4SS dependent manner^10,22^ and contributes to protein synthesis inhibition by slowing translation elongation rates^5^. Furthermore, SidI functions as a glycosyl hydrolase by cleaving GDP from GDP-mannose^22^. Building upon these previous findings, our structural characterization of SidI revealed a tRNA mimicry domain in its N-terminus and a glycosyl transferase domain in its C-terminus (Figure 2). Only a few proteins have been identified that mimic tRNAs, and these are present in both prokaryotes and eukaryotes and generally function in regulating the steps of translation elongation and termination^31,32,34^. They include the elongation factor EF-G/EEF2^32,33,70,71^, the termination factors RF1-3/eRF1/3^34,72^ and the ribosome recycling factors RRF^73^ and Pelota^74^. Each of these proteins possess domains that strongly mimic the shape and size of tRNAs^32–34,71,73,74^ and exhibit an electrostatic charge distribution that enables them to contact the ribosome^31,34,36,50^, despite not sharing much sequence homology with each other. These structural resemblances are also found in the tRNA mimicry domain of SidI (Figure 2f – i and S3d – i). Another striking feature of SidI is its enzymatic domain, which shares the fold and some conserved catalytic amino acids to known GDP mannosyl transferases (Figure 2b – e). It is this unique molecular coupling that endows SidI with the ability to directly target (via the tRNA mimicry domain) and enzymatically modify (via the GT domain) the host translation apparatus (Figures 1c – e, 2m), resulting in SidI being the most potent inhibitor of protein synthesis amongst the subset of *L.p.* EITs (Figure 1a – b, S4a). Mechanistically, the bipartite targeting and enzymatic capabilities of SidI mirros the mode of action exhibited by the toxin ricin, a heterodimeric ribosome-inactivating protein whose galactose binding B-chain enables targeting and uptake by cells and whose enzymatic A-chain functions as an N-glycoside hydrolase that depurinates adenine on the sarcin/ricin loop of 28S ribosomal RNA^20,75,76^. SidI, in similarity with ricin, requires an active enzymatic domain to potently inhibit protein synthesis in the picomolar range^20^ (Figure 1a – b and S4a).

Our *in vitro* assays show that SidI hydrolyzes GDP-mannose with high efficiency (Figure S3a). GDP-mannose is actively synthesized in the cytosol of mammalian cells^24^, and mannose concentrations are in the high micromolar range in circulating blood^77^. Thus, when GDP-mannose concentrations are non-limiting, as it is in the case of cell free translation extracts isolated from rabbit reticulocytes, SidI inhibits protein synthesis via an enzymatic poisoning of its targets (Figure 2l). To our surprise, this inhibition also occurs in cell free translation extracts isolated from the bacterium *E.coli* when they are supplemented with exogenous GDP-mannose (Figure S5b), indicative of the conserved nature of SidI’s targets in the protein synthesis apparatus in both prokaryotes and eukaryotes^78^. Interestingly, *L.p.* uniquely encodes an antitoxin named MesI (for meta-effector of SidI) to counteract the effects elicited by SidI^41^. MesI has been shown to directly bind to SidI inside bacterial cells^42^ and inhibit its GDP hydrolysis activity^22^. Indeed, *L.p.* mutants with a genomic deletion of the *mesI* locus fail to replicate intracellularly^41,42^, potentially due to the cytotoxic effect associated with unrestricted SidI activity.

### The targets

As central players in the process of protein synthesis, tRNAs in eukaryotic cells undergo processing by the tRNA ligase complex^79^ and amino-acylation (tRNA charging) by aminoacyl tRNA synthetases^80^. During protein synthesis, the tRNAs successively occupy privileged A- (aminoacyl), P- (peptidyl) and E- (exit) sites on the ribosome to facilitate translation elongation. The elongation of peptides on ribosomes is a cyclical, multi-step process that begins when the 80S ribosome sits on a start codon with the initiator methionyl-tRNA occupying the P-site, allowing for the delivery of mature ‘charged’ tRNAs by eEF1A to the A-site^50,81^. Upon successful accommodation, ribosomes catalyze the peptidyl transfer of amino acids from P-site tRNA onto the A-site tRNA^82^. The ribosome then moves three bases in the 5’-3’ direction along the length of the mRNA, thereby allowing the deacylated tRNA to occupy the E-site, the peptidyl tRNA to occupy the P-site and liberating the codon at the A-site for the next incoming charged tRNA to be accommodated^81^. This last step is catalyzed by the binding of elongation factor eEF2 to the ribosome A-site^83^. Thus, apart from tRNAs, tRNA mimicry proteins also transiently occupy the A-site of the ribosome to regulate translation elongation.

We present multiple lines of evidence here to indicate that SidI directly interferes with an early step of the elongation cycle. First, SidI shares a remarkable structural similarity to the domain architecture and overall fold of EF-G/eEF2^71,83^ (Figures 2a, 2f – h and S3f). Second, SidI precipitates multiple components of the cytosolic tRNA processing (RTCB and DDX1) and modification machinery (VARS and CARS) from cell extracts (Figure 1e, S1d and Table S1). Third, SidI directly interacts with ribosomes (Figure 1d) and eEF1A^10^ and catalyzes the mannosylation of the translation machinery (Figure 2m). Fourth, the expression of SidI in cells induces ribosome stalling in a manner that is dependent on its enzymatic function as a mannosyl transferase (Figure 2n – o). Fifth, SidI activates a stress response signature in cells that is exceptionally similar to one activated by the elongation inhibitors ANS, DD-B and DON that interfere with A-site function^50,51^, but not CHX, that occupies the E-site and inhibits tRNA translocation^51^ (Figure 3c – h, Figure 4, Figure S8 and Figure S9). We hence postulate that SidI acts by blocking either tRNA loading onto the A-site or the peptidyl transferase reaction to inhibit protein synthesis.

### Cellular consequences of the toxin-target relationship

As a general safeguard against various stressors, cells often employ signaling and/or transcriptional networks that aim to re-establish homeostasis or, if left unmitigated, induce cell death. This paradigm is typified by the ∼3 million ribosomes in eukaryotic cells^84^ that act as sentinels of cell health and function^44,85^. Indeed, a growing body of evidence posits that when cells are subjected to perturbations that induce ribosome stalling and collisions, ribosomes function as molecular ‘rheostats’ to gauge the intensity of the insult and mount specific adaptive responses^43–45,49^. Low basal levels of collisions are sensed and cleared by the ribosome quality control (RQC) pathway^43,44^, while intermediate and high levels of collisions that saturate RQC mediated events result in the induction of the ISR^44,46^ and the RSR^44,45,61,85^, respectively. In cells infected with *L.p.*, our data indicate that SidI secreted into the cytosol induces the RSR pathway that initiates on stalled ribosomes undergoing collisions, thereby activating the kinases ZAKα and p38^44,45^ and culminating in the accumulation of ATF3 in the nucleus of cells (Figure 3 and Figure 4). RSR activation is dependent on SidI’s mannosyl transferase activity (Figure 3g, Figure 4d and Figure S9). A feature of this pathway is the cellular reprogramming that allows for ATF3 protein build-up under conditions of the SidI/elongation inhibitor induced translation block (Figure 3f – h, Figure S8b). This type of regulation is unique to stress response pathways that simultaneously block a major fraction of protein synthesis in cells, while allowing for certain proteins to escape the imposed block and regulate downstream effects. Examples of such ‘escaper’ proteins include the chaperones that accumulate during the unfolded protein response (UPR)^86^, and the transcription factors ATF4 and CHOP that accumulate during the ISR^56^. While we cannot rule out an inhibition of degradative mechanisms that regulate ATF3 protein turnover^87,88^, we envision a scenario for the RSR wherein cis-regulatory determinants on the 5’- untranslated regions of the *ATF3* mRNA recruit the translation machinery and allow for its selective translation^59,89^. Notably, in cells challenged with large *L.p.* doses, ATF3 functions as a master transcription factor of the RSR pathway and regulates the activation of a cell death program, presumably via modulating specific gene expression in stressed cells (Figure 5 and Figure S13). We propose that high intracellular bacterial loads of *L.p.* induce a high ribosome collision rate that results in the induction of the RSR pathway. Furthermore, as ribosome collisions directly influence stress signaling^44,45,61,90^, immune gene expression^91^, cell cycle arrest^92^ and cell death^44^, we rationalize the association of these cellular outcomes to the secretion of the *L.p.* EITs. These findings complement earlier studies on host directed gene expression induced by low dose challenges of *L.p.* in macrophages^93^ and the amoeba *Dictyostelium discoideum*^94^. As SidI is largely dispensable for the intracellular replication of *L.p.*^8^, our results suggest that *L.p.*, via SidI, selectively hijacks the RSR pathway to activate the terminal program of cell lysis once bacterial replication is complete (Figure S14).

In conclusion, we report an unprecedented secreted bacterial toxin with a tRNA-like shape and glycosyl transferase activity, which we used to elucidate a ribosome-to-nuclear signaling pathway that regulates cell fate. As such, our study expands the portfolio of stress response pathway regulators derived from *L.p.*, underscoring *L.p.*’s amazing utility as a molecular tool box to uncover new mechanistic angles and fundamental aspects of host cell physiology^7,11, 16–18,38,39,95,96^.

## Acknowledgements

We thank all members of the S.M. and P.W. labs for critical evaluation and discussions of the data. We thank Srivats Venkatraman and Lorenzo Calviello from the S.F. lab for helpful discussions. We thank Julia Noack, CJ Sarabia, and Brian Wang, for technical assistance with the project. L.W. acknowledges a fellowship from the Damon Runyon Cancer Research Foundation (DRG-2312-17). SS is supported by Helen Hay Whitney Postdoc fellowship and K99 (1K99GM143527-01A1) from NIGMS. N.J.K acknowledges financial support from the National Institutes of Health (U19 AI135990). P.W. acknowledges financial support from the National Institutes of Health (R01GM032384) and the Howard Hughes Medical Institute. S.M. acknowledges financial support from the National Institutes of Health (R01GM140440), the Pew Charitable Trust (A129837) and a gift fund from the Chan-Zuckerberg Biohub. We thank the Vincent J. Coates Genomics Sequencing Laboratory at the California Institute for Quantitative Biosciences (QB3) for help with the RNA-seq experiments. We also thank the QB3 shared cluster for computational support.

## Author contributions

Conceptualization, A.Su., S.M.; Methodology, A.Su., L.W., T.M., M.V., S.S., D.L.S., N.J.K., S.F., A.Si., P.W., S.M.; Investigation, A.Su., L.W., T.M., M.V., E.S., E.H.P., S.K., D.L.S.; Formal Analysis, A.Su., L.W., M.V., D.L.S.; Writing – Original Draft, A.Su., L.W., S.M.; Writing – Review & Editing, A.Su., L.W., M.V., A.Si., P.W., S.M.; Visualization, A.Su., L.W., S.M.; Funding Acquisition, P.W., S.M.; Resources, S.S., S.F.; Supervision, A.Su., D.L.S., N.J.K., A.Si., P.W., S.M..

## Tables

**Table S1.** Mass spectrometry analyses of SidI interacting partners in HEK293T lysates and rabbit reticulocyte lysates.

**Table S2.** Cryo-EM data collection, reconstruction, and model refinement statistics.

**Table S3.** Log_2_fold changes of transcript expression levels in cells infected with *L.p.* or treated with ribosome stalling inducers ANS and DD-B.

**Table S4.** Gene set enrichment analyses on the ranked list of genes from the RNA-seq datasets.

**Table S5.** Inferred kinase activity scores in cells infected with *L.p.* for 1 hour by reanalysis of phosphoproteomics datasets in Noack *et al.*^62-preprint^.

**Table S6.** List of siRNAs and oligonucleotides.

## Materials & Correspondence

Further information and requests for resources and reagents will be fulfilled by Shaeri Mukherjee.

## Methods

### Key Resources Table

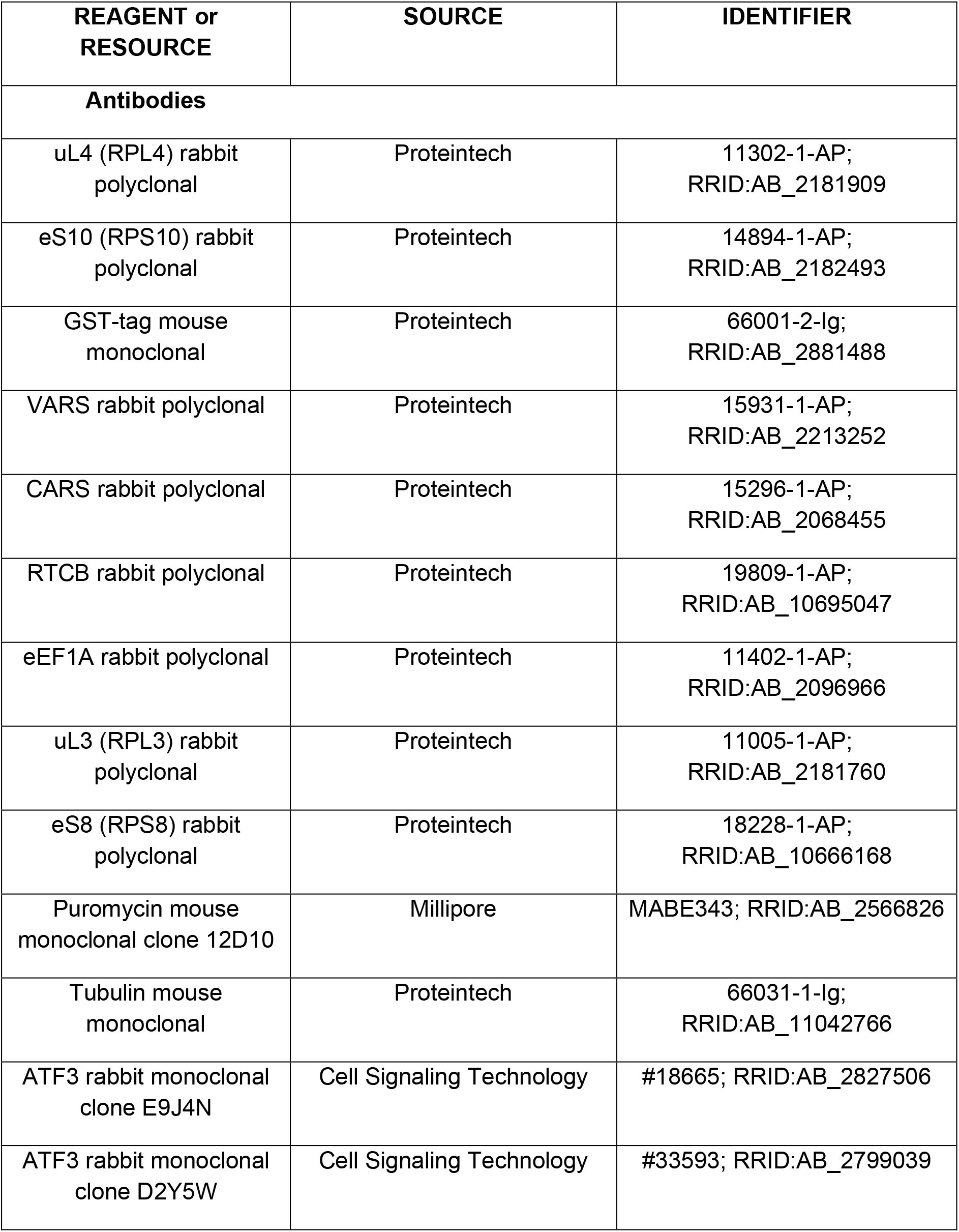
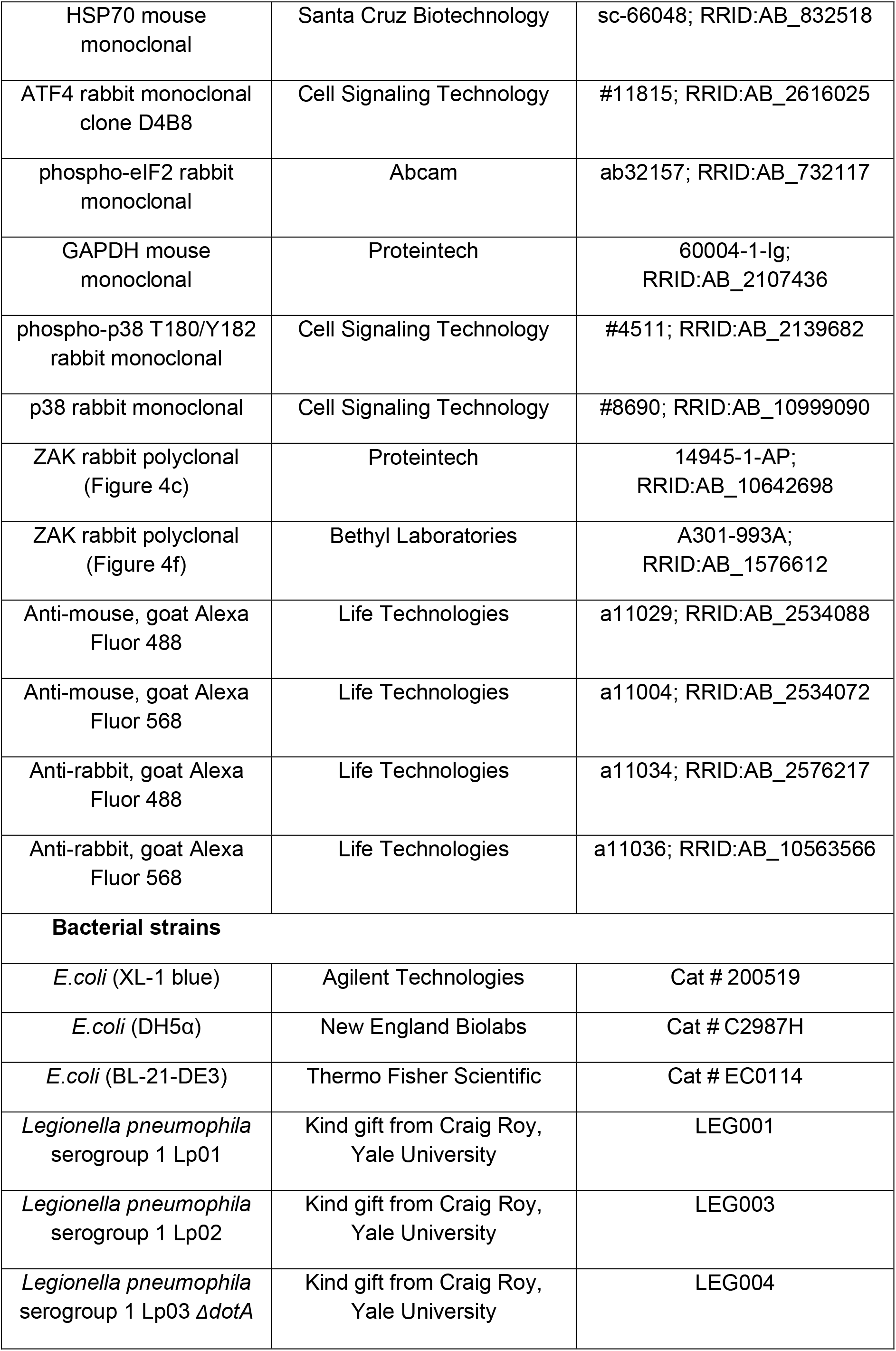
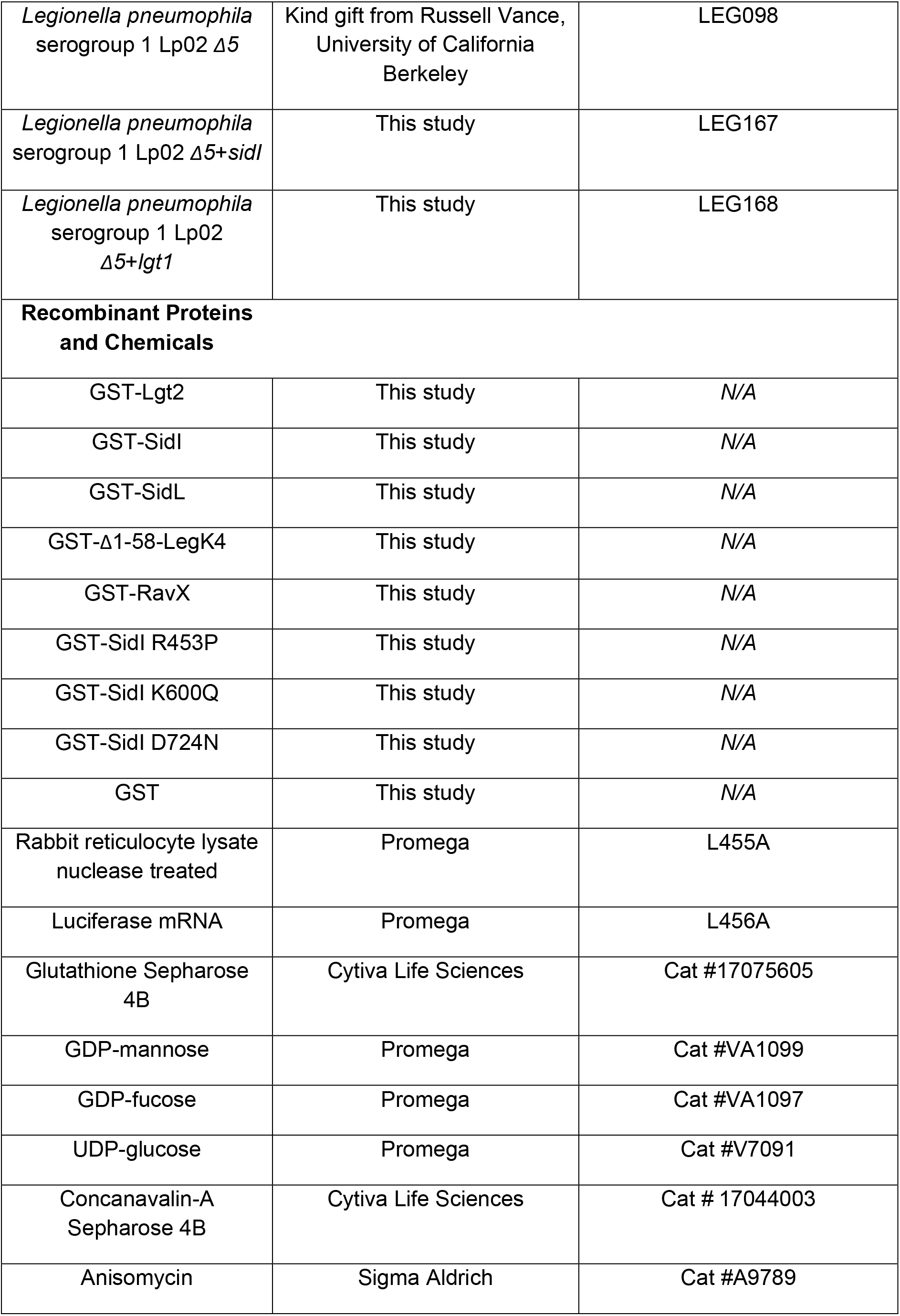
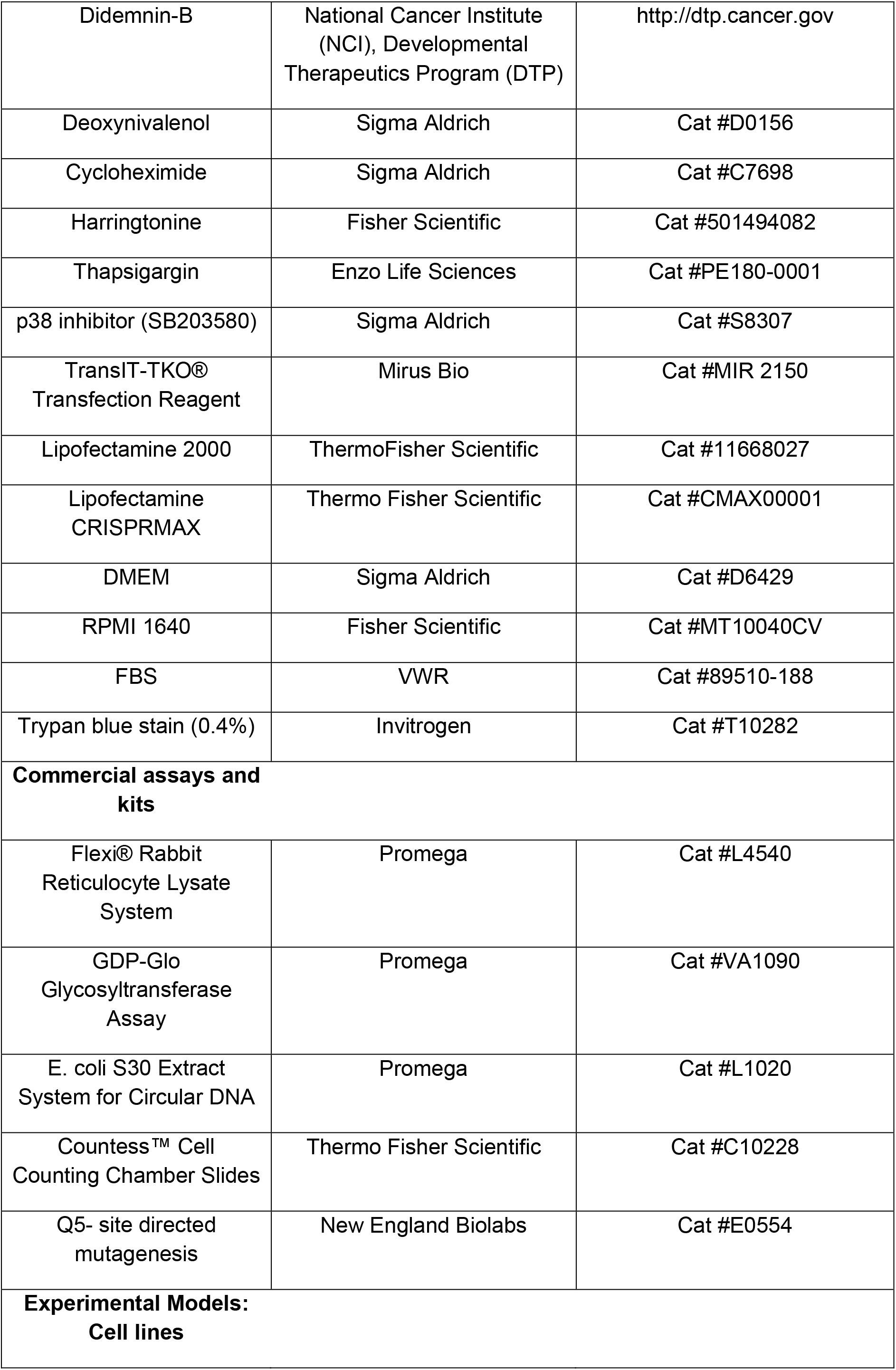
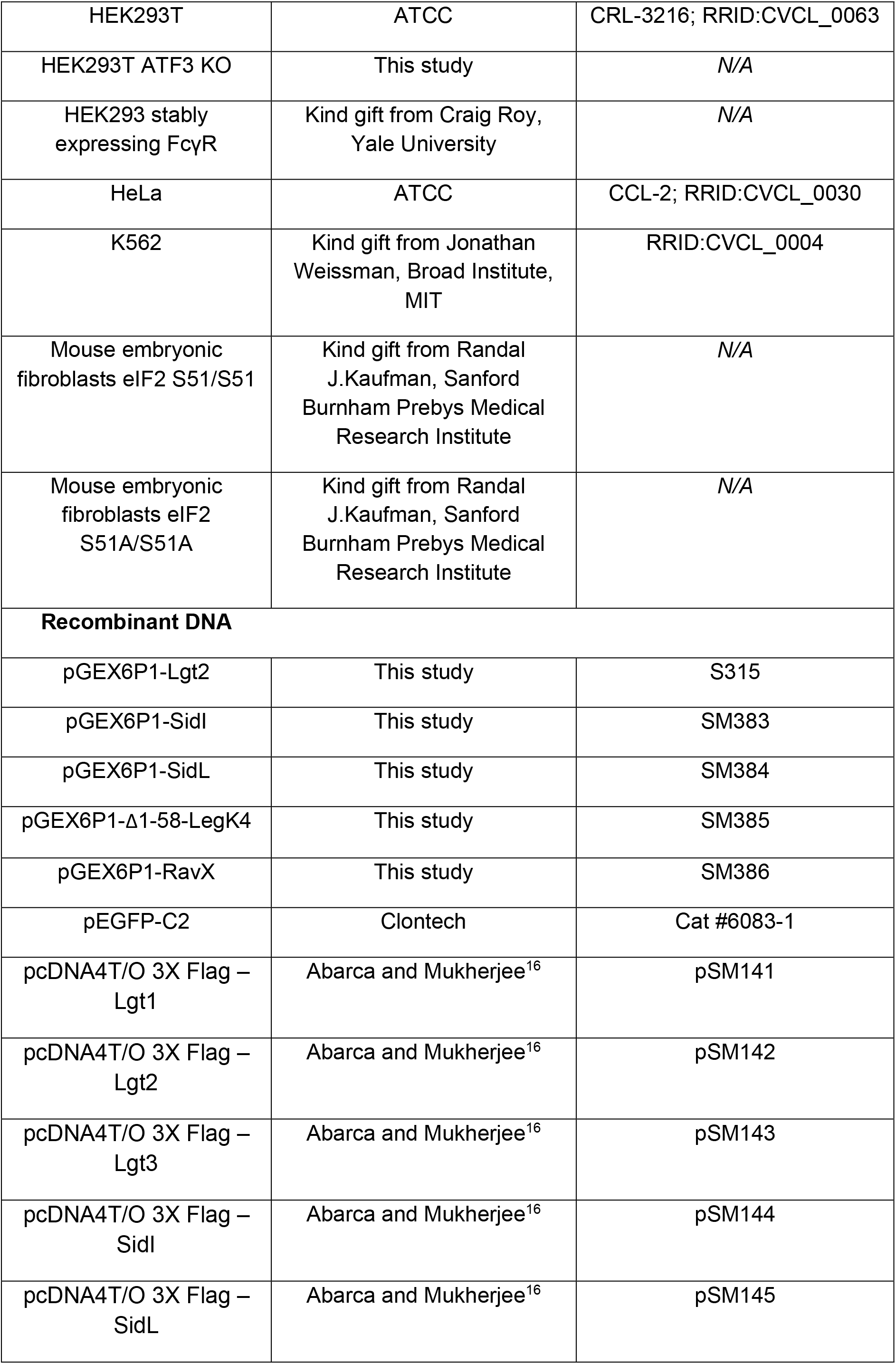
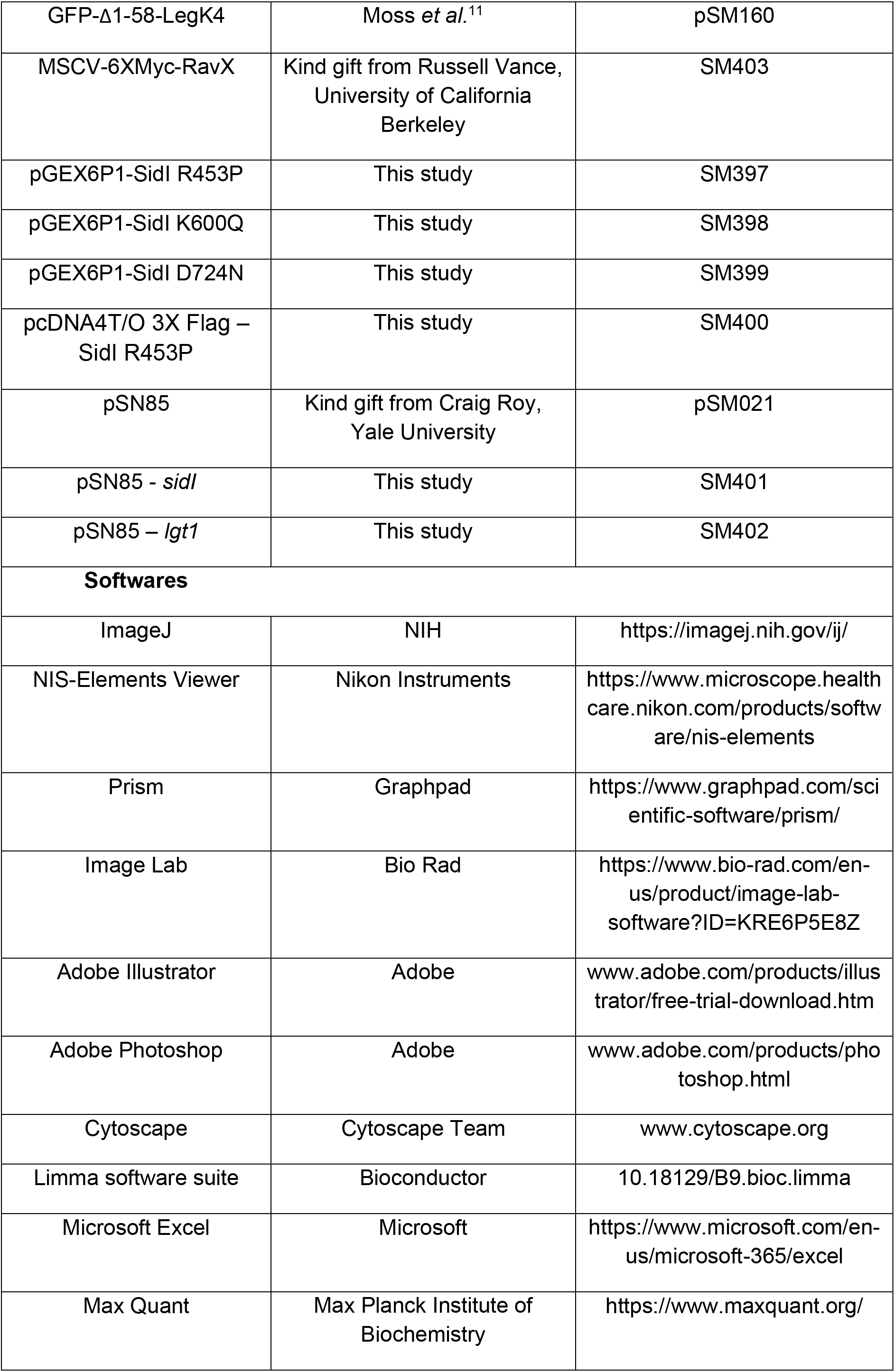
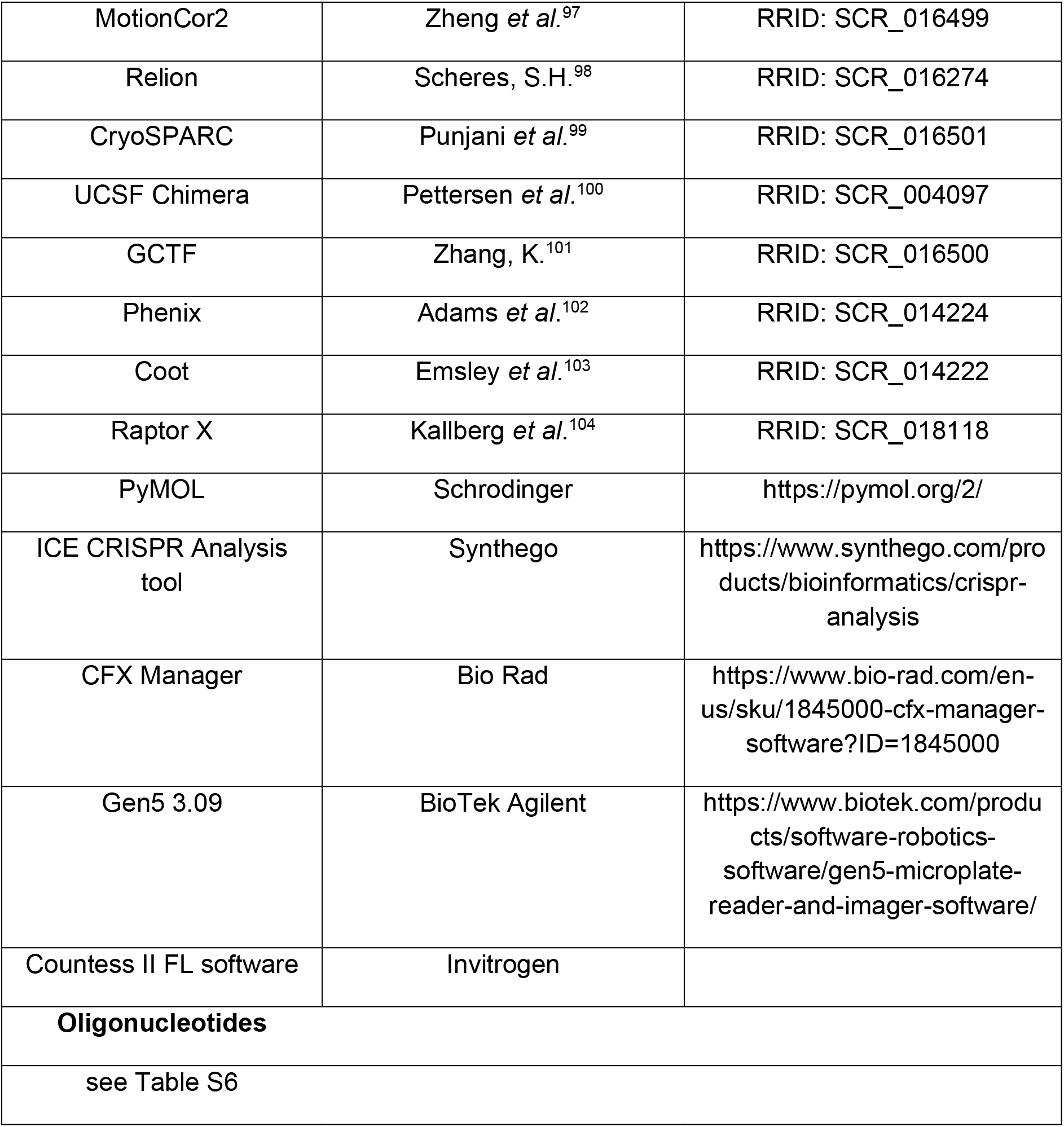

#### Cell lines and cell culture

HEK293T, HEK-293 FcγR, HeLa and MEF cells were cultured in DMEM-high glutamine media and K562 cells were cultured in RPMI-high glutamine media. Both DMEM and RPMI were supplemented with 10% fetal bovine serum (FBS) and the cells were maintained at 37 °C and 5% CO_2_. For the MEF cell cultures, DMEM media was further supplemented with penicillin-streptomycin antibiotic, sodium pyruvate and non-essential amino acid mixtures. HEK293T and HEK-293 FcγR cells were seeded in poly-L-lysine–treated plates. Cells were grown to ∼75 to ∼90% confluency for most experiments. Each cell line was routinely screened for mycoplasma contamination using the MycoScope™ PCR Mycoplasma Detection Kit (Genlantis Cat#MY01050).

#### Preparation of recombinant DNA

*L.p.* genes were amplified from *L.p.* (Lp01) genomic DNA by PCR and cloned into respective vectors. For mutagenesis, Q5 site directed mutagenesis kit (NEB) was used and the manufacturer’s instructions were followed.

#### Recombinant protein purification

*E. coli* BL21 (DE3) cultures expressing pGEX6P1 plasmids were first grown overnight at 37°C. These overnight cultures were then sub-cultured at 1:100 in LB supplemented with the appropriate antibiotics until OD_600_ values reached 0.6. This was followed by the induction of protein expression with 1 mM IPTG and the bacteria were cultured at 16°C overnight. The next day, bacterial cultures were centrifuged at 6000 rpm for 5 min at 4°C followed by incubation of pellets in bacterial lysis buffer [50 mM Tris pH 8, 100 mM NaCl, 1 mM ethylenediaminetetraacetic acid (EDTA), 200μg/mL lysozyme, 2 mM dithiothreitol (DTT), 10μg/mL DNase, and complete protease inhibitor] for 60 minutes on ice. The mixtures were then sonicated on ice [pulse on 30 seconds; pulse off 10 seconds – 30% amplitude for 5 minutes] and centrifuged at 15,000 g for 30 min at 4°C. The supernatant fractions were incubated with glutathione coupled sepharose beads overnight at 4°C with rotation. The beads were then transferred to a 20mL Maxi column with rubber stoppers and washed with 5 column volumes of wash buffer I (PBS, 0.05% Triton X-100, 1mM DTT), followed by wash buffer II (PBS, 0.05% Triton X-100, 0.5M NaCl, 1mM DTT). The GST-tagged proteins were then eluted with elution buffer (50 mM Tris pH 7.5, 150mM NaCl, 10 mM glutathione, 1mM DTT) and concentrated with Amicon Ultra centrifugal filters at 4°C with repeated mixing to prevent aggregate formation. Purified proteins were visualized by SDS-PAGE and Coomassie brilliant blue staining. Protein concentrations (in µM) were calculated by the Beer-Lambert equation A = εlc where A is the absorbance of the protein in solution at 280nM, ε is the calculated extinction coefficient of the protein based on its amino acid sequence, l is the light path length and c is the concentration of the protein in solution. The extinction coefficients used for calculations of GST-tagged proteins are as follows: GST-SidI (ext. coeff. = 125335); GST-Lgt2 (ext. coeff. = 113065); GST-SidL (ext. coeff. = 93920); GST-Δ1-58-LegK4 (ext. coeff. = 122270); GST-RavX (ext. coeff. = 62270). In certain cases, proteins were further purified by gel filtration chromatography using a Superdex 200 column attached to an AKTA-format FPLC (GE Healthcare) and running buffer containing 50mM Tris pH 7.5, 150mM NaCl and protease inhibitor cocktail. Elution fractions containing protein were analyzed by their A_280_ spectra, SDS-PAGE and Coomassie brilliant blue staining.

#### Cell free translation in rabbit reticulocyte lysates

For each treatment condition, luciferase mRNA (200ng) was translated for 60 minutes at 30°C in reaction mixtures containing nuclease treated rabbit reticulocyte lysates (12.5µL; ∼700µg of total protein content), amino acid mixture without leucine (10µM), amino acid mixture without methionine (10µM) and potassium chloride (70mM). By using two incomplete amino acid mixtures, a sufficient concentration of all amino acids is provided. After incubation, the samples were mixed in a 1:1 (v/v) ratio with luciferase assay reagent (Promega) in a 96-well white opaque plate and left at room temperature for 2 minutes. Luminescence readings were then obtained on a BioTek Cytation Imaging reader. The translation efficiency for each condition was calculated as a percentage of the raw luminescence values obtained when reaction mixtures were untreated or treated with GST alone. For the experiments is Figure 2o, rabbit reticulocyte lysates (12.5µL) were first incubated with 750pM of GST-SidI or 0.5µM of GST-SidI R453P for the indicated amounts of time at 30°C. After pre-treatment, amino acid mixtures, potassium chloride and 1µg of luciferase mRNA were added to the tubes and the samples were incubated for a further 60 minutes at 30°C and luminescence values were calculated as described above. Here, the translation efficiency for each condition was calculated as a percentage of the raw luminescence values obtained when the samples were not pre-treated with either GST-SidI or GST-SidI R453P (pre-treatment time 0’).

### SidI interactome analyses

#### Affinity-precipitation of interactors with glutathione sepharose coupled beads

HEK293T lysates were prepared using IP lysis buffer (150 mM NaCl, 25mM Tris-HCl pH=7.5, 1% Triton-X and protease cocktail inhibitor from Roche). Rabbit reticulocyte lysates were obtained from Promega. For each experimental replicate, the protein concentrations were quantified and 1mg of total protein lysate was incubated with 1µg/mL of purified GST or GST-SidI overnight at 4°C in a rotary mixer. 25µL of pre-washed glutathione sepahrose beads were added to the samples and the mixtures were incubated for a further 1 hour at 4°C in a rotary mixer. The beads were then washed 5x times with IP lysis buffer to remove any unspecific interactors. The bound interactors were eluted by boiling the beads in Laemmeli buffer containing β-mercaptoethanol and the eluates were run on a 4 – 15 % SDS-PAGE gel. The gels were silver stained (Pierce Silver Stain kit; Thermo Fischer Scientific) and the gel pieces containing prominent protein bands were excised and stored at −80°C for mass spectrometry.

#### Sample preparation

25mM NH_4_HCO_3_/50%Acetonitrile was added to the sample gel pieces after excisions and then dried by vacuum centrifugation. Some samples were reduced by the addition of 15mM TCEP in 25mM NH_4_HCO_3_, the supernatant removed, and then gel slices were alkylated by the addition of 50mM of chloroacetamide. Other samples were reduced by the addition of 10mM DTT in 25mM NH_4_HCO_3_, the supernatant was removed, and then alkylation proceeded by the addition of 55mM iodoacetamide in 25mM NH_4_HCO_3_. Gel slices were washed in 25mM NH_4_HCO_3_ and then dehydrated in 25mM NH_4_HCO_3_/50% Acetonitrile. They were then dried by vacuum centrifugation. A solution of 25mM NH_4_HCO_3_ containing 4ug/mL trypsin was then added to the gel slices and digested overnight at 37°C overnight. The resulting peptides in solution were transferred to a new tube and desalted on C18 spin columns. The eluting peptides were dried by vacuum centrifugation and then resuspended in 0.1% formic acid.

#### Mass spectrometry data acquisition and analysis

Samples were injected onto a column (360 µm O.D. x 75 µm I.D., New Objective) packed with 25cm of 1.8 µm Reprosil C18 particles with (Dr. Maisch). Peptides were separated by a reversed-phase gradient using an Easy-nLC 1200 (Thermo Scientific) with mobile phase A consisting of 0.1% formic acid, and mobile phase B consisting of 80% acetonitrile in 0.1% formic acid. Eluting peptides were directly injected into either an Q-Exactive Plus Orbitrap mass spectrometer (Thermo Scientific) or an Exploris 480 Orbitrap mass spectrometer (Thermo Scientific) equipped with a nanospray flex source and operating in positive ion mode. System for both instruments was monitored with QCloud2.On the Q-Exactive Plus mass spectrometer, the total acquisition time was 60 min, MS1 detection was performed in profile mode in the Orbitrap with 70,000 resolution, 300-1500 m/z scan range, 250 ms maximum injection, and an AGC target of 1e6. MS2 fragmentation was performed on charge states greater than 2, with automatic dynamic exclusion, an expected 15s peak width, data collection was in the Oribtrap in centroid mode with 17,500 resolution, 60ms maximum injection time, 5e4 AGC target, 200-2000 m/z scan range, and a 2.0 m/z isolation window and a 26% normalized HCD collision energy. On the Exploris 480 mass spectrometer, the total acquisition time was 60 min, MS1 detection was performed in profile mode in the Orbitrap with 120,000 resolution, 350-1250 m/z scan range, maximum injection time was set to “auto”, and a normalized AGC target of 100%. MS2 fragmentation was performed on charge states 2-6, with 30s dynamic exclusion after 2 occurrences and a +/- 10 ppm window, data collection was in the Oribtrap in profile mode with 15,000 resolution, maximum injection time set to “auto”, 200% normalized AGC target, and a 1.3 m/z isolation window and a 30% normalized HCD collision energy. All raw MS data were searched with MaxQuant (version 2.0.3.0) against the human proteome (Uniprot canonical protein sequences downloaded June 21, 2021) or against the rabbit proteome (*Oryctolagus cuniculus,* downloaded April 6, 2022). Peptides were filtered to 1% false discovery rate in MaxQuant. MaxQuant default parameters were used with the exception that label-free quantification was turned on with match between runs set to 0.7 min. Mass spectrometry data files (raw and search results) have been deposited to the ProteomeXchange Consortium (http://proteomecentral.proteomexchange.org) via the PRIDE partner repository. iBAQ intensity values were used for estimating the relative enrichment of GST-SidI interacting proteins compared to GST-only control samples.

#### Purification of 80S ribosomes

Monosomes were purified from K562 cells. A 5ml pellet of K562 cells was resuspended in 3 mL of Lysis Buffer (25 mM HEPES, pH 7.4, 300 mM KOAc, 10 mM MgOAc_2_, 1 mM DTT, 0.5% NP-40, complete EDTA-free protease inhibitor cocktail (Roche)), and lysed by short intervals of vortexing (10 min). Cell debris were removed by centrifugation for 10 min at 12,000 × g at 4°C and concentration of KOAc was adjusted to 500 mM. Lysate was loaded on a (3:1 v/v) 1M sucrose cushion (25 mM HEPES pH 7.5, 150 mM KOAc, 10 mM MgOAc_2_, 1 mM DTT, 1 M sucrose and .5 mM PMSF) and centrifuged at 115,800 x g for 15 hrs. The crude ribosomal pellet obtained after the spin was resuspended in buffer (25 mM HEPES pH 7.5, 150 mM KOAc, 10 mM MgOAc, 1 mM DTT, .5 mM PMSF, 1mM EDTA, 1x protease inhibitor cocktail, .2U/mL RNase Inhibitor) and treated with 1 mM puromycin for 15 mins on ice and then 15 mins at 25 C. The 80S monosome fraction and 40S and 60S subunits were then separated by loading onto a 15–30% continuous sucrose gradient (25 mM HEPES, pH 7.4, 500 mM KOAc, 10 mM MgOAc, 1 mM DTT, 1x protease inhibitor cocktail) and centrifuged at 49,500 x g for 18 hrs using a SW40 rotor. Monosomes were pelleted from the suitable fraction by centrifugation at 25,000 rpm for 14.5 hr using a TLA110 rotor. The pellets were suspended in storage buffer (20 mM HEPES, pH7.5, 100 mM KOAc, 10 mM MgOAc_2_, 1 mM DTT) and stored flash frozen in 10 µl aliquots.

#### *In vitro* ribosome binding assay

25µL of glutathione sepharose beads were pre-immobilized with 250nM of GST, GST-SidI or GST-SidI R453P in 100µL of ribosome binding buffer (50mM HEPES, 100mM KOAc, 5mM MgOAc_2_, 0.1% NP-40) at 4°C for 1 hour and then washed 3x times with ribosome binding buffer. GST/GST-SidI/GST-SidI R453P immobilized on beads were then incubated with 50nM of purified 80S ribosome fractions in 100µL of ribosome binding buffer at 4°C for 3 hours. The beads were eluted with 30µL of ribosome binding buffer containing 10mM glutathione. The eluates were then subjected to SDS PAGE and immunoblotting with antibodies against 40S and 60S subunit proteins.

#### GST-tag based affinity-precipitation of proteins from HEK293T lysates

HEK293T lysates were prepared using IP lysis buffer (150 mM NaCl, 25mM Tris-HCl pH=7.5, 1% Triton-X, 10mM Na3VO4, 40mM β-glycerophosphate, 10mM NaF and protease cocktail inhibitor from Roche). 1mg of total protein lysate was incubated with 1µg/mL of purified GST or GST-SidI overnight at 4°C in a rotary mixer. 25µL of pre-washed glutathione sepahrose beads were added to the samples and the mixtures were incubated for a further 1 hour at 4°C in a rotary mixer. The beads were then washed 5x times with IP lysis buffer to remove any unspecific interactors. The bound interactors were eluted by boiling the beads in Laemmeli buffer containing β-mercaptoethanol and the eluates were subjected to SDS-PAGE and immunoblotting with antibodies.

### Cryo-EM structure determination of SidI

#### Sample preparation for cryo-electron microscopy

For grid freezing, a 3 μl aliquot of 3 μM GST-SidI was applied onto the Quantifoil R 1.2/1/3 400 mesh Gold grid and waited for 30 s. A 0.5 μl aliquot of 0.1% Nonidet P-40 substitute was added immediately before blotting. The entire blotting procedure was performed using Vitrobot (FEI) at 10 °C and 100% humidity.

#### Electron microscopy data collection

Cryo-EM data was collected on a Titan Krios transmission electron microscope operating at 300 keV. Micrographs were acquired using a Gatan K3 direct electron detector. The total dose was 60 e^-^/ Å^2^, and 60 frames were recorded during a 5.7 s exposure. Data was collected at 81,000 x nominal magnification (0.844 Å/pixel at the specimen level), with a nominal defocus range of −1.0 to −2.0 μm.

#### Cryo-EM image processing

The micrograph frames were aligned using MotionCor2 ^97^. The contrast transfer function (CTF) parameters were estimated with GCTF (PMID: 26592709). Particles were picked using Gautomatch (developed by Kai Zhang, https://www2.mrc-lmb.cam.ac.uk/research/locally-developed-software/zhang-software/#gauto) without a template. Particles were extracted using a 64-pixel box size and classified in 2D in Relion^98^. Classes that showed clear protein features were selected. Selected particles were then split into three subsets for efficient data processing. Each subclass was subjected to 3D classification (k= 3) and the best was selected. For one of these subsets, particles in the best 3D class were refined to 7.0 Å resolution. These particles were then extracted at a smaller pixel size (0.844 Å) and refined to yield a 4.0 Å reconstruction, which was used as a reference structure for the following steps. Upon obtaining the reference structure, particles from all three subsets were imported to cryoSPARC^99^ for homogeneous refinement (using the 4.0 Å reconstruction in Relion as a reference structure) and heterogeneous refinement. The resulting particles from each subset were combined to go through one more round of heterogeneous refinement followed by nonuniform refinement, which results in the final reconstruction at 3.1 Å resolution, consisting of 749 k particles.

#### Atomic model building, refinement, and visualization

To build the atomic model for SidI, we first performed a structure prediction analysis using RaptorX^104^ and found that part of SidI shares structural homology with a few sugar-nucleotides. We then used the crystal structure of the *B. subtilis* BshA (PDB ID:5D00) as a template to build the C-terminal part of SidI (amino acids 355-865). The predicted structure of SidI C-terminal domain was manually docked into the EM density of SidI and was used as a template for model building. The models were then manually adjusted in Coot^103^. Having built the C-terminal domain, we continued to build the rest of SidI *de novo*. The complete model was then refined in Phenix^102^ using global minimization, secondary structure restraints, Ramachandran restraints, and local grid search. Then iterative cycles of manual rebuilding in Coot and Phenix were performed. The final model statistics were tabulated using Molprobity^105^. Molecular graphics and analyses were performed with the UCSF Chimera package^100^. UCSF Chimera is developed by the Resource for Biocomputing, Visualization, and Informatics and is supported by NIGMS P41-GM103311. The atomic model will be deposited into PDB and the EM map will be deposited into EMDB.

#### Nucleotide-sugar hydrolysis assays

Recombinantly purified proteins at the indicated concentrations were incubated with nucleotide-sugar precursors (50µM or 100µM) in glycosyl transferase buffer (50mM Tris pH = 7.4, 5mM MnCl_2_ for assays with UDP-glucose and GDP-mannose; 25mM HEPES pH = 7.5, 2.5mM MnCl_2_ for assays with GDP-fucose). For experiments in Figure S5a, crude ribosome extracts were added to the reaction mixtures. All samples were then incubated at 37°C for 60 minutes. After incubation, the samples were mixed in a 1:1 (v/v) ratio with nucleotide detection reagent (Promega) in a 96-well white opaque plate and left at room temperature in the dark for another 60 minutes. The addition of the nucleotide detection reagent quenches the glycosyl transferase reaction. Luminescence readings were then obtained on a BioTek Cytation Imaging reader. To measure nucleotide release for each experiment, raw luminescence values obtained by treating serial dilutions of pure UDP or GDP (Promega) with the nucleotide detection reagent for 60 minutes were used to plot a standard concentration curve. The slope of the standard curve was then used to calculate the amount of nucleotide hydrolysed under different treatment conditions.

#### Preparation of crude ribosome extracts

Eight 10 cm dishes of HEK293T cells at 80% confluency were used. After wash with cold PBS twice, cells were collected and resuspended in 1.5 mL cytosol buffer (50 mM HEPES, pH 7.4, 100 mM KOAc, 5 mM Mg(OAc)2, 0.01% digitonin, 40 U/ml Ribonuclease inhibitor (Promega, N2511), 1 mM DTT, and protease inhibitor cocktail), and then disrupted mechanically by passage through a pre-chilled 26G needle using a 10 mL syringe. Cellular debris were cleared by centrifugation at 4°C for 15 min at 12000rpm. The supernatant was collected and the concentrations of KOAc and MgAc2 in the supernatant were increased to 500 mM and 15 mM, respectively. NP-40 detergent was also added to a final concentration of 0.2% to disrupt ribosome-associated proteins. 300µl of sample was then layered over the 1mL sucrose cushion (20 mM HEPES pH 7.4, 500 mM KOAc, 15 mM MgAc2, 0.1 mM EDTA pH 7.4, 1 M sucrose) and centrifuged at 100,000 RPM for 60 minutes at 4°C in a TLA100.3 rotor (Beckman Coulter, 349490) with 3.5 mL polycarbonate tubes (Beckman Coulter, 349622). The supernatant was removed carefully with an aspirator, and the pellets were washed with 100µl ribosome binding buffer (50 mM HEPES, pH 7.4, 100 mM KOAc, 5 mM Mg(OAc)2) [add on top carefully and aspirate; do not disrupt pellet] and resuspended with 30 µl ribosome binding buffer. Before measuring the concentration of ribosomes by A260 with NanoDrop, ribosomes from different tubes were pooled. Ribosomes were aliquoted and flash-frozen in liquid nitrogen and stored at −80°C. The concentration of ribosomes in the extract was calculated by Beer-Lambert equation A = εlc where A is the absorbance of the ribosomes at A260, ε is the extinction coefficient of a monosome (4.2 x 10^7^), l is the light path length and c is the concentration of the ribosomes in solution.

#### *In vitro* mannosylation and concanavalin A pulldown

1.4µM of recombinantly purified GST, GST-SidI or GST-SidI R453P were incubated with or without 1mM of GDP-mannose and 30nM of ribosome extracts in glycosyl transferase buffer (50mM Tris pH = 7.4, 5mM MnCl_2_) at 37°C for 60 minutes. 25µL of concanavalin-A sepharose beads were washed with ribosome binding buffer containing 0.2% NP-40 and the washed beads were added to the samples and incubated at 4°C for 3 hours in a rotary mixer. The beads were then washed 3x times with ribosome binding buffer and bound mannosylated molecules were eluted with ribosome binding buffer containing 200mM methyl-α-D-mannopyranoside (Sigma-Aldrich, M6882). The eluted fractions were then subjected to SDS-PAGE and immunoblotting with antibodies.

#### Bacterial strains and infection protocol

All *L.p.* strains were grown on charcoal yeast extract (CYE) agar plates supplemented with iron (FeNO_3_ 0.135g/10mL) and cysteine (0.4g/10mL). Chloramphenicol (10µg/mL), IPTG (0.1mM), and thymidine (100µg/mL) were added to CYE agar plates as needed. For infection experiments, primary patches of *L.p.* were grown for 2 days at 37°C on CYE plates. Single colonies of bacteria were picked and grown as heavy patches for 2 further days on CYE plates at 37°C. Heavy patches were then harvested, diluted in AYE broth supplemented with the required chemicals (as appropriate) and grown overnight at 37°C with shaking (220rpm) until the OD_600_ = ∼3. HEK293 FcγR cells were infected at a multiplicity of infection (MOI) of 50 or 5. For opsonization, *L.p.* bacteria were diluted in complete media and mixed with a *Legionella* polyclonal antibody (Invitrogen, Cat #PA1-7227) at 1:1,000 dilution. The mixtures were then incubated for 20 minutes at room temperature in a rotary mixer. Immediately after addition of the opsonized *L.p.*mixtures, cells were centrifuged for 5 min at 1,000 rpm to facilitate spin-fection. Cells were then incubated at 37 °C for 60 minutes to allow bacterial uptake via the FcγR. After 60 minutes, the cells were washed with 1x PBS to remove extracellular bacteria and replenished with DMEM supplemented with 10% FBS. Infected cells were then cultured at 37 °C until time of harvest.

#### Transfection

HEK293T cells were transfected with plasmid vectors using the TransIT-TKO Transfection Reagent. siRNA treatments in HeLa and HEK293 FcγR cells were carried out using a pool of siRNAs and Lipofectamine 2000. Expression or knockdown efficiencies (>85%) were checked after every experiment by immunoblotting. Each individual siRNA in the pool was also tested for knockdown efficiency with similar results. A list of siRNA sequences used in this study can be found in Table S6.

#### RNA extraction and quantitative real time PCR

RNA isolation was performed using the Direct-zol™ RNA Miniprep Plus kit (Zymo Research) according to the manufacturers protocols. The yield and the integrity of RNA were determined by spectrophotometer NanoDrop 2000c For qRT PCR analyses, 1µg of RNA was converted to cDNA using the QuantiTect Reverse Transcription Kit (Qiagen). Quantitative real time PCR reactions were performed with cDNA using iTaq Universal SYBR Green supermix (Bio-Rad) in 384 well plates and 500nM of forward and reverse primers on a Bio-Rad CFX384 Thermal cycler. The fold changes in the relative quantifications were calculated according the ΔΔCt method. See table S6 for a list of qRT-PCR primers.

#### Puromycin-pulse chase assays

Cells were treated with 2µg/mL puromycin for 10 minutes followed by puromycin wash out with PBS. Cells were then chased with complete medium for 50 minutes, washed with PBS and lysed with RIPA buffer. The lysates were then subjected to immunoblotting. For cells infected with *L.p.*, chloramphenicol (25 µg/mL) was added 15 minutes prior to puromycin addition to inhibit bacterial translation.

#### Cell extract preparation and immunoblotting

Cells were washed three times with PBS and collected immediately at 4°C in RIPA lysis buffer containing phosphatase inhibitors (1% Triton X-100, 20mM Tris-HCl, pH 8.0, 0.1% SDS, 0.05% sodium deoxycholate, 150mM NaCl, 10mM Na3VO4, 40mM β-glycerophosphate, 10mM NaF) and complete protease inhibitors (5×; Roche). The detergent soluble supernatant fractions were immediately processed for SDS– PAGE and immunoblotting with antibodies.

#### Polysome profiling

HEK293T cells grown in 10cm plates were transfected with constructs expressing GFP, FLAG-SidI or FLAG-SidI R453P for 24 hours. Cells were then treated with cycloheximide (100 µg/mL) by adding it to the media for 5 minutes before harvesting. Cells were washed with ∼5ml ice cold PBS + cycloheximide and spun at 16,000g for 1 minute at 4°C. The pellet fraction was resuspended in lysis buffer (10 mM HEPES pH = 7.9, 1.5 mM MgCl_2_, 10 mM KCl, 0.5 mM DTT, 1% Triton X-100, 100 ug/ml cycloheximide). The cells were incubated for 10 minutes on ice to allow swelling. Cells were then triturated ten times through a pre-chilled 26G needle and the debris were spun at 1500 g for 5 minutes at 4°C to pellet nuclei. The supernatant fraction of the lysate was then flash frozen and stored at −80°C if required. 200 µl of lysate was layered on top of a 10% to 50% sucrose gradient containing 100 mM KCl, 20 mM HEPES (free acid) pH = 7.6, 5 mM MgCl_2_, 1mM DTT and 100 µg/mL cycloheximide and spun at 36,000 rpm for 2 hours at 4°C in a SW-41 swinging bucket rotor. The tubes were then punctured at the bottom with a 16G needle connected to a syringe pump and ribosome populations across the gradient were analysed by measuring absorbance values at 254nm using a spectrophotometer.

#### RNA sequencing

After RNA extraction, mRNA was enriched by using Oligo dT beads from the KAPA mRNA Capture Kit (KK8581). Subsequent library preparation steps of fragmentation, adapter ligation and cDNA synthesis were done on the enriched mRNA using the KAPA RNA HyperPrep kit (KK8540). Truncated universal stub adapters were used for ligation, and indexed primers were used during PCR amplification to complete the adapters and to enrich the libraries for adapter-ligated fragments. Samples were checked for quality on an AATI (now Agilent) Fragment Analyzer. Samples were then transferred to the Vincent J. Coates Genomics Sequencing Laboratory (GSL), a QB3-Berkeley Core Research Facility at UC Berkeley, where Illumina sequencing library molarity was measured with quantitative PCR with the Kapa Biosystems Illumina Quant qPCR Kits on a BioRad CFX Connect thermal cycler. Libraries were then pooled evenly by molarity and sequenced on an Illumina NovaSeq6000 150PE S4 flowcell, generating 25M read pairs per sample. Raw sequencing data was converted into fastq format, sample specific files using the Illumina bcl2fastq2 software on the sequencing centers local linux server. For data analysis, kallisto, a program for quantifying abundances of transcripts from RNA-Seq data was used. Release 94 of the human reference transcriptome GRCh38 was downloaded from ENSEMBL on 7/20/2019. Relative abundances (reported as transcripts per million TPM values^106^ and estimated counts (est_counts) of each transcript in each sample were estimated by alignment free comparison of k-mers between the pre-processed reads and the transcriptome using KALLISTO version 0.43.0^107^. Further analysis was restricted to transcripts with TPM >= 10 in at least one sample. Differentially expressed genes were identified by comparing replicate means for contrasts of interest using LIMMA version 3.30.8^108,109^. Genes were considered significantly differentially expressed if they were statistically significant (at 5% FDR) with an effect size of at least 2x (absolute log2 fold change >= 1) for a given contrast. Principal components analysis (PCA)^110^ was carried out on the matrix of sample vs. transcript log (TPMs) by mean centering the values for each gene and extracting principal components by singular value decomposition as implemented in Numpy 1.12.1^111^. Gene set enrichment analyses on the ranked list of genes was performed as previously described^52^. The raw sequencing data has been submitted to the Gene Expression Omnibus (GEO) server.

#### Indirect immunofluorescence and confocal microscopy

Indirect immunofluorescence (IF) was performed as follows: cells grown on coverslips were washed in phosphate-buffered saline (PBS) and fixed in freshly prepared PBS supplemented with 4% paraformaldehyde (Electron Microscopy Sciences, Hatfield, USA) for 30 min at room temperature (RT). Cells were permeabilized and blocked in blocking buffer (0.05% saponin in 0.5% BSA) for 30 min at room temperature. Primary antibodies were incubated for 1 hour at RT or overnight at 4°C in blocking buffer. Cells were subsequently labelled with appropriate Alexa 488/568-tagged fluorescent-conjugated secondary antibodies. Images were acquired using an inverted Nikon Eclipse Ti-E spinning disk confocal microscope equipped with a Prime 95B 25mm CMOS camera (Photometrics) camera. Identical settings for each channel were maintained throughout single experiments. For figure presentation only, the images were channel-separated, and each channel was merged after correcting for contrast using Image-J (NIH) or Adobe Photoshop CS3 (Adobe Systems).

#### CRISPR-Cas9 mediated knockout of ATF3

Ribonucleoprotein complexes containing 3.9 picomoles of ATF3 sgRNA guides and 3 picomoles of purified recombinant Cas9 protein (Synthego) were transfected into HEK293T cells using Lipofectamine CRISPRmax (Invitrogen). 72 hours post transfection, cells were trypsinized and single cell dilutions were plated into 2 x 96 well plates (∼1 cell/well) and allowed to grow in complete media for 7 days to facilitate clonal expansion. Viable clones were then trypsinized and seeded into replica 24 well plates and allowed to grow. A primary screen for CRISPR-Cas9 knockout efficiency was performed by lysing the clonal cell populations and subjecting the lysates to SDS-PAGE and immunoblotting with antibodies against ATF3. Clonal populations positive for the loss of ATF3 protein were expanded and their genomic DNA was isolated. Genotyping of clones was performed by PCR-amplifying exon 2 of the *ATF3* gene using the primers F: tggccacatttgcacataggcaca and R: acccagcaaccccatccctacc. The amplicons were sequenced and analyzed using Synthego’s Inference of CRISPR Edits (ICE) tool. The ATF3 knockout HEK293T cell line clone D10 presented with indels in exon 2 of the *ATF3* gene and a complete loss of ATF3 protein as determined by immunoblotting with antibodies against ATF3.

#### Measurement of cell viability

After indicated treatments, media was aspirated, and the cells were carefully scraped and diluted in fresh media. Media containing cells was then mixed in a 1:1 (v/v) ratio with 0.4% trypan blue, a dye that only stains non-viable cells with compromised membranes. 10µL of this mixture was mounted onto Countess cell counting chambers (Invitrogen) and the number of viable cells was determined by automatic counting using a Cell Countess FL II machine programmed with a plugin to highlight stained (non-viable) and unstained (viable) cells. The percentage of viable cells was calculated by normalizing total viable cell numbers to the counts obtained under control conditions.

#### Cell free transcription and translation in *E.coli* S30 extracts

For each treatment condition, 1µg of luciferase DNA (pBEST*luc* Promega L485A) was transcribed and translated for 60 minutes at 37°C in reaction mixtures containing *E.coli* S30 extract (7.5µL), amino acid mixture without leucine (10µM), amino acid mixture without methionine (10µM) and *E.coli* S30 premix. 100µM of GDP-mannose was added to the reaction mixtures when indicated. After incubation, the samples were mixed in a 1:1 (v/v) ratio with luciferase assay reagent (Promega) in a 96-well white opaque plate and left at room temperature for 2 minutes. Luminescence readings were then obtained on a BioTek Cytation Imaging reader. The translation efficiency for each condition was calculated as a percentage of the raw luminescence values obtained when reaction mixtures were untreated.

#### Inferred kinase activity prediction from phosphoproteomics datasets

Phosphoproteomics datasets generated by Noack *et al.*^62-preprint^ were utilized for calculating kinase activity scores. To detect phosphorylation sites on host proteins during *L.p.* infections, HEK293 FcγR were infected for 1 hour with the *L.p.* WT strain or the *ΔdotA* strain at a MOI = 100. After infection, cells were washed with ice-cold PBS, scraped, and pelleted by centrifugation at 1500 rpm for 5 minutes at 4°C. Cell pellets were then lysed by probe sonication in three pulses of 20% amplitude for 15 seconds on ice in lysis buffer [8M urea, 150 mM NaCl, 100 mM ammonium bicarbonate, pH = 8; protease inhibitor and phosphatase inhibitor cocktails]. In order to remove insoluble precipitates, lysates were centrifuged at 16,000 g at 4°C for 30 min. 6 mg of protein isolated from each infection condition was reduced with 4 mM tris(2-carboxyethyl) phosphine for 30 min at room temperature and alkylated with 10 mM iodoacetamide for 30 min at room temperature in the dark. Remaining alkylated agent was quenched with 10 mM 1,4-dithiothreitol for 30 min at room temperature in the dark. The samples were diluted with three starting volumes of 100 mM ammonium bicarbonate, pH 8.0, to reduce the urea concentration to 2 M. Samples were incubated with 50 μg of sequencing grade modified trypsin (Promega) and incubated at room temperature with rotation for 18 hours. The sample pH was reduced to approximately 2.0 by the addition of 10% trifluoroacetic acid (TFA) to a final concentration of 0.3% trifluoroacetic acid. Insoluble material was removed by centrifugation at 16,000 g for 10 min. Peptides were desalted using SepPak C18 solid-phase extraction cartridges (Waters). The columns were activated with 1 ml of 80% acetonitrile (ACN), 0.1% TFA, and equilibrated 3 times with 1 ml of 0.1% TFA. Peptide samples were applied to the columns, and the columns were washed 3 times with 1 ml of 0.1% TFA. Peptides were eluted with 1.2 ml of 50% ACN, 0.25% formic acid. Phosphopeptides were enriched by immobilized metal affinity chromatography. In-house prepared iron nitriloacetic acid coupled beads (Fe-NTA resin) were loaded into pre-equilibrated silica C18 microspin columns. Dried peptide samples were resuspended in a solution of 200 μl 75% ACN and 0.15% TFA. Peptide samples were mixed twice with the Fe-NTA resin, allowing the peptides to incubate for 2 minutes between each mixing step. The resin was rinsed four times with 200 μl of 80% ACN, 0.1% TFA. In order to equilibrate the columns, 200 μl of 0.5% formic acid was applied twice to the resin and columns. Peptides were then eluted from the resin onto the C18 column by mixing and incubating the Fe-NTA resin with 200 μl of 500 mM potassium phosphate, pH = 7.0 for 2 minutes. Peptides bound to the C18 column were washed three times with 200 μl of 0.5% formic acid. The C18 columns were removed from the vacuum manifold and the bound peptides were eluted by centrifugation at 1000g with 75 μl of 50% ACN, 0.25% formic acid. Peptides were dried with a centrifugal adaptor and stored at - 20°C until analysis by liquid chromatography and mass spectrometry. Samples were separated by a reverse phase gradient over a nanoflow column (360 μm O.D. x 75 μm I.D.) packed with 25 cm of 1.8 μm Reprosil C18 particles and directly injected into an Orbitrap Fusion Lumos Tribrid Mass Spectrometer (Thermo). Total acquisition time was 100 minutes. Raw MS data were searched with MaxQuant against the human proteome (UniProt canonical protein sequences downloaded January 11, 2016). Kinase activity scores were calculated based on log_2_fold change values of all detected phosphosites (12751 phosphosites from 3113 proteins) during WT *L.p.* infection in comparison with *ΔdotA* infection using the bioinformatic tool PhosFate (http://phosfate.com)^112^.

#### Bioinformatic mining of ATF3 ChIP-seq tracks

The human hg38 build on the UCSC genome browser was mined. The gene tracks represent the NCBI Ref Seq with the exons for each gene highlighted. The histone H3 lysine 27 acetylation tracks contain information relevant to the regulation of transcription from the ENCODE Project^113,114^. The Layered H3K27Ac tracks show where modification of histone proteins is suggestive of enhancer and, to a lesser extent, other regulatory activity. The actual enhancers are typically just a small portion of the area marked by these histone modifications. These tracks are a cumulative representation of data collected from 7 different cell lines (GM12878, H1-hESC, HSMM, HUVEC, K562, NHEK and NHLF) and use a transparent overlay method of displaying data from a number of cell lines in the same vertical space. Each of the cell lines in this track is associated with a particular colour. The transcription factor ChIP-seq peaks for ATF3 were mined from ENCODE 3 and were generated by the ENCODE Transcription Factor ChIP-seq Processing Pipeline. Methods documentation and full metadata for each track can be found at the ENCODE project portal, using the ENCODE file accession (K562 = ENCFF937OKC; A549 = ENCFF851UTY; HepG2 = ENCFF137OEY; Liver cells = ENCFF146URA). The display for this track shows site location with the point-source of the peak marked with a red coloured vertical bar and the level of enrichment at the site indicated by the darkness of the item. The score values were computed at UCSC based on signal values assigned by the ENCODE pipeline. The input signal values were multiplied by a normalization factor calculated as the ratio of the maximum score value (1000) to the signal value at 1 standard deviation from the mean, with values exceeding 1000 capped at 1000.

#### Quantification and Statistical Analysis

The number of replicates and statistical parameters for each experiment are listed in the corresponding figure legends. Where indicated, p-values were calculated with a two-tailed students t-test. For the densitometric analysis of western blots, protein bands on membranes were captured on a BioRad Chemidoc Touch apparatus. The exposure times were varied to obtain appropriate signal intensities of the protein bands. The intensities of each of the bands were quantitated using the Image-J gel analysis tool.

---

**Figure S1.**
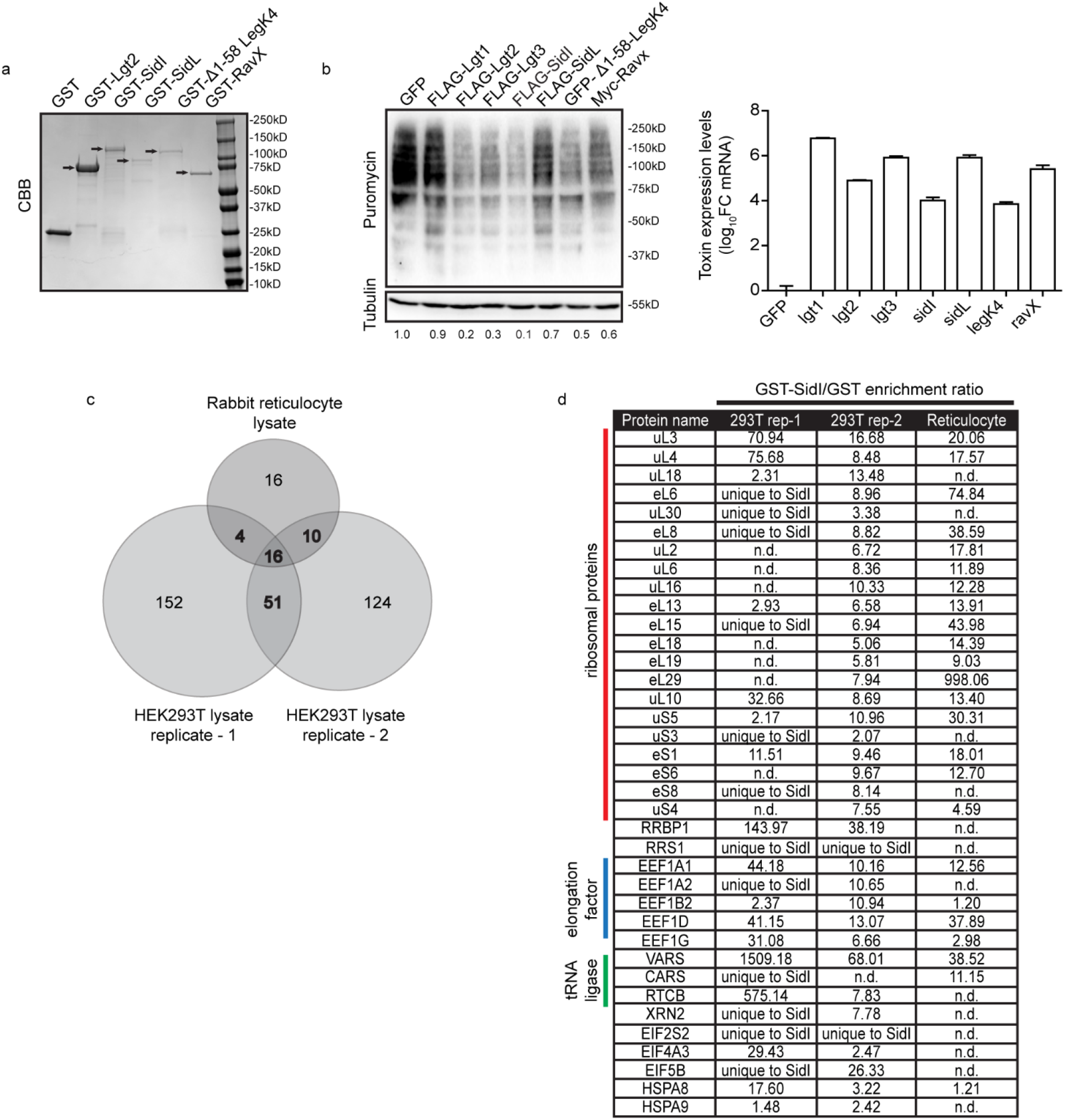
Mass spectrometry analyses of SidI interacting partners highlights the translation machinery as targets of SidI. (a) SDS-PAGE and Coomassie brilliant blue staining of gels loaded with recombinantly purified GST-tagged *L.p.* toxins (arrows) and GST. (b) Cells expressing epitope tagged *L.p.* toxins subjected to puromycin pulse-chase analyses. Immunoblotting of lysates was performed using antibodies against puromycin and Tubulin. Data are representative of 2 independent experiments. Values below the immunoblots represent densitometric quantification of puromycylated peptides normalized to the amount of Tubulin in cells. Expression levels of *L.p.* toxins were determined by qRT-PCR (Data represent mean ± standard error of the mean; n > 3 replicates per condition). Data were normalized to the average Ct values of the reference gene ribosomal protein S29 (*RPS29*). (c) Venn diagram depicting the common interacting partners enriched in GST-SidI pulldown eluates from HEK293T cell lysates (n = 2) and rabbit reticulocyte lysates (n = 1). (d) List of translation machinery components selectively enriched by GST-SidI across the three different experiments. GST-SidI enrichment is calculated as a ratio of the peptide intensities observed in GST-SidI pulldown eluates over GST pulldown eluates or marked as peptide intensities unqiuely measured only in GST-SidI pulldown eluates. *n.d.* – not detected.

**Figure S2.**
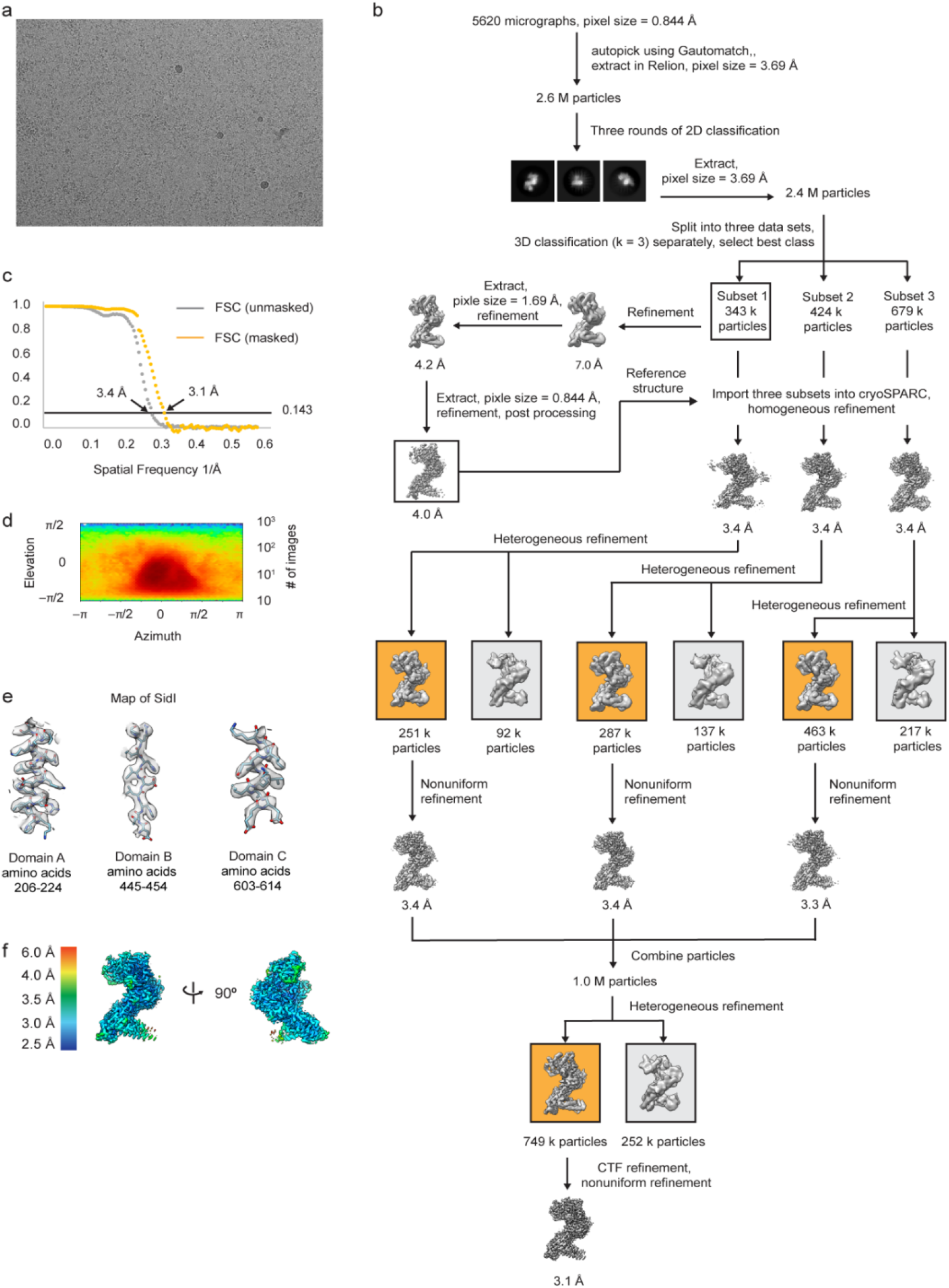
Cryo-EM workflow and characterization of SidI structure. (a) Representative micrograph showing the quality of data used for the final reconstruction of the SidI structure. (b) Data processing scheme of the SidI structure. (c) Fourier shell correlation (FSC) plots of the 3D reconstructions of SidI unmasked (grey), masked (orange). (d) Orientation angle distribution of the SidI reconstruction. (e) Electron microscopy maps of different regions of the SidI structure showing the quality of the data and the fit of the model. (f) Local resolution map of the SidI structure

**Figure S3.**
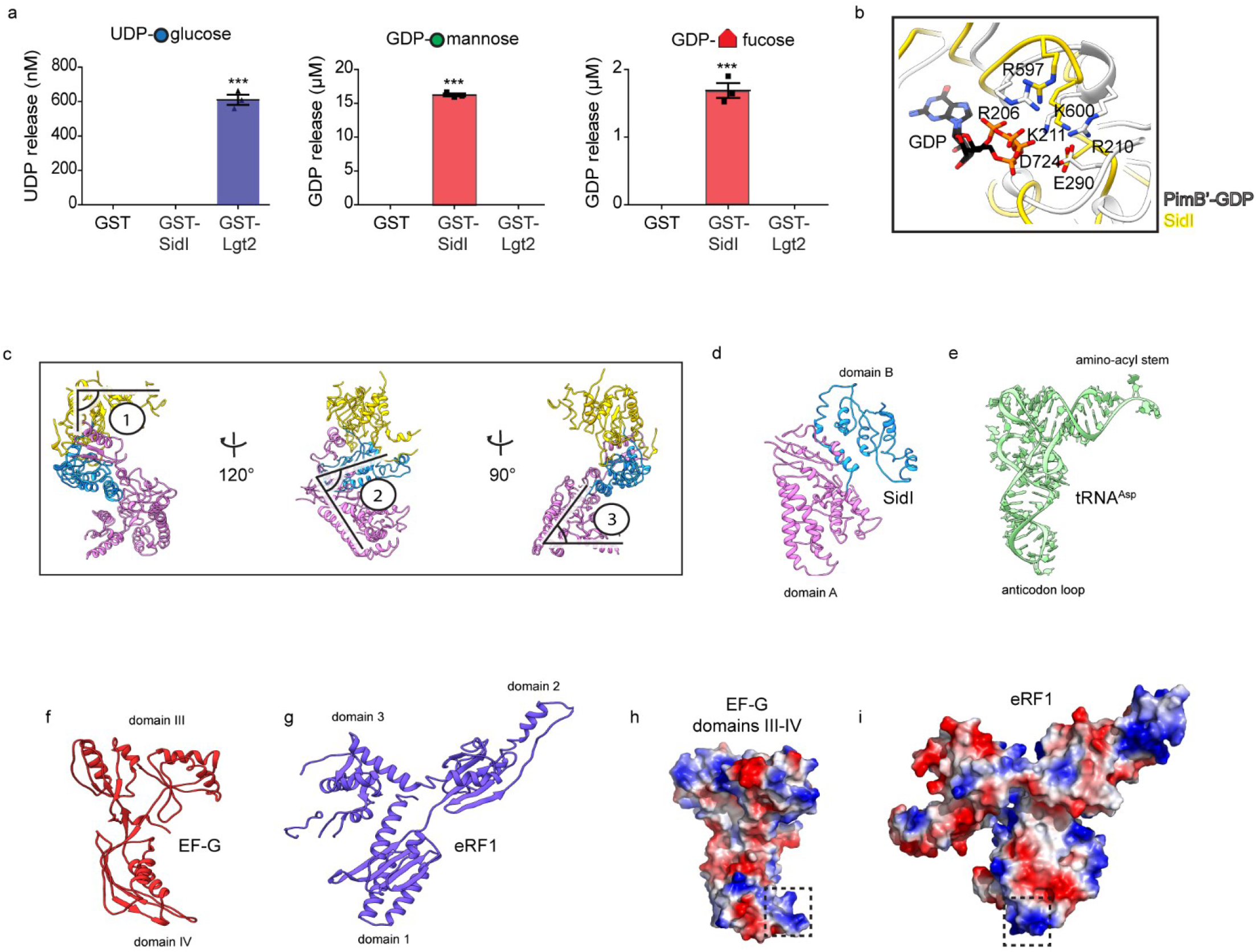
Characterization of SidI enzymatic function and tRNA-like shape. (a) GDP hydrolysis assay with 1µM of GST, GST-SidI and GST-Lgt2. 50µM of nucleotide-sugars were povided as precursors. Histograms depict amount of nucleotide release (Data represent mean ± standard error of the mean; n = 3 replicates per treatment condition). p-values were calculated by students t-test. ***p<0.001. (b) Overlay of SidI (gold) and GDP-PimB’ (light grey) showing the structural similarity between the enzymatic pockets and the conserved amino acids. (c) Different views of the SidI structure showing the three distinct kinks. (d – g) Comparison of the 3D architecture of the tRNA mimicry domains of SidI, EF-G and eRF1 to the structure of tRNA_asp_. (h – i) Surface charge distribution on the tRNA mimicry domains of EF-G and eRF1. Box highlights a positively charged loop region that contacts the ribosome.

**Figure S4.**
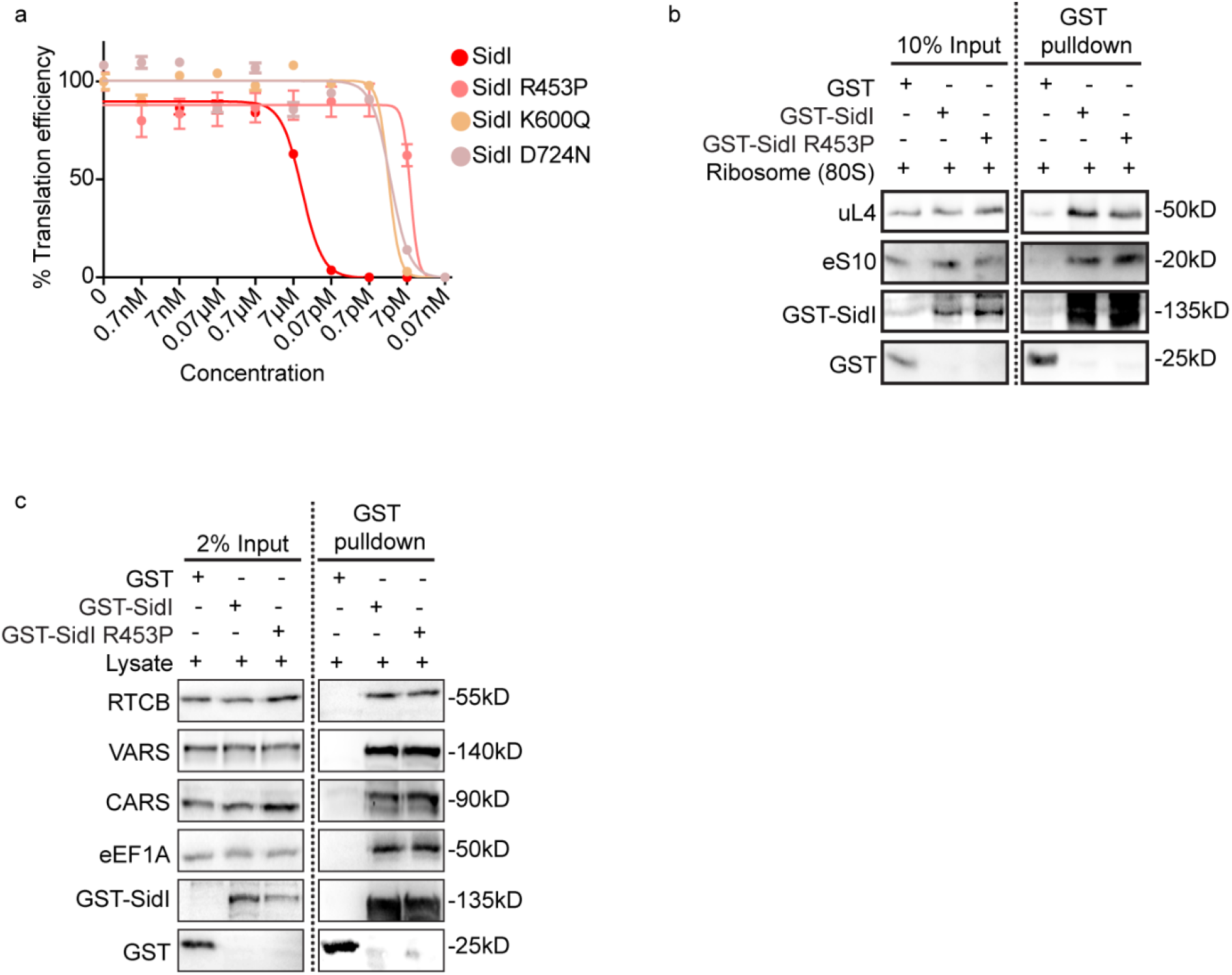
Enzymatic deficient mutants of SidI are attenuated in their ability to inhibit protein synthesis but still bind to the translation machinery. (a) Cell free translation of luciferase mRNA in RRLs incubated with purified GST-SidI, GST-SidI R453P; GST-SidI K600Q and GST-SidI D724N. Graph depicts the %translation efficiency of luciferase upon incubation with log-fold concentrations of proteins (Data represent mean ± standard error of the mean; n = 3 replicates per treatment condition). IC_50_ values were calculated by non-linear regression analysis. (b) Purified ribosome pulldown with GST, GST-SidI and GST-SidI R453P. Immunoblotting was performed using antibodies against 60S (uL4/RPL4) and 40S (eS10/RPS10) ribosomal proteins and GST. Data are representative of 2 independent experiments. (c) Immunoblot of tRNA interacting proteins VARS, CARS, RTCB and eEF1A precipiated by GST, GST-SidI or GST-SidI R453P from HEK293T lysates. Data are representative of 2 independent experiments.

**Figure S5.**
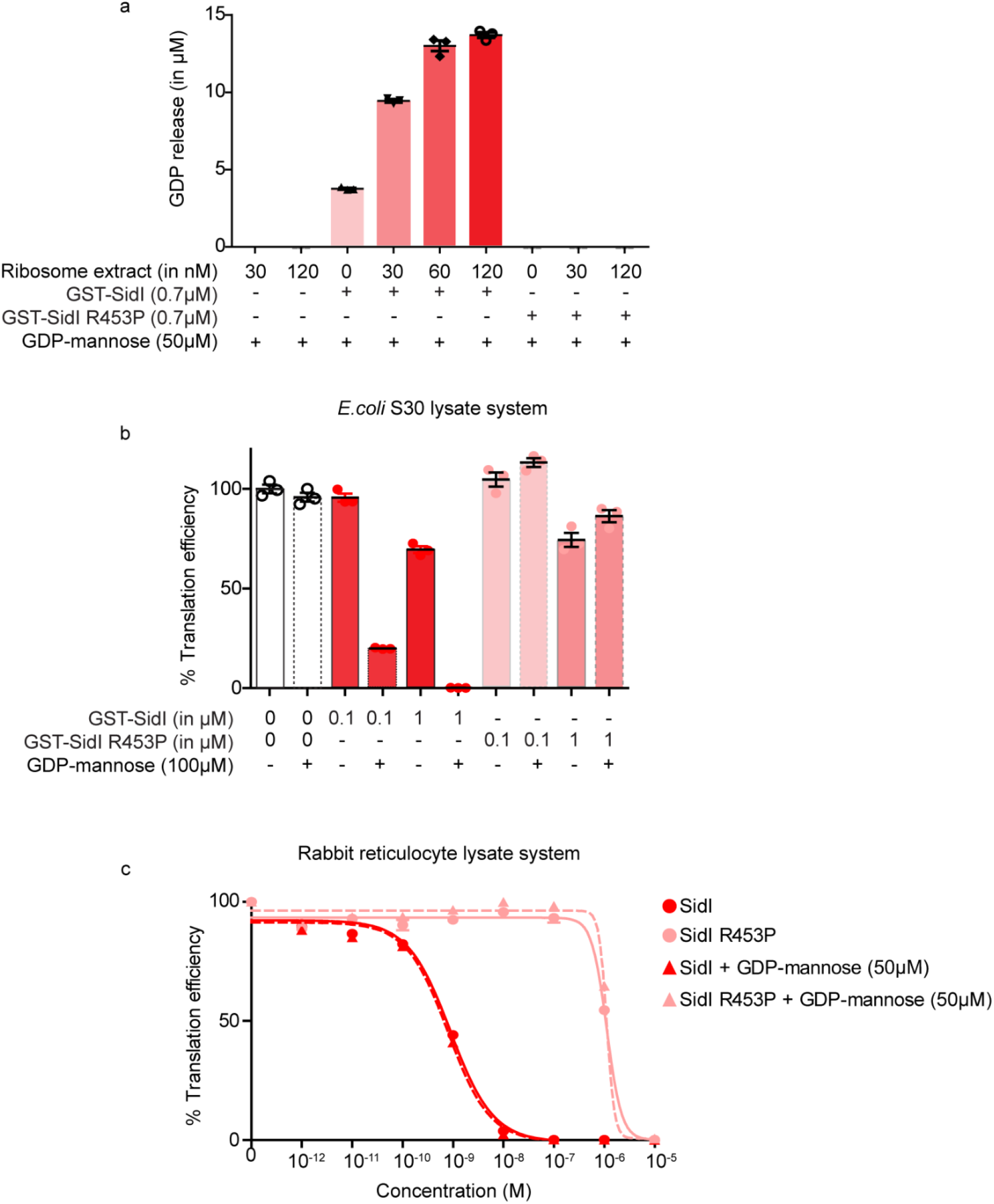
SidI inhibits protein synthesis in both prokaryotic and eukaryotic cell free extracts when GDP-mannose is present and by potentially targeting the ribosome. (a) GDP hydrolysis assay with GST-SidI or GST-SidI R453P incubated with increasing amounts of crude ribosome extracts purified from HEK293T cells in the presence of 50µM of GDP-mannose as precursor. Histograms depict amount of nucleotide release (Data represent mean ± standard error of the mean; n = 3 replicates per treatment condition); n.d. – not detected. (b)) Cell free transcription and translation of luciferase mRNA in *E.coli* S30 lysates incubated with GST-SidI or GST-SidI R453P in the absence or presence of 100µM GDP-mannose. Graph depicts the %translation efficiency of luciferase upon incubation with 0.1µM and 1µM of purified proteins (Data represent mean ± standard error of the mean; n = 3 replicates per treatment condition). (c) Cell free translation of luciferase mRNA in RRLs incubated with purified GST-SidI or GST-SidI R453P in the absence or presence of 50µM GDP-mannose. Graph depicts the %translation efficiency of luciferase upon incubation with log-fold concentrations of purified proteins (Data represent mean ± standard error of the mean; n = 3 replicates per treatment condition).

**Figure S6.**
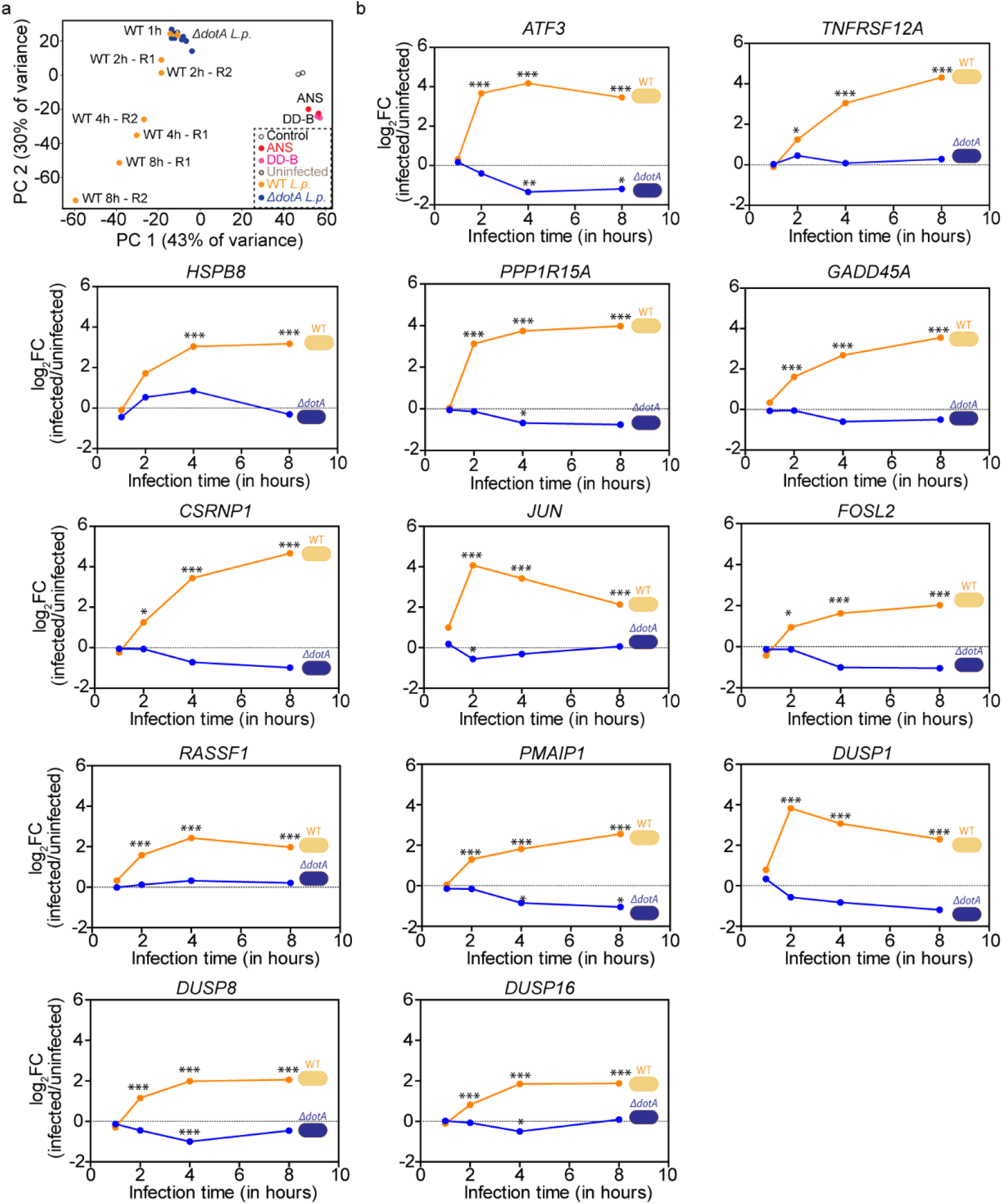
Analyses of RNA-seq datasets reveals a dynamic induction of stress transcripts during *L.p.* infection in an effector dependent manner. (a) Principal component analyses on all differentially expressed genes in HEK293 FcγR cells infected with WT and *ΔdotA L.p.* for 1 hour, 2 hours, 4 hours, and 8 hours and in HEK293T cells treated with ANS (0.1µM) and DD-B (0.1µM) for 4 hours. Coloured circles depict replicates. (b) Fold induction of upregulated transcripts in common with transcripts induced by ribosome stalling insults that regulate stress signaling and cell death. Expression levels after WT and *ΔdotA L.p.* infections normalized to uninfected controls. Data represent mean ± standard error of the mean; n = 2 replicates per time point. p-values were calculated by students t-test. *p<0.05; **p<0.01; ***p<0.001.

**Figure S7.**
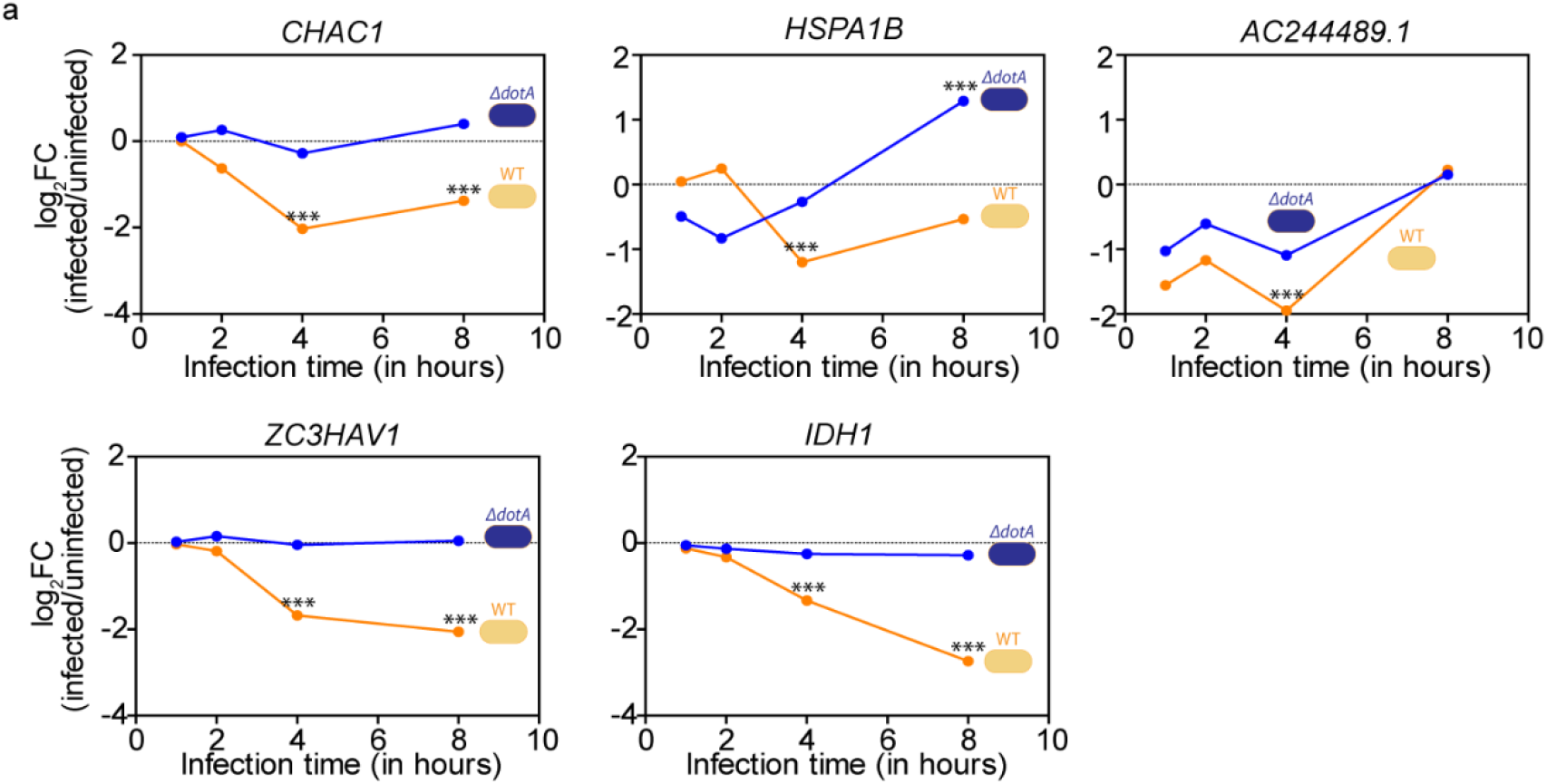
Analyses of downregulated transcripts during *L.p.* infection in common with ribosome stalling events. (a) Fold induction of downregulated transcripts upon WT and *ΔdotA L.p.* infection normalized to uninfected controls. Data represent mean ± standard error of the mean; n = 2 replicates per time point. p-values were calculated by students t-test. ***p<0.001.

**Figure S8.**
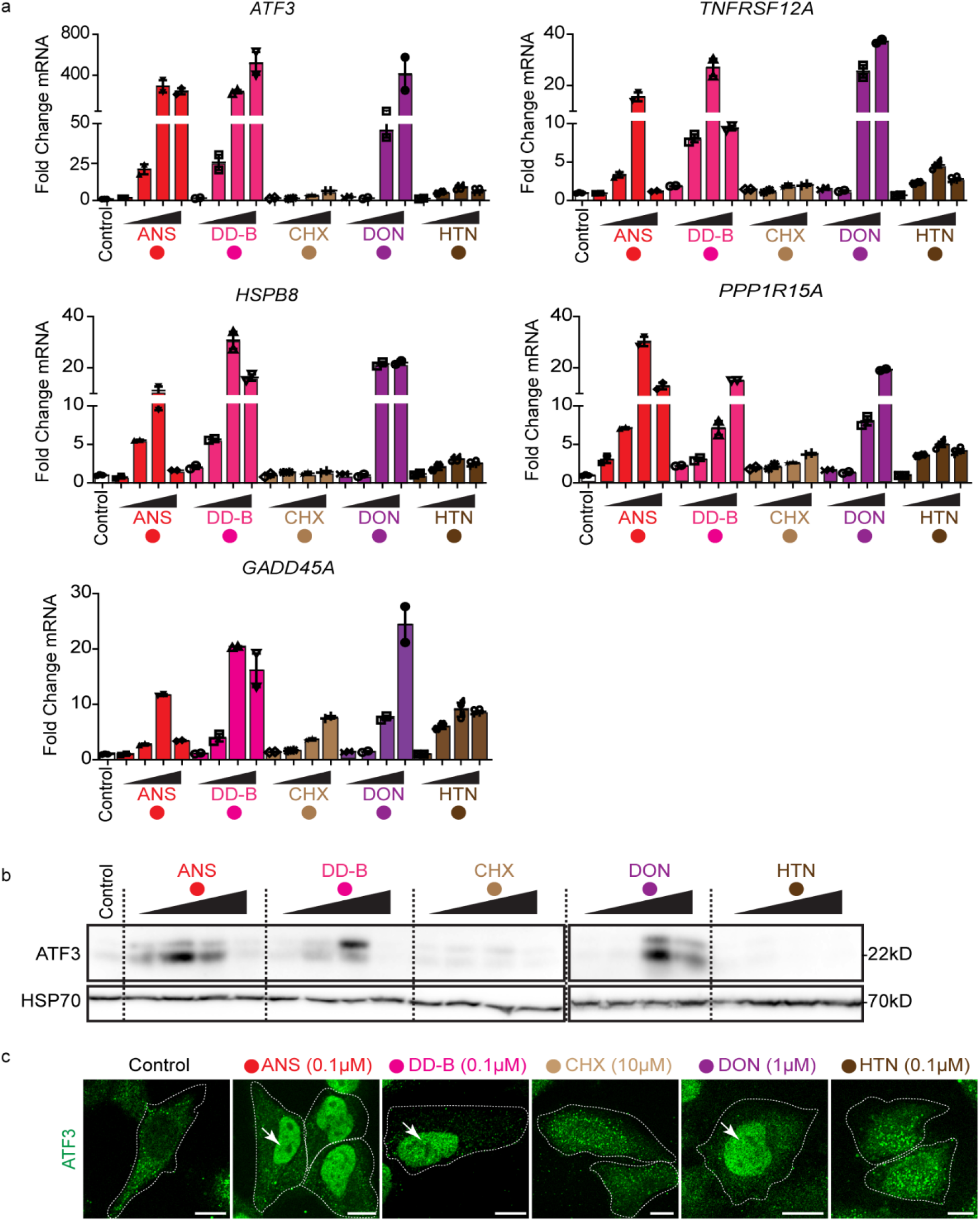
Specific inhibitors of ribosome function activate stress transcript induction and ATF3 accumulation in the nucleus. (a) Fold induction of *ATF3*, *TNFRSF12A*, *HSPB8*, *PPP1R15A* and *GADD45A* mRNA measured by qRT-PCR in HEK293T cells treated with ANS (0.01µM, 0.1µM, 1µM, 10µM); DD-B (0.001µM, 0.01µM, 0.1µM, 1µM); CHX (0.01µM, 0.1µM, 1µM, 10µM); DON (0.01µM, 0.1µM, 1µM, 10µM); and HTN (0.01µM, 0.1µM, 1µM, 10µM) for 4 hours (Data represent mean ± standard error of the mean; n = 2 replicates per condition). Transcript expression values were normalized internally to the reference gene *RPS29* and expressed as fold change over the levels in vehicle treated controls. (b) Immunoblotting of ATF3 in lysates from HEK293T cells treated with ANS (0.01µM, 0.1µM, 1µM, 10µM); DD-B (0.001µM, 0.01µM, 0.1µM, 1µM); CHX (0.01µM, 0.1µM, 1µM, 10µM); DON (0.01µM, 0.1µM, 1µM, 10µM); and HTN (0.01µM, 0.1µM, 1µM, 10µM) for 4 hours. Data are representative of 3 independent experiments. (c) Indirect immunofluorescence of ATF3 in HeLa cells treated with ANS, DD-B, CHX, DON and HTN at the indicated concentrations. Scale bar = 10µM. Data are representative of 2 independent experiments. Nuclear region is marked by arrows.

**Figure S9.**
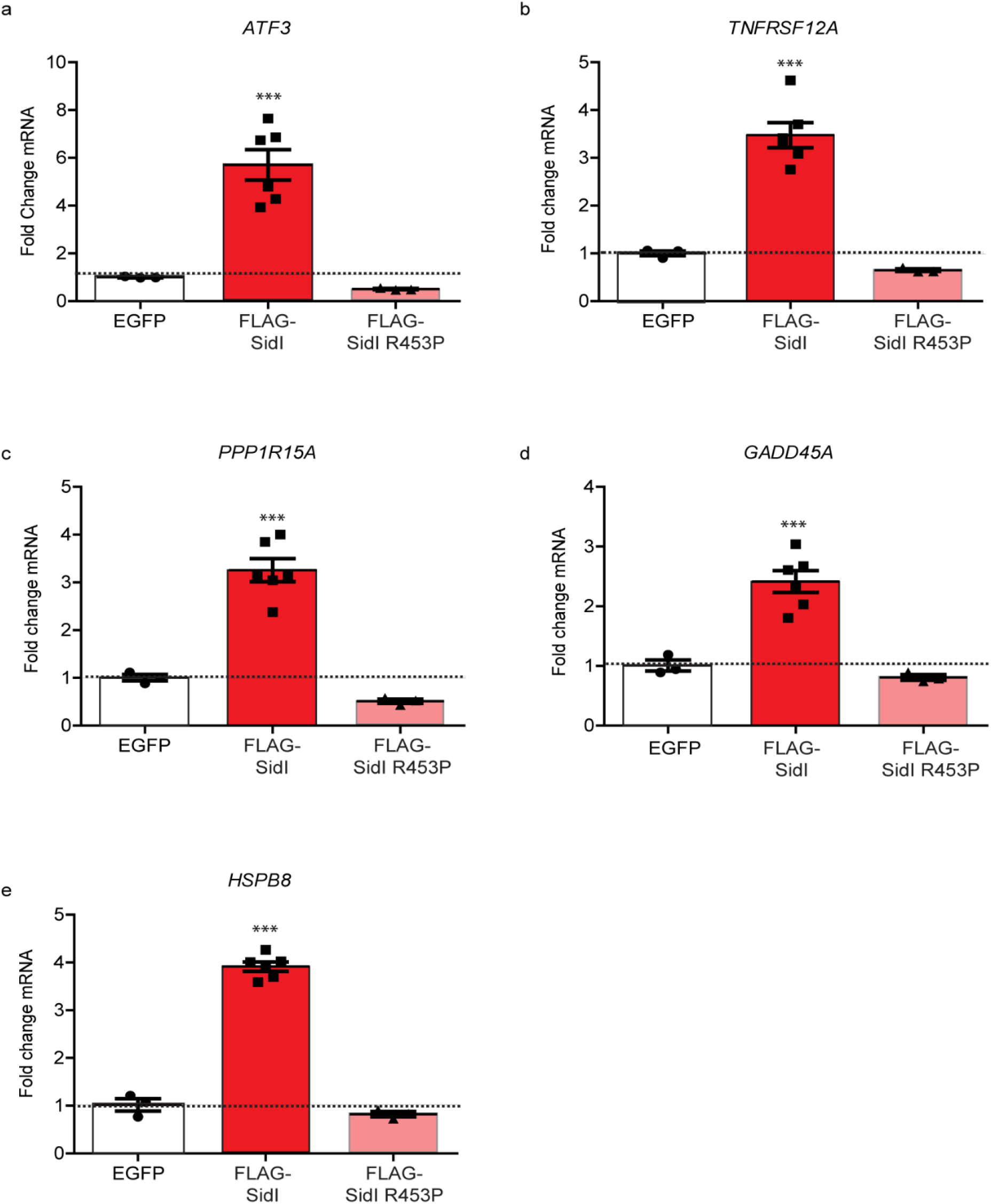
Glycosyl transferase activity of SidI is essential for stress transcript induction. (a – e) Fold induction of stress transcript mRNAs measured by qRT-PCR in HEK293T cells expressing GFP, FLAG-SidI or FLAG-SidI R453P (Data represent mean ± standard error of the mean; n ≥ 3 independent replicates). Transcript expression values were normalized internally to the reference gene *RPS29* and expressed as fold change over the levels in GFP expressing cells.

**Figure S10.**
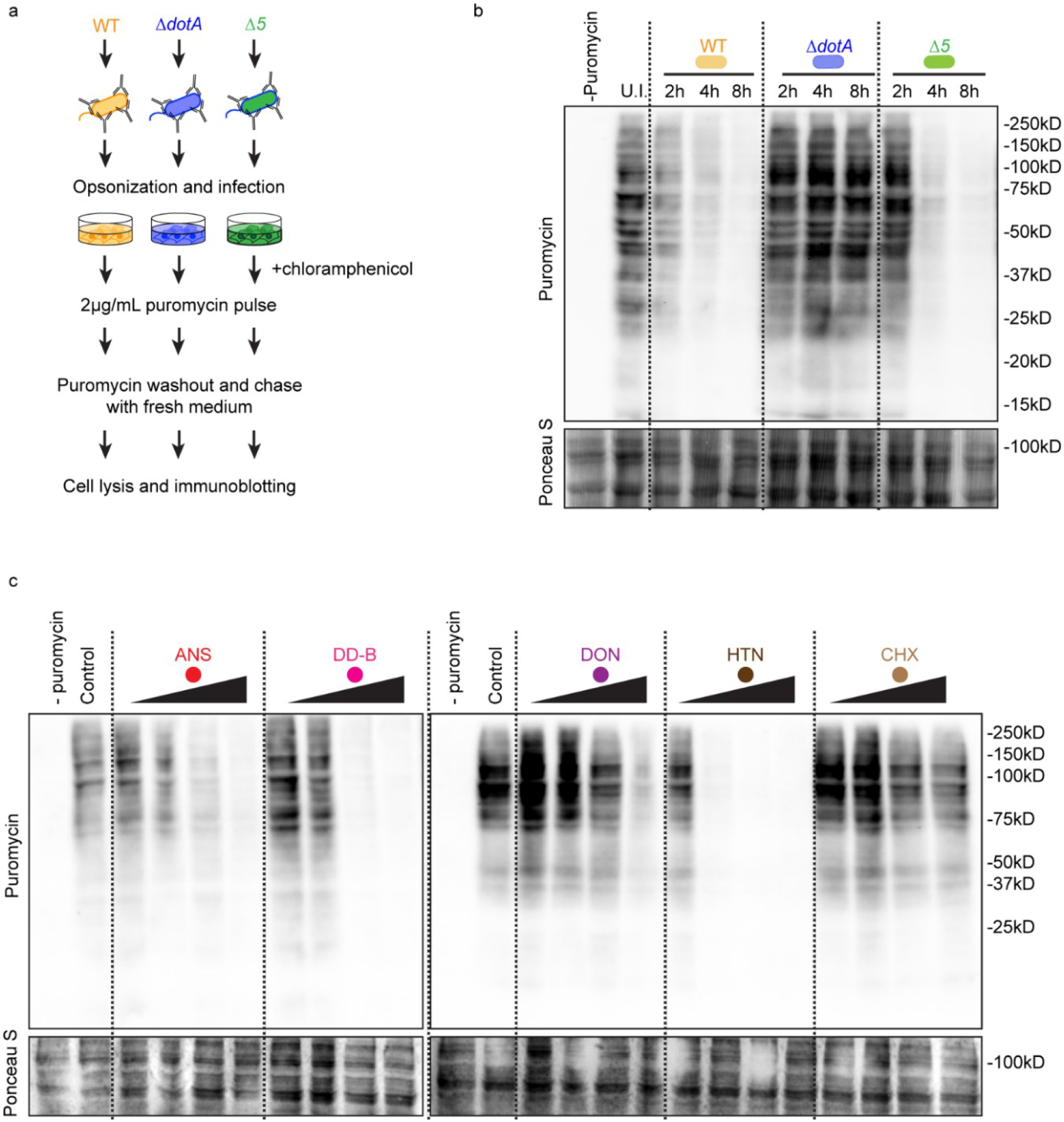
*L.p.* infection and elongation inhibitor treatment conditions inhibit protein synthesis. (a) Schematic of the experimental setup to interrogate nascent protein synthesis rates in HEK293 FcγR cells infected with WT, *ΔdotA* and *Δ5 L.p.* strains. (b) Immunoblotting of lysates from cells infected with WT, *ΔdotA* and *Δ5 L.p.* was performed using antibodies against puromycin. Ponceau S staining of membranes serves as a loading control. Data are representative of 2 independent experiments. (c) Immunoblotting of lysates from cells treated with ANS (0.01µM, 0.1µM, 1µM, 10µM); DD-B (0.001µM, 0.01µM, 0.1µM, 1µM); CHX (0.01µM, 0.1µM, 1µM, 10µM); DON (0.01µM, 0.1µM, 1µM, 10µM); and HTN (0.01µM, 0.1µM, 1µM, 10µM) for 4 hours was performed using antibodies against puromycin. Ponceau S staining of membranes serves as a loading control. Data are representative of 2 independent experiments.

**Figure S11.**
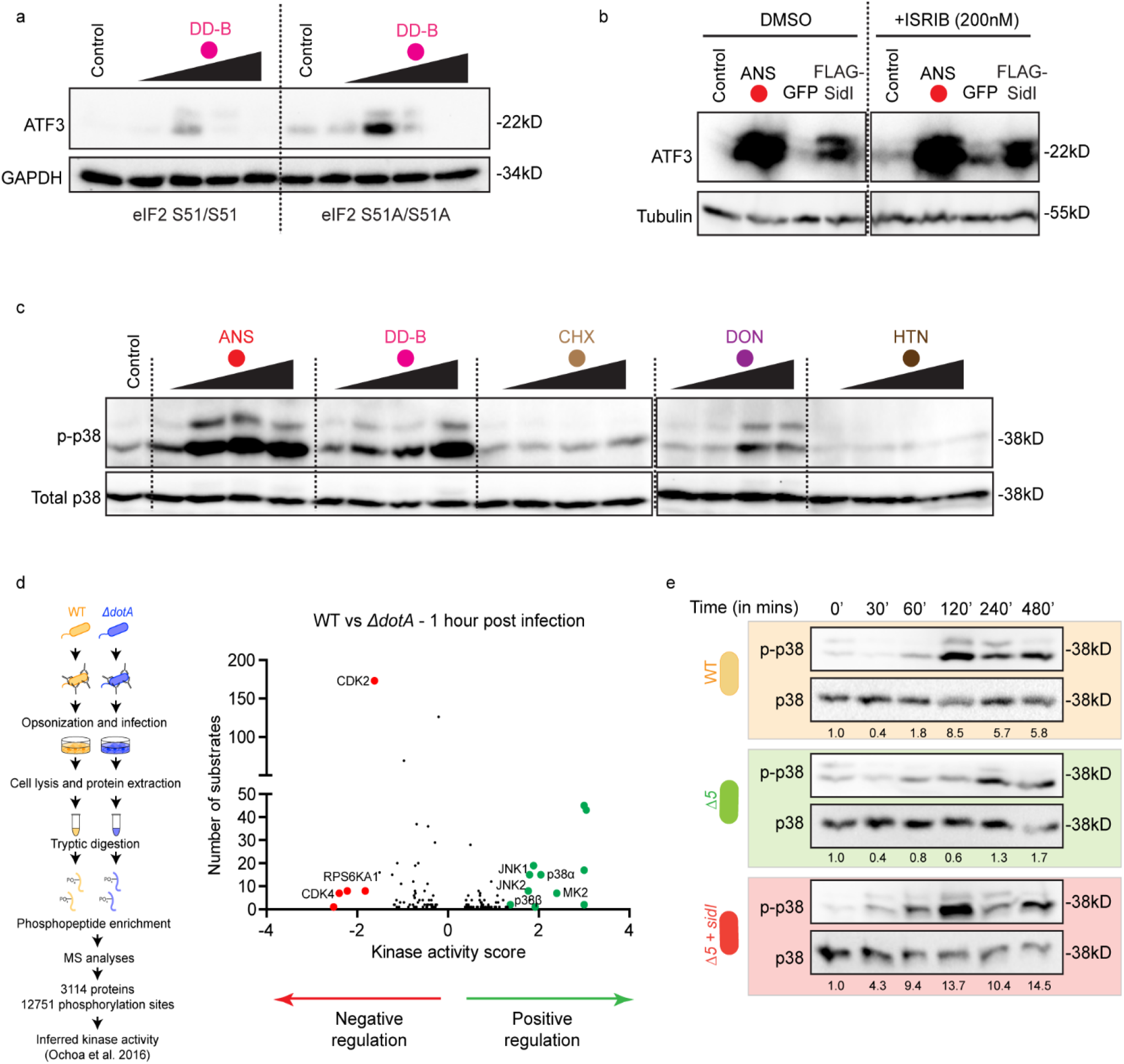
Ribosome stalling events activate p38 phosphorylation and do not require an active ISR for ATF3 accumulation. Immunoblotting of ATF3 in lysates from (a) WT or eIF2 *S51A^+/+^* mouse embryonic fibroblast cells treated with DD-B (0.001µM, 0.01µM, 0.1µM and 1µM) for 4 hours; and (b) HEK293T cells treated with ANS (0.1µM) or transfected with FLAG-SidI in the absence or presence of ISRIB (200nM). GAPDH (a) and Tubulin (b) serve as loading controls. (c) Immunoblotting of phospho-p38 and total p38 in lysates of HEK293T cells treated with ANS (0.01µM, 0.1µM, 1µM, 10µM); DD-B (0.001µM, 0.01µM, 0.1µM, 1µM); CHX (0.01µM, 0.1µM, 1µM, 10µM); DON (0.01µM, 0.1µM, 1µM, 10µM); and HTN (0.01µM, 0.1µM, 1µM, 10µM) for 4 hours. (d) Phosphoproteomics pipeline and inferred kinase activation in WT *L.p.* infected cells (MOI = 100) based on the analyses of phosphorylation sites on proteins. Kinases of the stress activated protein kinase pathway are positively regulated and highlighted in green. (e) Immunoblotting of phospho-p38 and total p38 in lysates from HEK293 FcγR cells infected with WT, *ΔdotA* and *Δ5 L.p.* strains (MOI = 50). Values below immunoblots represent densitometric quantification of phospho-p38 levels normalized to total p38 levels.

**Figure S12.**
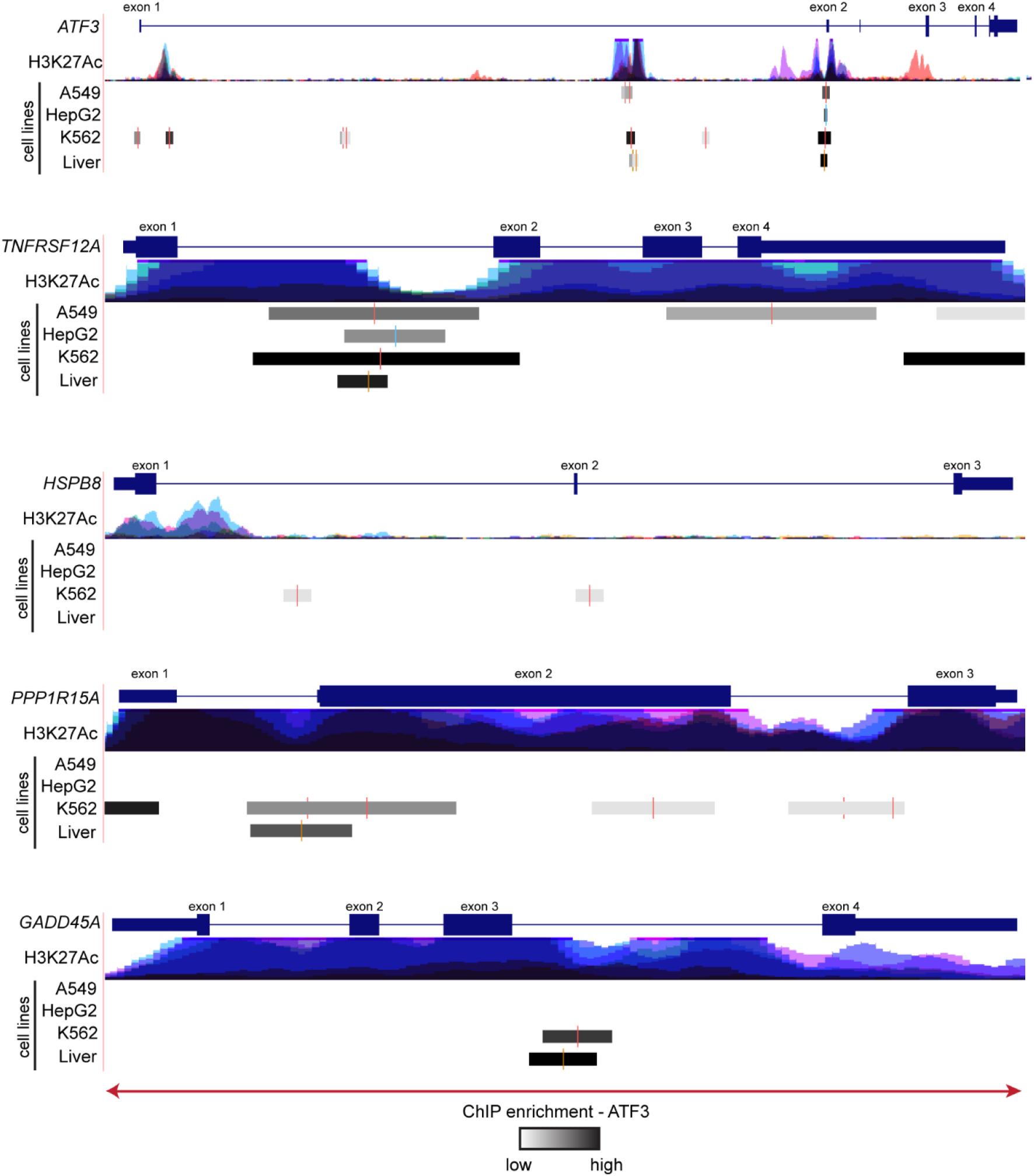
ATF3 binds to genomic regions of stress inducible transcripts. Mapping and enrichment of ATF3 binding zones on the *ATF3*, *TNFRSF12A*, *HSPB8*, *PPP1R15A* and *GADD45A* genes across four cell types. H3K27Ac tracks mark acetylation of lysine 27 of the H3 histone protein across 7 cell types (GM12878, H1-hESC, HSMM, HSMM, K562, NHEK and NHLF cells). Tracks of each cell type are marked with a different colour and displayed as a transparent overlay. The enrichment scores for ATF3 peaks were extracted from the ENCODE project datasets and visualized using UCSC genome browser (human hg38 assembly). The display for this track shows site location with the point-source of the peak marked with a coloured vertical bar and the level of enrichment at the site indicated by the darkness of the item. The enrichment values were computed based on signal values assigned by the ENCODE pipeline. The input signal values were multiplied by a normalization factor calculated as the ratio of the maximum score value (1000) to the signal value at 1 standard deviation from the mean, with values exceeding 1000 capped at 1000.

**Figure S13.**
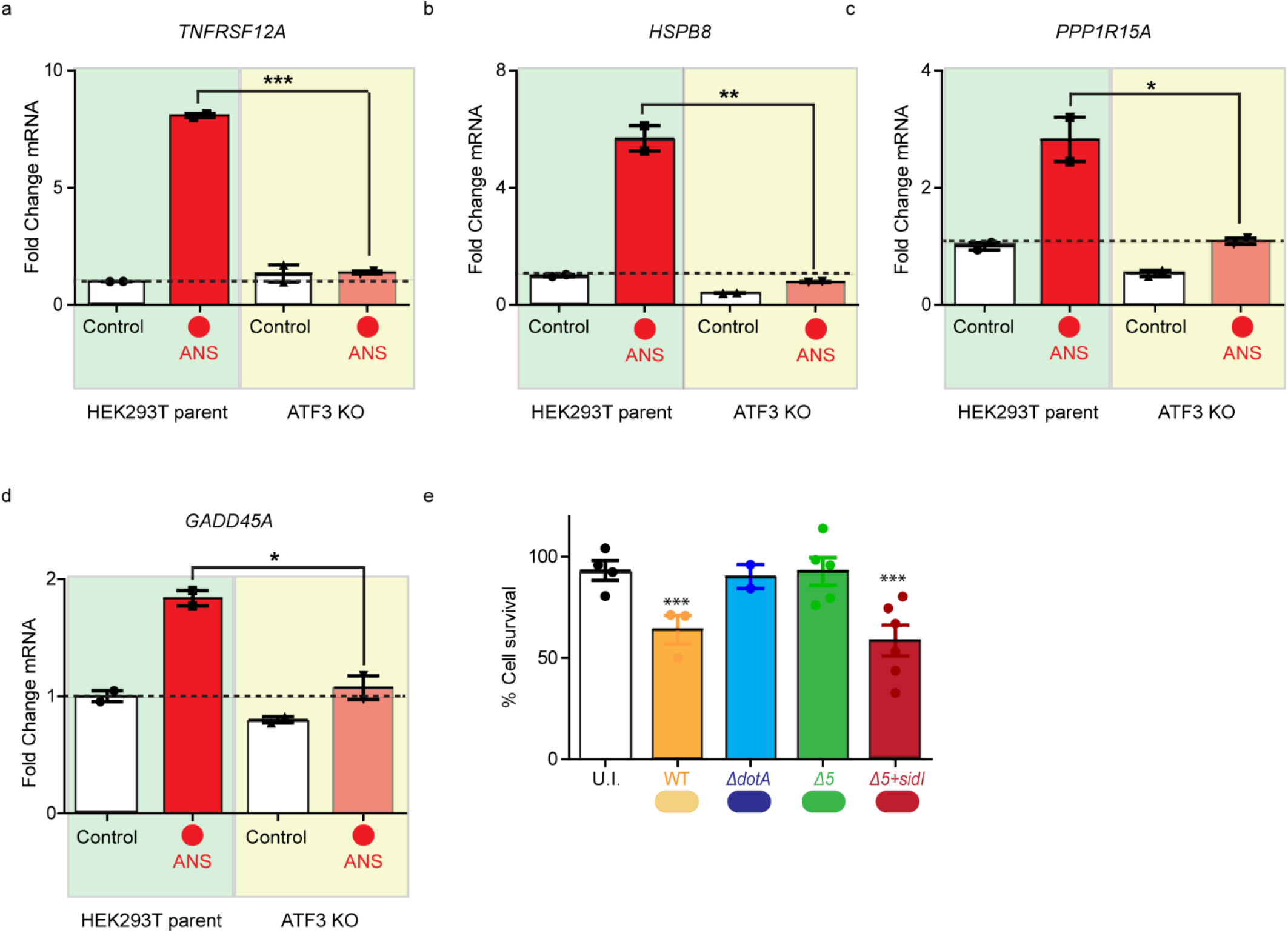
ATF3 accumulation is necessary for stress transcript induction and activation of cell death. (a – d) Fold change induction of *TNFRSF12A*, *HSPB8*, *PPP1R15A* and *GADD45A* mRNA in parent and ATF3 KO HEK293T cells treated with ANS (0.1µM) for 4 hours and measured by qRT-PCR (Data represent mean ± standard error of the mean; n = 2 independent experiments). Transcript expression values were normalized internally to the reference gene *RPS29* and expressed as fold change over the levels in vehicle treated controls. p values calculated by students t-test. *p<0.05; **p<0.01; ***p<0.001. (e) Cell viability measurements in HEK293 FcγR cells infected with WT, *ΔdotA*, *Δ5 or Δ5* + *sidI L.p.* strains (MOI = 50) for 10 hours (Data represent mean ± standard error of the mean; n ≥ 3 independent replicates per condition). p-values were calculated by students t-test. ***p<0.001.

**Figure S14.**
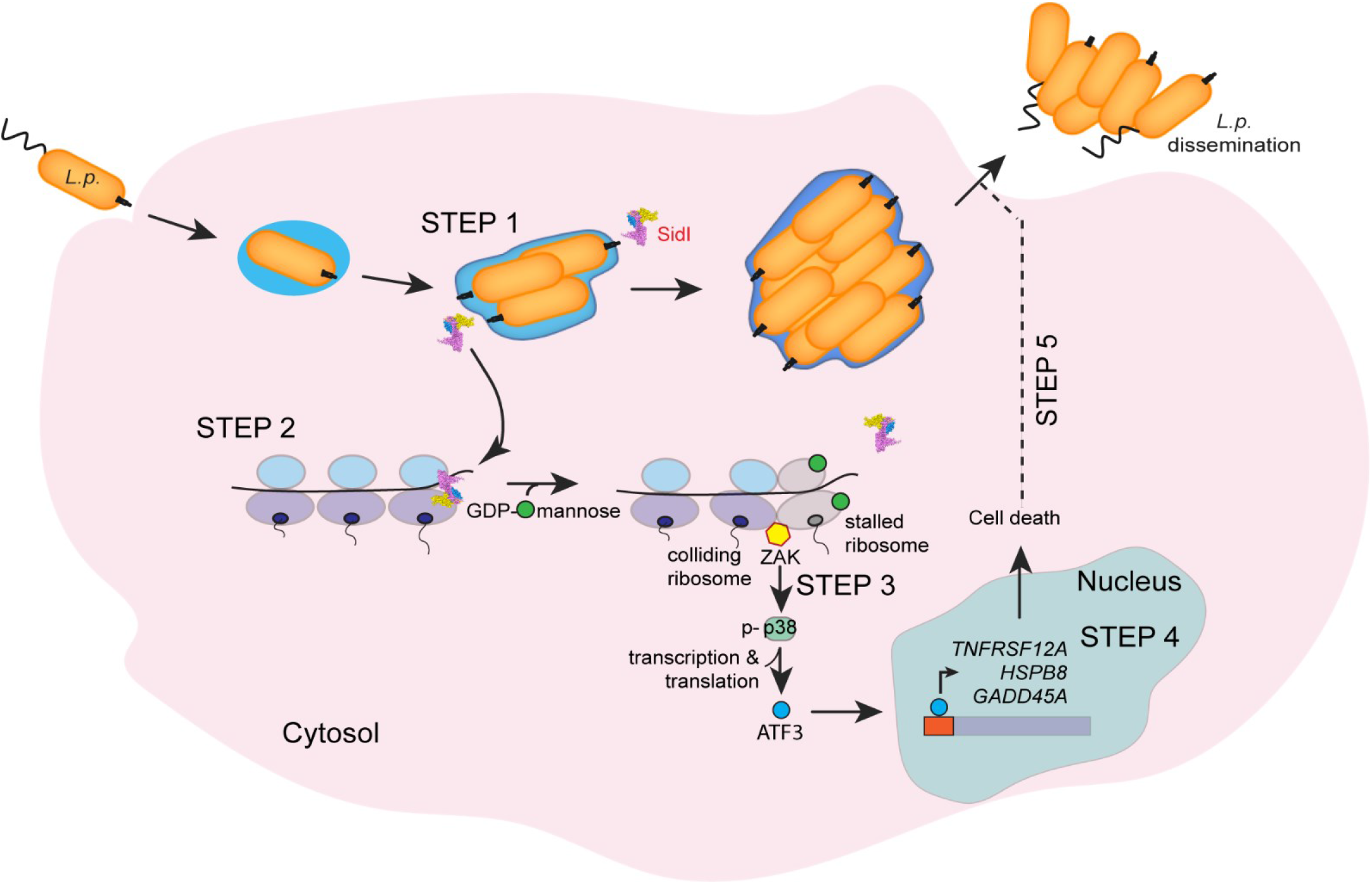
*Legionella* activates a ribosome to nuclear signaling pathway. A five step schematic model of how *L.p.* hijacks a stress response pathway. **Step 1** – *L.p.* infects the host cell, establishes its replicative niche and secretes SidI into the host cell cytosol. **Step 2** – SidI by way of its tRNA mimicry domain is targeted specifically to components of the host translation machinery, including the ribosomes depicted in the figure. In the presence of cytosolic GDP-mannose, SidI mannosylates the ribosome and causes translation elongation defects that result in an accumulation of stalled and collided ribosomes in cells. **Step 3 –** Collided ribosomes are sensed as ‘aberrant’, resulting in the recruitment and activation of the kinase ZAKα that in turn phosphorylates and activates p38. This stress pathway activation results in the accumulation and translocation of ATF3 from a diffuse cytosolic disposition to the nucleus. **Step 4 –** ATF3 binds to genomic regions of stress inducible transcripts and regulates the expression. **Step 5 –** ATF3, via the transcriptional program it orchestrates, induces host cell lysis, thereby resulting in the dissemination of replicated bacteria into the extracellular milieu.

