## Supplemental table S2 for "A *Legionella* toxin mimics tRNA and glycosylates the translation machinery to trigger a ribotoxic stress response"

| Structure | SidI (PDB ID: XXXX) |
| --- | --- |
| **Data collection** | |
| Microscope | Titan Krios |
| Voltage (keV) | 300 |
| Nominal magnification | 81000x |
| Exposure navigation | Image shift |
| Electron dose (e^-^Å^-2^) | 60 |
| Dose rate (e^-^/pixel/sec) | 7.5 |
| Detector | K3 summit |
| Pixel size (Å) | 0.844 |
| Defocus range (μm) | 1.0-2.0 |
| Micrographs | 5620 |
| **Reconstruction** | |
| Total extracted particles (no.) | 2622904 |
| Final particles (no.) | 748767 |
| Symmetry imposed | C1 |
| FSC average resolution, masked (Å) | 3.1 |
| FSC average resolution, unmasked (Å) | 3.4 |
| Applied B-factor (Å) | 187.1 |
| Reconstruction package | Relion 3.0.8 Cryosparc v3.3.1 |
| **Refinement (numbers in this section will change when model is complete)** | |
| Protein residues | 777 |
| Ligands | 0 |
| RMSD Bond lengths (Å) | 0.002 |
| RMSD Bond angles (^o^) | 0.620 |
| Ramachandran outliers (%) | 0.00 |
| Ramachandran allowed (%) | 8.56 |
| Ramachandran favored (%) | 91.44 |
| Poor rotamers (%) | 3.30 |
| CaBLAM outliers (%) | 5.13 |
| Molprobity score | 2.40 |
| Clash score (all atoms) | 9.40 |
| B-factors (protein) | 42.01 |
| B-factors (ligands) | N/A |
| EMRinger Score | 1.94 |
| Refinement package | Phenix 1.17.1-3660-000 |
