## Supplemental table S6 for "A *Legionella* toxin mimics tRNA and glycosylates the translation machinery to trigger a ribotoxic stress response"

**Table S6. List of sgRNAs, siRNAs and primers used in this study.**

**sgRNAs**

| **Gene target** | **sgRNA sequence** |
| --- | --- |
| ATF3 guide #1 | U*C*C*GAGGCAGAGACCUGGCC + Synthego modified EZ Scaffold |
| ATF3 guide #2 | C*C*U*UUGUCAAGGAAGAGCUG + Synthego modified EZ Scaffold |
| ATF3 guide #3 | C*C*A*GCGCAGAGGACAUCCGG + Synthego modified EZ Scaffold |

**siRNAs**

| **Gene target** | **siRNA/Commercial product** |
| --- | --- |
| Control siRNA | Mission siRNA Universal Negative control (Sigma-Aldrich Cat #SIC001) |
| ZAK (MAP3K20) | #1 – 5’ – CTATGATTACATTAACAGT – 3’  #2 – 5’ – GGATCACTCTATGATTACA – 3’  #3 – 5’ – CCTCAAGTCAAGAAACGTT – 3’ |
| p38 (MAPK14) | #1 – 5’ – CCTAAAACCTAGTAATCTA – 3’  #2 – 5’ – GAAGCTCTCCAGACCATTT – 3’  #3 – 5’ – CTCCGAGGTCTAAAGTATA – 3’ |
| ATF3 | #1 – 5’ – GGAGGACTCCAGAAGATGA – 3’  #2 – 5’ – CTGGGTCACTGGTGTTTGA – 3’  #3 – 5’ – CACGTGCAGTATCTCAAGA – 3’ |

**Primers (5’🡪3’)**

| **Gene target** | **Primers** |
| --- | --- |
| *lgt1* | Forward Primer ATACGGTCGCCCGATATTTG  Reverse Primer TCCTTTCCTCTTGGCACTTATT |
| *lgt2* | Forward Primer GGAATTTGGCACAGGCCGAGT  Reverse Primer GCACCAGATCAGAGGCAGCA |
| *lgt3* | Forward Primer TCTGATCTGGTGCGTTGGG  Reverse Primer AGCAGGCTGTTGACCATCCTTT |
| *sidI* | Forward Primer TGGTACGATTCGTCCTGGCA  Reverse Primer TGTGGGTCACCTGCCAGTTT |
| *sidL* | Forward Primer GCGCATAAAGGGCGCGATAG  Reverse Primer ACTCCACCGCCACAATTCCA |
| *legK4* | Forward Primer GCCATCAGGTGAAACAATTCC  Reverse Primer CTGCTGCCGAAATCAACTAAAT |
| *ravX* | Forward Primer GGGTTAGTCCTTTCGATACACTATT  Reverse Primer CGCAGGTACAGAGCAATGATA |
| *ATF3* | Forward Primer CCTCTGCGCTGGAATCAGTC  Reverse Primer TTCTTTCTCGTCGCCTCTTTTT |
| *TNFRSF12A* | Forward Primer GACCGCACAGCGACTTCT  Reverse Primer CACGAAGGTCAGGCTCAGA |
| *PPP1R15A* | Forward Primer ATGATGGCATGTATGGTGAGC  Reverse Primer AACCTTGCAGTGTCCTTATCAG |
| *HSPB8* | Forward Primer CTCCTGCCACTACCCAAGC  Reverse Primer GGCCAAGAGGCTGTCAAGT |
| *GADD45A* | Forward Primer GAGAGCAGAAGACCGAAAGGA  Reverse Primer CACAACACCACGTTATCGGG |
| *RPS29* | Forward Primer CGCTCTTGTCGTGTCTGTTCA  Reverse Primer CCTTCGCGTACTGACGGAAA |
| SidI R453P mutagenesis | Forward Primer TATTCATATTcctCCACCAGATTGC  Reverse Primer TCAAGATGAGTATCCTCATC |
| SidI K600Q mutagenesis | Forward Primer TCGTCCTGGCcAAGGGTTTGA  Reverse Primer ATCGTACCAAAAGATAAGATATTAGG |
| SidI D724N mutagenesis | Forward Primer TTGTCGAATGaATGACATGGG  Reverse Primer ACGTATTTACAGTTTTGC |
